# A Substrate-Responsive Catalytic Gate for Peptide Ligation

**DOI:** 10.64898/2026.11.29.769881

**Authors:** Shi Qi, Wenyu Du, Jiahui Ma, Yufan Guo, Jiarong Mo, Xinya Hemu

## Abstract

How enzymes achieve selectivity between competing reaction pathways that share a common catalytic intermediate remains a fundamental question in understanding their functional diversification. In the legumain family, from which peptide asparaginyl ligases (PALs) have naturally diverged, sequence features such as ligase-activity determinants (LADs) have proven useful for identifying and engineering ligases but these empirical rules do not explain how a highly conserved catalytic center governs the competition between aminolysis and hydrolysis. Here, combining a sliding-window ensemble classifier with protein-language-model screening, we mapped the sequence-function landscape of a hydrolase EchAEP1, independently identified a ligation-activity peak centered on LAD1 but beyond the conventional LAD composition. Remarkably, a single G243Q substitution increased the intermolecular ligation-to-hydrolysis ratio approximately 59-fold primarily through a pronounced suppression of hydrolysis, yielding a contrasting enzyme pair on a near-identical sequence background. Despite similar active-site architectures observed in crystal structures, molecular dynamics simulations revealed distinct conformational behavior of the thioester acyl-enzyme intermediate of G243Q, as the basal oxyanion-hole occupancy was suppressed without excluding attacking water, whereas favorable peptide-attack geometries were associated with restoration of oxyanion-hole occupancy to near-wild-type levels. Notably, the direction of this peptide-associated response varied across enzyme backgrounds, while reduced basal oxyanion-hole occupancy consistently correlated with increased relative ligation selectivity across different enzymes and pH conditions. Together, we propose a context-dependent nucleophile-gated oxyanion-hole model, which suggest that catalytic conformational competence, rather than nucleophile accessibility alone, is a critical dimension of reaction-pathway selectivity, providing a mechanistic framework for understanding how enzyme sequence and substrate geometry jointly shape catalytic outcomes.

## Introduction

Peptide ligases selectively catalyze peptide-bond formation, enabling topological reshaping and terminal modification of peptide chains, and provide important biocatalytic tools for the synthesis and screening of cyclic peptide drugs, precise protein modification, and bioconjugation^1^. Among these enzymes, peptide asparaginyl ligases (PALs) are distinguished by their high catalytic efficiency and broad substrate tolerance^2–4^. PALs and the widely distributed hydrolytic enzymes asparaginyl endopeptidases (AEPs) belong to the same legumain family of cysteine proteases but exhibit opposite catalytic functions: ligation and hydrolysis, respectively^5,6^. Despite their different functional preferences, both share highly conserved active-site architectures and proceed through a common thioester acyl-enzyme intermediate, formed by nucleophilic attack of the catalytic cysteine on the backbone carbonyl of a substrate Asn or Asp residue. During deacylation, amine nucleophiles compete with water for attack on this intermediate, yielding ligation and hydrolysis products, respectively. For PALs, these nucleophiles include N-terminal peptide amino groups, Lys side-chain amines^7^, and non-peptide amines^8^, enabling backbone cyclization and intermolecular ligation. The ability of PALs to favor ligation over hydrolysis in aqueous environments makes them an intriguing model for understanding how conserved catalytic machinery controls selectivity between competing reaction pathways.

To explain the functional divergence of these two enzyme classes, it is necessary to understand the factors that partition the common catalytic intermediate between the competing aminolysis and hydrolysis pathways. Previous structural and functional studies have identified several regions influencing the catalytic performance of plant legumains. Structure-guided engineering of OaAEP1 identified a gatekeeper residue in the S2 pocket that modulate catalytic efficiency, leading to the highly active C247A variant^9^. Subsequent cross-species comparisons and functional studies led to the proposal of a marker of ligase activity (MLA), showing that a loop adjacent to the substrate-binding region can act together with active-pocket residues to influence cyclization efficiency^6^. However, these studies primarily addressed catalytic efficiency rather than the factors governing ligation versus hydrolysis. As the collection of natural PALs expanded, we proposed the ligase-activity determinants (LADs) hypothesis based on studies of homologous enzymes from *Viola*^5^. This hypothesis linked LAD1 site in S2 pocket and LAD2 site in the prime-side substrate-binding region to ligation preference. It successfully guided the engineering of ligation activity into both natural AEP^10^ and consensus-designed hydrolases^11^, demonstrating the role of local substrate-binding regions in regulating enzyme function.

As works for LAD-based engineering accumulated, the limits of LAD-based control over reaction selectivity have become apparent in both directions. Installing LAD-positive features in AEPs could raise ligation/hydrolysis (L/H) ratio significantly, but does not fully abolish hydrolysis^10,11^. Conversely, reciprocal substitution of LAD residues in natural ligases lower L/H ratio but does not yield efficient hydrolases^5^. These asymmetric outcomes suggest that LAD residues are important modulators of reaction selectivity, but their effects depend on the broader enzyme sequence context. How this sequence context influences the competition between aminolysis and hydrolysis remains unclear.

One conventional explanation lies in how these residues influence the active-site environment, particularly water accessibility. Any enzyme that forms a peptide bond in water must suppress attack by a vast excess of solvent, and in other peptide-macrocyclizing families this is achieved structurally, by sequestering the catalytic center from bulk solvent. For instance, the PatGmac macrocyclization insertion partly occludes the active site^12^, PCY1 stabilizes a closed conformation on binding its recognition peptide^13^, the Sortase A surface groove is delineated by flexible loops that adjust upon substrate binding^14^, and a lipocalin-like domain contributes to the SurE substrate cleft^15^ (**Figure 1a**). The mature PAL core domain, by contrast, is comparatively open and flat and lacks any equivalent occluding element^6,16^, so its preference for aminolysis in a water-rich environment cannot be explained by occlusion of the active site or exclusion of water. If the two nucleophiles are not separated in space, they must be distinguished at the catalytic center itself.

**Figure 1.**
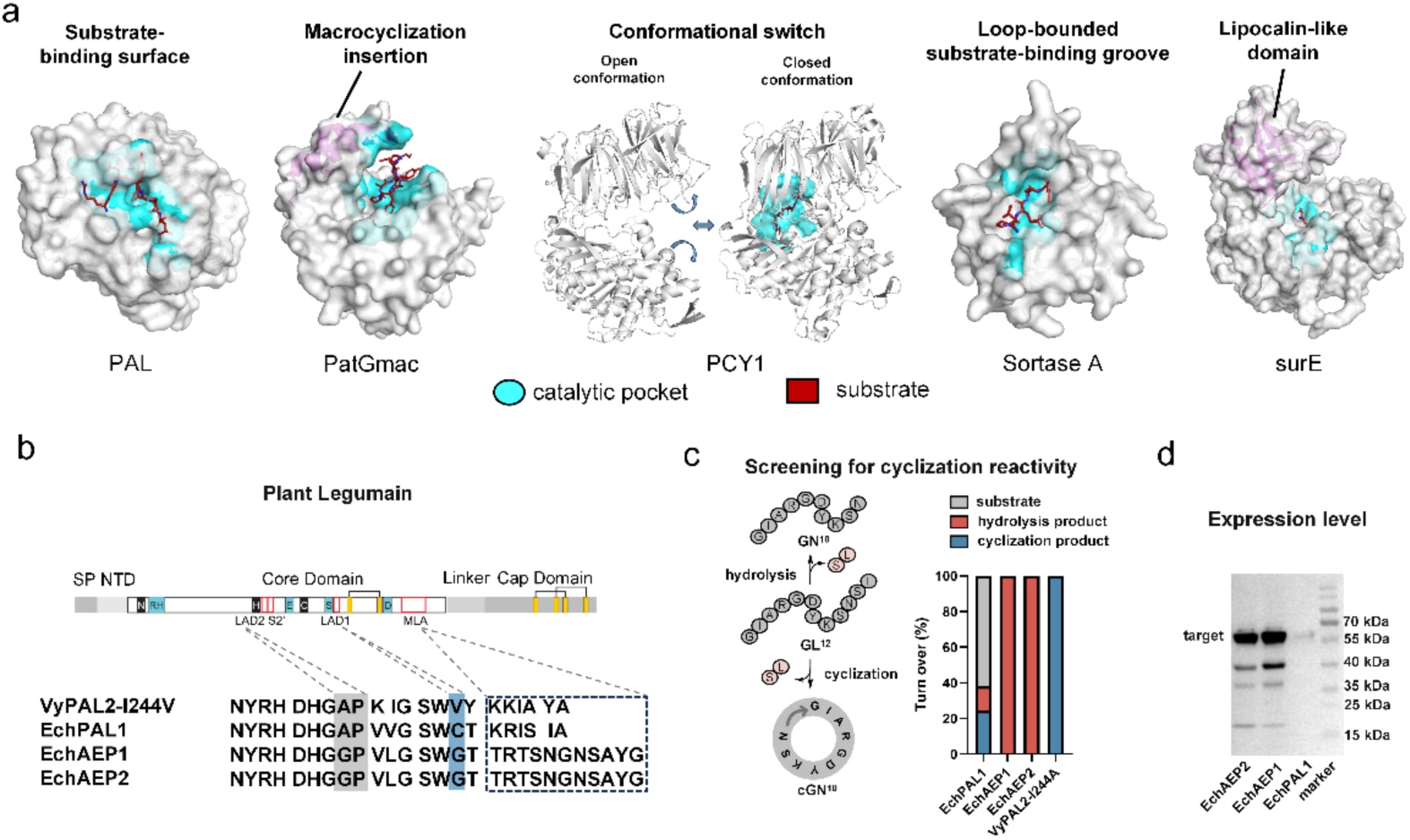
Mining and activity characterization of legumain enzymes from *Exallage chrysotricha*. (a) Comparison of substrate-binding modes of PAL, PatGmac, PCY1, Sortase A, and SurE. Catalytic pockets are shown in blue, peptide ligands in red, and the featured structures “macrocyclization insertion” of PatGmac and “lipocalin-like domain” of SurE in purple. The double-headed arrow indicates the transition between open and closed conformations of PCY1. (b) LAD-site sequences and MLA-region features of natural enzymes from *Exallage*. (c) Schematic of the cyclization reaction and cyclization assays of natural *Exallage* legumains. Peptide substrate GL12 was incubated with 10 µL of crude enzyme supernatant in citrate buffer at pH 5.0 and 37 °C for 1 h. Blue, cyclic product cGN10. Red, hydrolysis product GN10. (d) Recombinant expression levels of *Exallage* legumains compared using Western blotting.

Previous work has implicated interactions between prime-side substrate and the S2′ pocket influence reaction-pathway partitioning by affecting displacement of catalytic water and the stability of the acyl-enzyme intermediate, and in human legumain prime-side substrate affinity, together with the conformation and protonation state of the catalytic cysteine, has been linked to transpeptidation activity^17^ ^18^. LAD1, however, is located in the non-prime-side S2 region, and how a substitution there influences a competition between water and an incoming peptide that take place on the prime side remains unclear^19^. These questions highlight gaps in our understanding of the mechanisms governing reaction selectivity in legumains and underscore the need to clarify the relationship between local conformational organization and catalytic selectivity. We recently showed that conformational preorganization of the catalytic center governs the efficiency of acyl-enzyme formation and its pH dependence in this family^20^. Whether the conformational properties of the intermediate also determine which nucleophile goes on to react with it has not been examined.

Here, we extensively mined sequence information from the existing collection of natural enzymes and used directed evolution to obtain a pair of contrasting enzymes differing by only a single residue. By combining enzymatic characterization, crystal-structure analysis, and molecular dynamics simulations, we investigated the molecular basis of ligation-hydrolysis partitioning, focusing on the relationship between active-site conformational organization and catalytic pathway selectivity. Our results indicate that the oxyanion hole influences the partitioning between ligation and hydrolysis through conformational rearrangements triggered by the binding of an incoming peptide. These findings advance our understanding of the principles underlying catalytic divergence in plant legumains and provide a mechanistic foundation for the directed engineering and de novo design of peptide ligases.

## Results

### Legumains from *Exallage chrysotricha* Reveals limitation of LAD-based understanding of Ligation-Hydrolysis Selectivity

Previous mining of natural PALs results in enrichment of *Viola* PALs, limiting the sequence diversity of the ligases examined. Because the genus *Hedyotis* is also known for cyclic peptide production^21^, we selected widely distributed *Exallage chrysotricha* (basinonym. of *Hedyotis chrysotricha*^22^) as the model plant. Transcriptome sequencing and molecular cloning yielded three legumain sequences, which were named as EchAEP1, EchAEP2, and EchPAL1, respectively, based on the sequence features of LAD1, LAD2, and MLA regions (**Figure 1b**). We first assessed cyclization activity with crude enzyme using GL12 (GIARGDYKSNSL) as a model substrate at pH 5.0. At 1 h checkpoint, EchAEP1 and EchAEP2 completely hydrolyzed the substrate. Surprisingly, although EchPAL1 was predicted to be a typical PAL, only 39% substrate was reacted and the hydrolytic product accounted for 1/3 of the conversion, indicating substantially weaker cyclization performance than that of the natural ligase VyPAL2 (**Figure 1c**). This result exposed limitations in the LAD-based prediction of reaction selectivity and suggested that additional sequence features may contribute to pathway partitioning. Because EchPAL1 exhibited very low expression, we selected EchAEP1, the most highly expressed legumain from the same plant, for subsequent mechanistic investigations (**Figure 1d**).

### Systematic Sequence Exploration Yields Variants with Activities Approaching Those of Natural Ligases

To fully exploit the evolutionary information contained in the natural enzyme collection, we first extracted ESMC 600M embeddings for 49 natural legumains with reported activities. Principal component analysis (PCA) of these embeddings revealed a two-dimensional sequence landscape in which AEPs, hybrid legumains (HYB, either LAD1 or LAD2 site is ligase-type), and PALs exhibited a broad progression along PC1 toward increasing ligation preference, with EchAEP1 positioned at the negative end of PC1 within the AEP cluster (**Figure S1**). Because this collection contained relatively few AEPs, we added 1,334 LAD-predicted AEP sequences from a database of 1,500 legumains^19^ to supplement the negative class. Their centroid in sequence space was close to that of experimentally confirmed AEPs, providing preliminary support for this data-augmentation strategy (**Supplementary Note 1.1, Figure S2**). We then applied a sliding-window strategy with bag-of-words features and built classifiers based on local sequence segments using four machine-learning algorithms: decision tree (DT), logistic regression (LR), random forest (RF), and support vector machine (SVM)^23^ (**Figure S3**). After parameter optimization and cross-validation, we selected two complementary settings, a small window, w2 (word length 1), and a large window, w9 (word length 2), to identify sequence features associated with catalytic reaction selectivity at different scales (**Figure S4**). Using the first reported PAL, butelase-1, as the reference sequence, we calculated the submodel evaluation scores for individual positions and mapped them onto the protein structure. Key regions that were discontinuous in sequence were found to cluster mainly on the reaction-binding surface (**Figure 2a,b**). In particular, LAD1 and LAD2 ranked prominently with the smaller w2 window, whereas the larger w9 window highlighted the MLA region. An ensemble model constructed from the selected submodels achieved classification accuracies of 95.92% and 97.96% for w2 and w9, respectively, when evaluated retrospectively on the training set of 49 natural enzymes (**Figure S5**).

**Figure 2.**
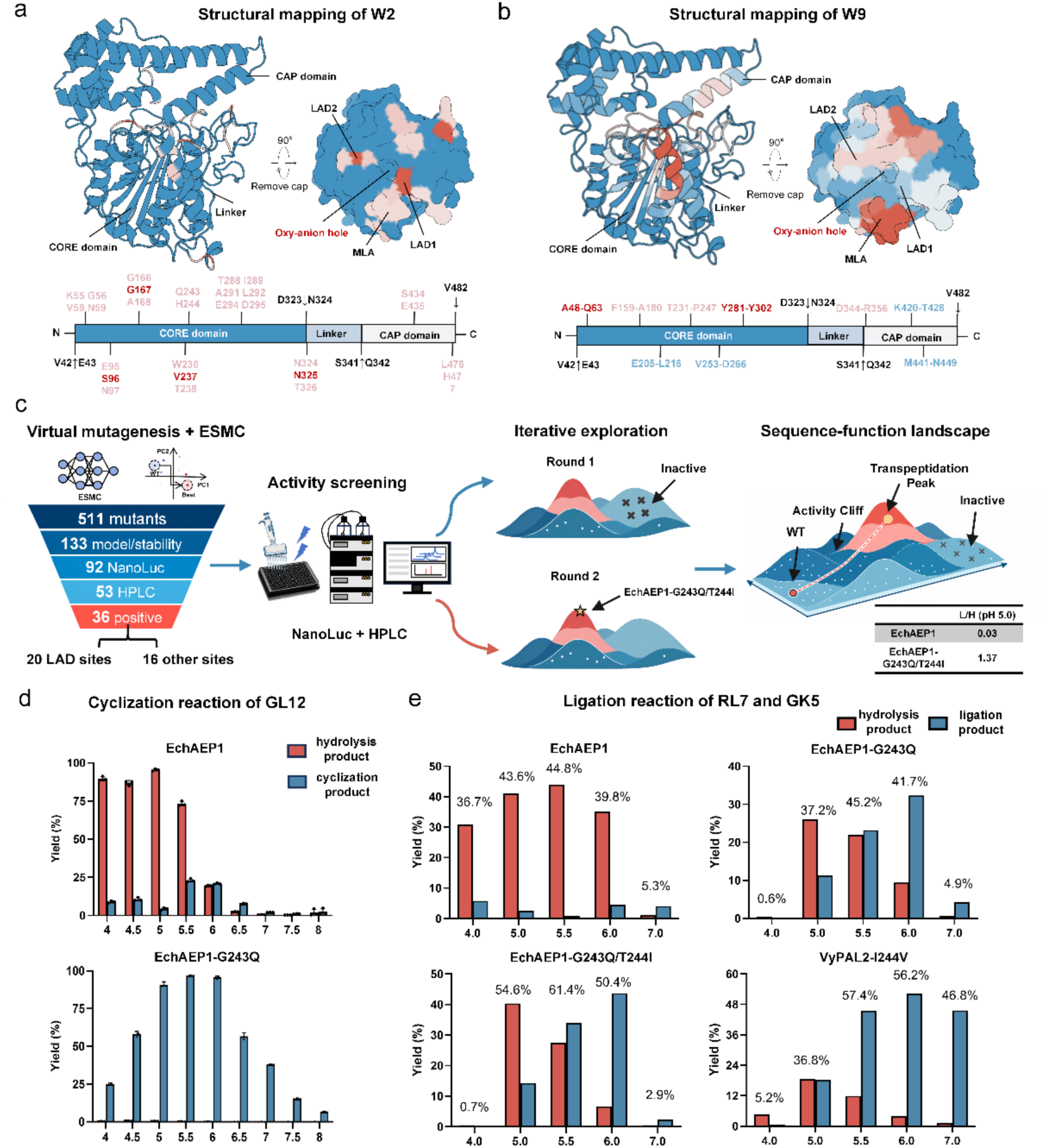
Systematic sequence exploration and characterization of purified enzymes. (a) Assessment of residue-level classification importance by ensemble identifier with a window size of 2 and (b) a window size of 9. Deeper red indicates higher classification importance. (c) Sequence-space exploration using an ensemble model and a protein language model (Supplementary Note 1.2). Virtual mutagenesis and ESMC embeddings guided variant selection, followed by iterative NanoLuc and HPLC screening. A total of 511 mutants were evaluated, yielding 36 positive variants from 20 LAD-associated and 16 additional sites. Iterative exploration identified an improved variant, EchAEP1-G243Q/T244I, with an increased ligation-to-hydrolysis (L/H) ratio at pH 5.0.(d) Characterization of cyclization reactions catalyzed by EchAEP1 and EchAEP1-G243Q at different pH values. Enzyme, 40 nM; peptide substrate GL12, 20 µM; reaction at 37 °C for 15 min. (e) Characterization of ligation reactions catalyzed by EchAEP1, EchAEP1-G243Q, EchAEP1-G243Q/T244I, and VyPAL2-I244V at different pH values. Enzyme, 80 nM; substrate A1 (Ac-RDGYNSL-NH2), 20 µM; substrate A2 (GIKRK), 100 µM. Reactions were conducted at 37 °C for 1, 60, 30, and 30 min for EchAEP1, EchAEP1-G243Q, EchAEP1-G243Q/T244I, and VyPAL2-I244V, respectively, before data collection.

We next performed virtual single-site saturation mutagenesis of the EchAEP1 core domain, obtaining 133 candidate single substitutions after joint screening by model score and stability (**Figure 2c**). Of these, 92 were selected for experimental validation based on the distribution of candidate positions. Following preliminary NanoLuc screening^24^ and quantitative liquid-chromatography analysis, we obtained reaction data for 49 single substitutions (**Figure S6**). 36 performed better than the wild type, notably including 33 beneficial substitutions outside the residue combinations specified by the LAD hypothesis. Of these, 17 occurred at known LAD positions and the other 16 were distributed across other positions (**Supplementary Note 1.2** and **Table S1**). These results demonstrate that with limited prior activity data, the model-guided strategy can broaden the exploration of functionally relevant positions and amino acid substitutions beyond those captured by conventional LAD rules.

The ESMC sequence space showed that single substitutions at EchAEP1 position 243 clustered around a cyclization-activity peak and other beneficial substitutions shifted toward this peak (**Figure S6**). We enumerated combinations of two to five beneficial substitutions and constructed a L/H ratio prediction model using round 0 single-point mutation data. The resulting combinations covered a broader region of sequence space, and candidates closer to the peak had higher predicted values (**Figure S7-S10**). By jointly considering predicted values and distances, we selected 30 combinatorial variants on the opposite side of the peak for experimental validation. Surprisingly, ligation activity did not continue to increase, instead, many expressed variants showed no detectable activity (**Figure S11**). In the second round, we sampled more densely around the ligation-activity peak (**Figure S12-S14**) and identified favorable double substitutions, 243Q-244I and 243N-244I, with L/H ratio values of 1.37 and 1.24, respectively, approaching the value of 1.51 for the positive control VyPAL2-I244V (**Figure S15**). The local sequence-function landscape revealed that changes at only a few residues could produce abrupt activity shifts, while a broad inactive region existed beyond the peak (**Figures S16-S19**). Analysis of candidate LAD sequences revealed a trajectory through embedding space from the hydrolytic region toward the transpeptidation peak without extending far into the inactive region. LAD2+ sequences remained near EchAEP1, whereas LAD1+ sequences and combinations of LAD1+ with LAD2+ approached or reached the peak (**Figure S20**).

Activity characterization of purified wild-type EchAEP1 and two variants, G243Q and G243Q/T244I (**Figure S21**), further confirmed that the single G243Q substitution was sufficient to markedly shift reaction selectivity toward peptide ligation over hydrolysis. G243Q produced only trace amount of hydrolysis by-products across the tested pH range of 4-8, as determined by HPLC analysis (**Figure 2d**). Even at the acidic pH of 4.5, when substrate conversion exceeded 50%, the C/H ratio still reached 58.1, surpassing reported butelase-2 and conLEG variants engineered by natural LAD substitutions^10^ ^11^. We next performed the more stringent intermolecular ligation assay. To minimize the influence of substrate conversion on the L/H ratio, we kept enzyme and substrate amounts constant and adjusted reaction times so that substrate conversion at pH 5.5 was within the 40-60% range. At approximately 45% conversion, G243Q increased the ligation-to-hydrolysis (L/H) ratio from 0.018 to 1.055, representing a 59-fold improvement over the wild type. The best combinatorial variant, EchAEP1-G243Q/T244I, showed a further increase in L/H to 1.24 at 61.4% conversion. When the pH was further increased to 6.0 and the reaction time extended, L/H reached 6.52 at 50.4% conversion, approaching the value observed for the natural ligase VyPAL2-I244V (**Figure 2e**).

### Crystal Structures Reveal Features of Oxyanion-Hole Preorganization

To investigate the structural basis for the enhanced ligation function of EchAEP1-G243Q, we conducted crystallographic studies of the wild type and the mutant. Because direct cocrystallization did not yield an enzyme-substrate complex, we drew on the substrate-like binding mode of the human legumain-cystatin E/M complex^25^ and determined the structure of the EchAEP1 catalytic domain in complex with CtCYS1, a phytocystatin from *Clitoria ternatea* (**Figure 3a**). To our knowledge, this represents the first crystal structure of a plant legumain-phytocystatin complex (2.3 Å resolution, PDB 9VA4). The side chain of CtCYS1 Asn179 inserts into the S1 pocket, while Ser180 and Leu181 correspond to the P1′ and P2′ positions, respectively. The binding position and local conformation of this fragment closely resemble those of substrate residues Asn5-Ser6-Leu7 in the Protenix^26^ predicted EchAEP1-RGDYNSL complex.

**Figure 3.**
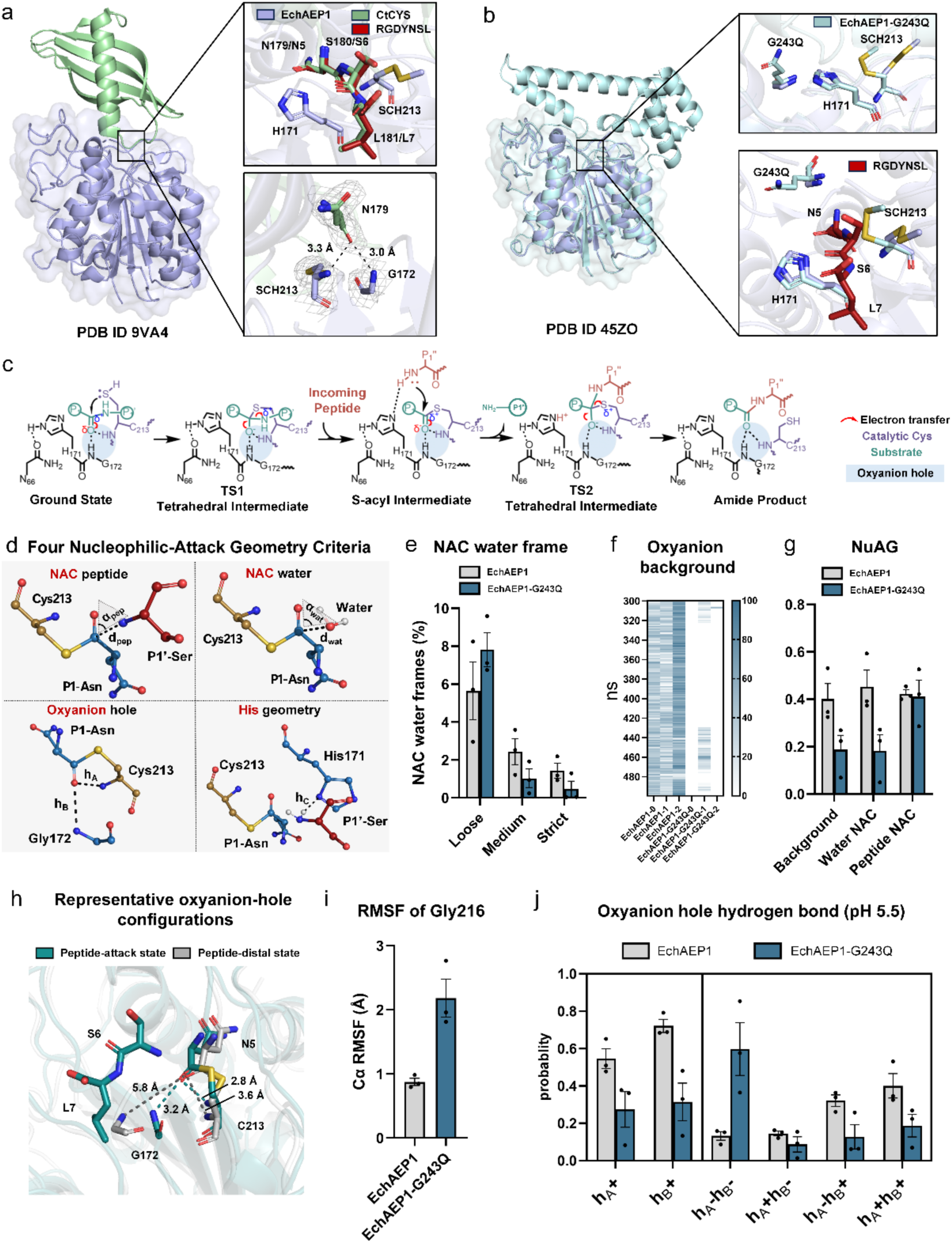
Structural and dynamic characterization of the catalytic center and oxyanion-hole formation in EchAEP1 and G243Q. (a) Crystal structure of EchAEP1 in complex with CtCYS1. (b) Crystal structure of the EchAEP1-G243Q zymogen. (c) Proposed mechanism of legumain-catalyzed peptide ligation. (d) Geometric criteria for nucleophilic attack on the thioester intermediate. d and α denote the nucleophile–carbonyl carbon distance and Nu–C–O angle, respectively; hA and hB indicate oxyanion-hole hydrogen bonds involving Cys213 and Gly172, respectively; hC denotes the hydrogen bond between the nucleophile and His171 ND1. (e) Fraction of trajectory frames satisfying three progressively stringent water-attack criteria (Loose_Wat, Medium_Wat, and Strict_Wat) in EchAEP1 and G243Q at pH 5.5. (f) Background probabilities of oxyanion-hole formation in EchAEP1 and G243Q during the 300–500 ns interval with the attacking peptide more than 10 Å from the catalytic center. Each band represents the percentage of frames with a formed oxyanion hole per 1 ns, colored from white (0%) to blue (100%). (g) Background and conditional probabilities of oxyanion-hole formation in EchAEP1 and G243Q at pH 5.5, with water- and peptide-attack conformations satisfying the Loose Nu criterion (Water NAC and Peptide NAC, respectively). (h) Representative oxyanion-hole configurations of EchAEP1-G243Q at pH 5.5. Superimposed snapshots show the peptide-attack (green) and peptide-distal (grey) states. Dashed lines indicate distances between the Asn5 thioester carbonyl oxygen and the backbone amide nitrogens of Gly172 and Cys213. (i) Residues showing large and statistically significant differences in fluctuation between EchAEP1 and EchAEP1-G243Q at pH 5.5. (j) Differences in formation probabilities of the two oxyanion-hole hydrogen bonds in EchAEP1 (grey) and EchAEP1-G243Q (blue) at pH 5.5. hA and hB represent the hydrogen bonds involving Cys213 and Gly172, respectively; “+” and “−” denote the formed and unformed states of a hydrogen bond.

In structure 9VA4, the backbone carbonyl oxygen of Asn179 lies 3.0 and 3.3 Å from the backbone amide nitrogens of Gly172 and catalytic Cys213, respectively, distances compatible with potential hydrogen bonding. This geometry suggests that a preorganized oxyanion hole is already present in the substrate-like bound state. Compared with the ordered-water-mediated hydrogen-bond network described for VdiPAL1, this structure more directly depicts the spatial relationship between a substrate-like carbonyl oxygen and the backbone amides of the catalytic center. A similar arrangement is also present in the human legumain-cystatin E/M complex (PDB 4N6O), suggesting that oxyanion-hole preorganization may be a relatively conserved structural feature of the legumain family.

Because attempts to obtain both a G243Q-CtCYS1 complex and a peptide-aldehyde adduct of G243Q were unsuccessful, we instead determined the crystal structure of its zymogen (PDB 45ZO) and compared it with the wild-type EchAEP1-CtCYS1 complex (PDB 9VA4) and the zymogen of a natural ligase VdiPAL1 (PDB 9VSX)^20^ (**Figure 3b**). Structural superposition showed that the introduced Gln243 side chain extends outward from the catalytic pocket. The catalytic residue Cys213 is present in an SCH-modified form in both EchAEP1 structures, but its side chain points away from the substrate reaction region in the wild-type complex and toward the substrate position in the mutant zymogen. Despite this difference, all three structures exhibited similar arrangements of the Cys and Gly backbone amides that form the oxyanion hole, suggesting that its underlying geometry is conserved across these enzyme states. However, because G243Q was crystallized as a zymogen rather than a substrate-bound or acyl-enzyme intermediate, these static structures cannot establish whether the oxyanion hole remains properly organized during different stages of catalysis. We therefore turned to molecular dynamics simulations to examine its conformational dynamics and selected EchAEP1 and G243Q, which differ by a single residue but exhibit markedly different ligation properties, as a model system.

### Molecular Dynamics Simulations Suggest a Nucleophile-Gated Oxyanion Hole

To investigate the molecular basis of reaction selectivity in legumains, we focused on the second, deacylation step by constructing thioester acyl-enzyme intermediates of EchAEP1 and G243Q for molecular dynamics simulations, in which a water molecule or the N-terminal amino group of a peptide competes to attack the carbonyl carbon of the thioester acyl-enzyme intermediate, producing hydrolysis or ligation products, respectively (**Figure 3c**). This process is influenced by nucleophilic-attack geometry, transition-state stabilization by the oxyanion hole, and histidine-assisted proton transfer (**Figure 3d**).

We first examined the hypothesis that ligases favor peptide ligation by restricting water access to the catalytic center. The free N-terminal amino group of the P1′ leaving peptide was used as a model for an incoming peptide nucleophile. Because water proximity alone does not necessarily indicate catalytic competence, we defined three progressively stringent criteria for water near-attack conformations (NACs), incorporating the catalytic requirements outlined above. The Loose_Wat criterion considered only nucleophilic attack distance (d_wat_) and angle (a_wat_); Medium_Wat additionally required oxyanion-hole formation with Cys213-NH and G172-NH; and Strict_Wat further incorporated the geometry required for His171-assisted proton transfer.

Applying these criteria revealed a striking distinction between water accessibility and catalytic competence. Contrary to the water-exclusion hypothesis, G243Q exhibited a higher frequency of geometrically favorable water-attack configurations than wild-type EchAEP1 (7.82% versus 5.64% of simulation frames under the Loose_Wat criterion, **Figure 3e**). However, incorporating the oxyanion-hole requirement reversed this trend, reducing the corresponding frequencies to 1.03% in G243Q and 2.44% in EchAEP1. Consequently, only 13% of the Loose_Wat configurations in G243Q satisfied the additional oxyanion-hole criterion, compared with 43% in EchAEP1. These results suggest that G243Q suppresses hydrolysis not by sterically excluding attacking water, but by reducing the population of water-attack configurations supported by a properly organized oxyanion hole. Thus, rather than acting as a constitutively preorganized catalytic feature, the oxyanion hole may serve as a dynamic determinant of reaction selectivity.

To further characterize the effect of G243Q on oxyanion-hole organization, we examined the distribution of hydrogen-bonding states in frames of which the released P1′ peptide had moved away from the catalytic center and was no longer positioned for nucleophilic attack. The oxyanion hole was classified into two states according to the hydrogen bond formation between the thioester carbonyl oxygen and the backbone NH groups of Gly172 and Cys213. Compared with wild-type EchAEP1, G243Q showed a pronounced shift toward configurations forming both hydrogen bonds, from 45.18 % in wt to 2.78 % in the mutant (**Figure 3f, Supplementary Note 1.3**). These observations suggest that G243Q destabilizes the preorganized oxyanion-hole configuration, potentially contributing to its markedly reduced hydrolytic activity. By analyzing frames with Ser-Leu associated, we observed that G243Q restored a wild-type-like probability of oxyanion-hole formation when the peptide nucleophile adopted a favorable attack geometry.

We next investigated whether this impaired oxyanion-hole organization was associated with the identity and attack geometry of the nucleophile. We defined nucleophile approach geometry (NuAG) criterion based on a loose attack distance (d_pep_ ≤4.5 Å) and angle (a_pep_ 70°-110°)^27^ for primary comparisons to balance sampling frequency and geometric plausibility. When water satisfied the NuAG criterion, the probabilities of oxyanion-hole formation were 0.45 for wild-type EchAEP1 and 0.18 for G243Q, close to their respective background probabilities (0.40 and 0.19). In contrast, when the peptide nucleophile adopted a favorable attack geometry, the probability in G243Q increased to 0.41, approaching that of wild-type EchAEP1 (0.42, **Figure 3g**). Representative snapshots of EchAEP1-G243Q at pH 5.5 illustrate intact and disrupted oxyanion-hole hydrogen-bond networks in the peptide-attack and peptide-distal conformations, respectively (**Figure 3h**).

Together, the probability analyses indicate that favorable peptide-attack geometries, but not water-attack geometries, are associated with substantially enhanced oxyanion-hole formation in G243Q. This nucleophile-dependent coupling may contribute to the preferential suppression of hydrolysis relative to peptide ligation observed experimentally.

To identify the structural changes underlying impaired oxyanion-hole organization in G243Q, we compared residue-level fluctuations between the two enzymes. Gly216 stood out as the only residue exhibiting both a substantial change in RMSF and statistical significance, with its RMSF increasing approximately 2.5-fold, from 0.87 in EchAEP1 to 2.18 in G243Q (**Figure 3i**). Notably, Gly216 is located on an α-helix immediately beneath Cys213, whose backbone NH group contributes one of the two oxyanion-hole hydrogen bonds (bond A). The increased mobility of this region may contribute to reduced oxyanion-hole stability. Indeed, hydrogen-bond analysis revealed that the formation probability of bond A decreased from 0.55 to 0.27, while the population of configurations lacking both oxyanion-hole hydrogen bonds increased markedly from 13% to 60% (**Figure 3j**). Interestingly, substitutions of the adjacent Ser215 with smaller residues (S215A and S215G) also favored ligation (**Figure S22**), suggesting that local structural perturbations in this region may influence reaction selectivity. These results link the G243Q substitution to altered local dynamics near Cys213 and a pronounced shift toward poorly organized oxyanion-hole configurations, providing a plausible structural explanation for the preferential suppression of hydrolysis.

### Reduced Oxyanion-Hole Organization Correlates with Ligation Selectivity across pH and Enzyme Backgrounds

To examine whether the relationship between oxyanion-hole organization and reaction selectivity extends beyond the G243Q substitution, we investigated the effects of pH and enzyme background on oxyanion-hole dynamics. Building on the analysis at pH 5.5 (**Figure 3e**), we extended the water-attack analysis to pH 7.0 (**Figure 4a**), where ligation was more strongly favored. The fraction of water NACs satisfying the oxyanion-hole requirement decreased from 43% to 31% in EchAEP1, whereas it increased modestly from 13% to 19% in G243Q.

**Figure 4.**
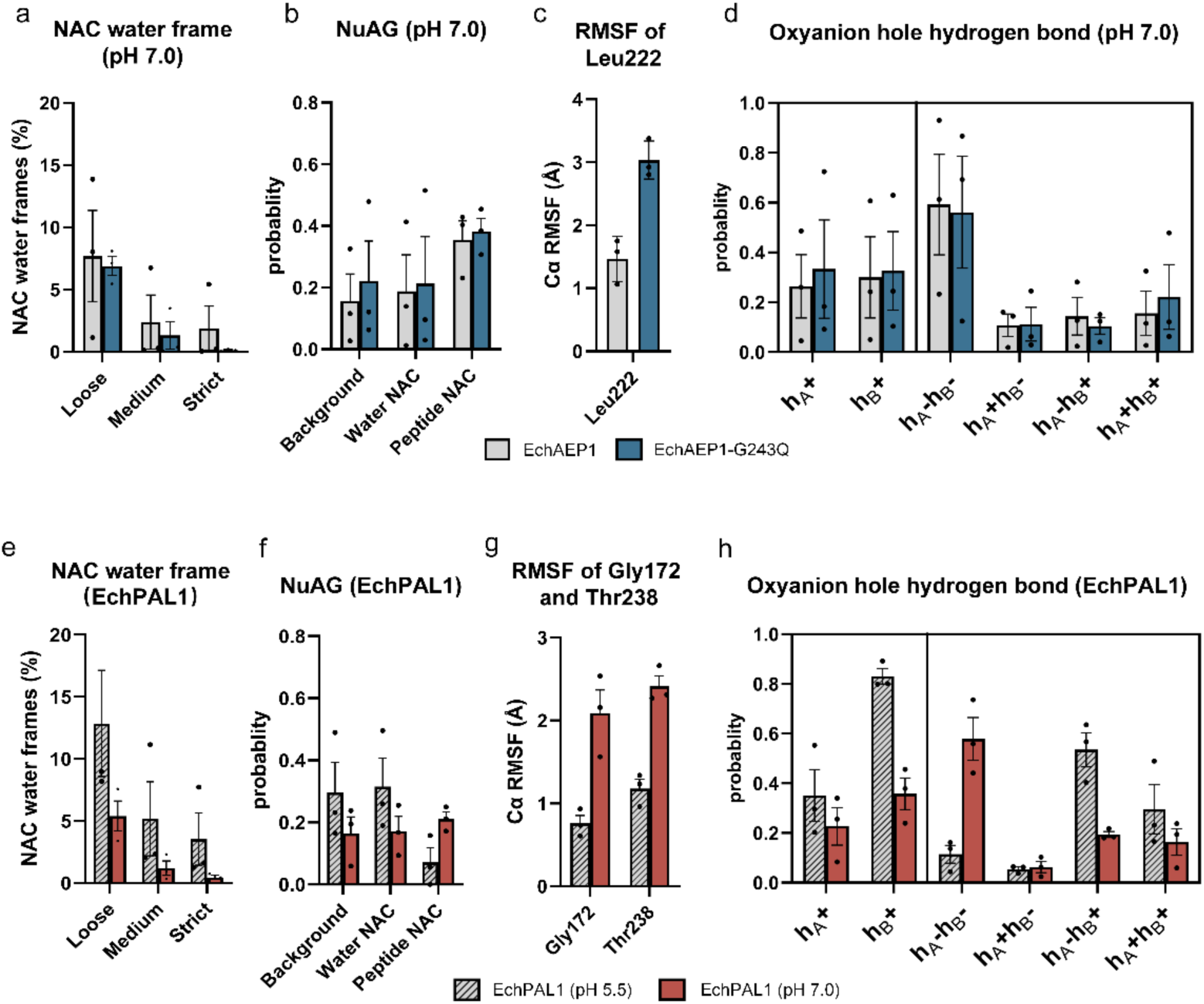
Effects of pH and enzyme background on oxyanion-hole dynamics. (a–d) Comparison of EchAEP1 (grey) and G243Q (blue) at pH 7.0. (a) Fractions of frames satisfying three water-attack criteria (Loose_Wat, Medium_Wat, and Strict_Wat). (b) Background and conditional oxyanion-hole formation probabilities under water- and peptide-attack configurations, defined using the NuAG criterion. (c) Residues exhibiting significant differences in RMSF exceeding the fluctuation-magnitude threshold. (d) Oxyanion-hole hydrogen-bond state probabilities. (e–h) Comparison of EchPAL1 at pH 5.5 and 7.0. (e) Fractions of frames satisfying the three water-attack criteria, with pH 5.5 values shown as hatched bars. Red, blue, and purple indicate Loose_Wat, Medium_Wat, and Strict_Wat, respectively. (f) Background and conditional oxyanion-hole formation probabilities (NuAG criterion). (g) Residues exhibiting significant pH-dependent RMSF differences exceeding the fluctuation-magnitude threshold. (h) Oxyanion-hole hydrogen-bond state probabilities. In panels d and h, hA and hB denote hydrogen bonds involving Cys213 and Gly172, respectively; “+” and “−” indicate formed and unformed states.

We next examined the background probability of oxyanion-hole formation. In EchAEP1, this probability decreased markedly from 0.40 at pH 5.5 to 0.16 at pH 7.0, approaching the consistently low values observed for G243Q (0.19 and 0.22, respectively; **Figures 3g** and **4b**). At pH 7.0, peptide-attack configurations in EchAEP1 were associated with an increased oxyanion-hole formation probability of 0.35, whereas water-attack configurations showed a probability close to background (0.19). At pH 7.0, the increased Gly216 flexibility previously observed in G243Q was no longer evident relative to EchAEP1, with only Leu222, located away from the oxyanion hole, meeting both the fluctuation-magnitude and statistical-significance criteria (**Figure 4c**). The oxyanion-hole hydrogen-bonding patterns of EchAEP1 also resembled those of G243Q (**Figure 4d**). Thus, increasing pH shifted EchAEP1 toward an oxyanion-hole response resembling that of G243Q at pH 5.5, consistent with the reduced difference in ligation selectivity between the two enzymes at higher pH.

We next examined EchPAL1, which was predicted to be a ligase but exhibiting low substrate conversion and predominantly hydrolytic activity. Increasing pH from 5.5 to 7.0 reduced both the background probability of oxyanion-hole formation (0.30 to 0.16) and the fraction of water NACs satisfying the oxyanion-hole requirement (40% to 22%; **Figure 4e, f**). However, its peptide-associated response differed from that of G243Q. At pH 5.5, favorable peptide-attack configurations were associated with a markedly lower oxyanion-hole probability (0.07) than the background (0.30), whereas at pH 7.0, the conditional probability increased slightly above the background (0.21 versus 0.16). These findings indicate that the relationship between peptide-attack geometry and oxyanion-hole formation is dependent on both enzyme background and pH, rather than representing a universally positive response.

Despite these differences in nucleophile-dependent responses, the distribution of oxyanion-hole hydrogen-bonding states revealed a striking common pattern. Configurations lacking both oxyanion-hole hydrogen bonds (A−B−) accounted for only 11% and 13% of frames in the hydrolysis-favoring EchPAL1 and EchAEP1 systems at pH 5.5, respectively. In contrast, these configurations represented 58%, 59%, 56%, and 60% of frames in EchPAL1 at pH 7.0, EchAEP1 at pH 7.0, G243Q at pH 7.0, and G243Q at pH 5.5, respectively (**Figure 3j, Figure 4d** and **h**). Thus, across the examined enzyme backgrounds and pH conditions, increased relative ligation selectivity was consistently associated with enrichment of configurations lacking both oxyanion-hole hydrogen bonds. This suggests that reduced basal oxyanion-hole organization may represent a shared conformational feature of ligation-favoring conditions, even when the underlying nucleophile-dependent responses differ.

To assess the robustness of these observations, we repeated the analyses using alternative geometric criteria (**Supplementary Note 1.4**). The distance-only peptide proximity criterion yielded results broadly consistent with NuAG, while water-associated trends were maintained under more stringent geometric criteria. Together, these cross-system comparisons support a consistent association between increased ligation selectivity and reduced oxyanion-hole organization, despite differences in nucleophile-dependent conformational responses across enzyme backgrounds.

## Discussion

Previous work on catalytic efficiency shows that the first rate-limiting acylation step in legumain catalysis correlates with the pH-dependent population of near-attack conformations^20^. The present study further shows that partitioning between ligation and hydrolysis during the second deacylation step is closely associated with the dynamic organization of the oxyanion-hole. While ligation selectivity in PALs has traditionally been explained by restricted water access to the acyl-enzyme intermediate, our simulations suggest an alternative mechanism: reaction selectivity is regulated by changes in the conformational occupancy of the oxyanion hole upon nucleophile association. The G243Q substitution shifts the conformational equilibrium to reduce the population of fully organized oxyanion-hole configurations when the peptide nucleophile is distant, whereas favorable peptide-attack geometries are associated with a restoration of their occupancy to wild-type-like levels. Consequently, water attack is less likely to occur in a catalytically competent configuration, whereas peptide attack can still benefit from a properly organized oxyanion hole, providing a possible explanation for the observed reversal in reaction selectivity (**Figure 5**). This nucleophile-gated oxyanion hole model highlights the importance of coupling between substrate geometry and catalytic-site dynamics, extending the understanding of reaction selectivity beyond static active-site structures.

**Figure 5.**
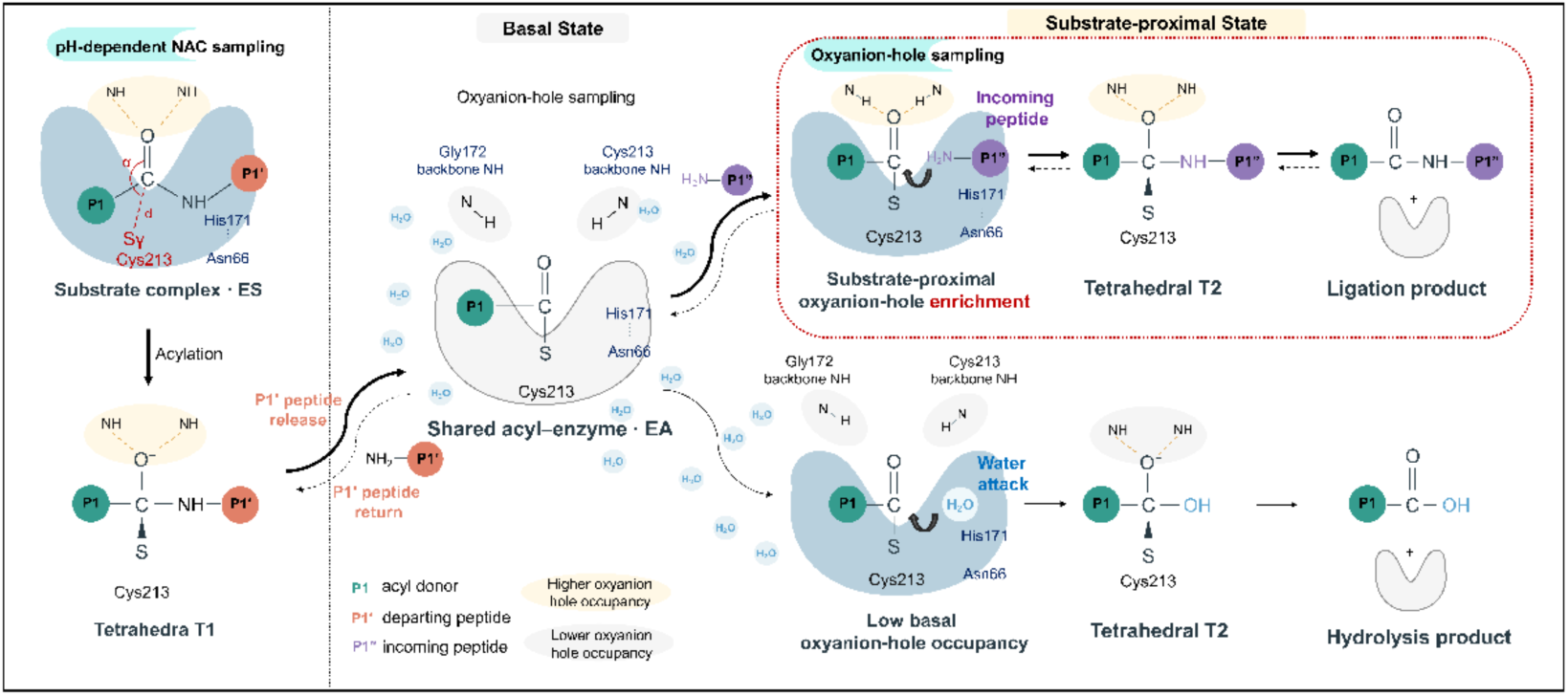
Proposed model of nucleophile-dependent reaction selectivity in legumains. During acylation, pH-dependent near-attack conformations (NACs) contribute to the formation of the shared thioester acyl-enzyme intermediate (EA) via the first tetrahedral intermediate (T1). During deacylation, low basal oxyanion-hole occupancy limits hydrolytically competent configurations despite water accessibility. In G243Q, favorable incoming-peptide attack geometries are associated with increased oxyanion-hole occupancy, favoring ligation. Both pathways proceed through the second tetrahedral intermediate (T2). P1, P1′, and P1″ represent the acyl donor, departing peptide, and incoming peptide, respectively.

More broadly, flexible peptide substrates may influence catalysis not only by supplying reactive groups but also by reshaping the conformational ensembles of the enzymes that recognize them^28^. In sortase B, the acyl-donor P1 threonine stabilizes a catalytic arginine, facilitating oxyanion-hole formation^29^. In contrast, in legumains, a properly organized oxyanion hole supports catalysis during both acylation and deacylation, but its coupling to substrate recognition may differ between these stages. During acylation, P1-Asn recognition helps position the native peptide substrate for productive attack by the catalytic cysteine. After acylation, however, our 500 ns simulations of the thioester acyl-enzyme intermediate showed that the oxyanion hole could remain fully formed even when the P1-Asn side chain disengaged from the S1 specificity pocket (**Supplementary Note 1.3**). These observations suggest that, after acylation, oxyanion-hole organization can be maintained without persistent anchoring of the acyl-donor P1 residue. Interestingly, recent demonstration of PAL-mediated ligation at non-Asx junctions using preactivated thioesters highlight the distinct chemical and recognition requirements associated with different catalytic stages, as the P1-Asn recognition required for efficient cleavage of native peptide substrates need not be maintained during subsequent ligation^30,31^. Our simulations may thus help explain the relaxed P1 specificity observed with preactivated thioesters. More importantly, our findings extend substrate-dependent oxyanion-hole organization from catalytic stabilization to reaction selectivity: favorable peptide-attack geometries, but not water-attack geometries, were associated with increased oxyanion-hole occupancy in G243Q. Such coupling may redistribute catalytically competent conformations and thereby influence the relative free-energy barriers of competing reaction pathways^13,32^.

Additional analysis of histidine-associated proton-transfer geometries revealed trends broadly consistent with experimental reaction partitioning (**Supplementary Note 2.1**). Although the fixed histidine tautomer precludes direct assessment of proton transfer, these observations highlight the importance of considering nucleophile activation alongside oxyanion-hole organization when evaluating catalytic competence. We speculate that changes in conformational entropy upon substrate binding, together with their coupling to enthalpic effects, may contribute to the catalytic efficiency and selectivity of these reactions. This perspective emphasizes the dynamic coupling between flexible peptide substrates and the active site and offers a useful approach for investigating other acyltransferases that recognize and transform flexible peptides: incorporating the substrate’s own conformational properties into mechanistic analysis, and examining how substrate-binding-driven redistribution of conformational ensembles affects competing reaction pathways, thereby extending the study of catalytic selectivity from static active-site structures to dynamic conformations and reaction free-energy landscapes.

Beyond the mechanistic findings, this study demonstrates the value of systematic, data-guided exploration of enzyme sequence–function relationships. Conventional enzyme engineering often relies on known functional residues or structural intuition to select mutation sites, potentially overlooking alternative substitutions and less obvious regulatory positions. By integrating protein-language-model representations, sliding-window exploration, predictive modeling, and iterative experimental screening, our strategy reduced reliance on such prior assumptions and enabled broader exploration of the mutational landscape using limited existing activity data. Importantly, this approach not only identified beneficial substitutions beyond conventional LAD rules but also independently converged on LAD1 as a major determinant of reaction selectivity, illustrating how systematic sequence exploration can complement established empirical rules, providing a more comprehensive basis for identifying functional residues and understanding their sequence-context dependence.

Together, our sequence-space and mechanistic analyses suggest that the oxyanion hole functions as a substrate-responsive catalytic gate, whose conformational organization is shaped by key residues, sequence context, and nucleophile-approach geometry. Rather than acting as universal molecular switches, LAD residues influence reaction selectivity through this dynamic catalytic environment. This complexity presents a fundamental challenge for de novo PAL design: reconstructing the conserved catalytic architecture or introducing known ligase-associated residues may not be sufficient to reproduce efficient ligation. A mechanistic understanding of how sequence context and substrate recognition shape the conformational states governing competing reaction pathways will therefore be essential for the rational design of peptide ligases with predictable reaction selectivity.

## Acknowledgement

We acknowledge the staff of beamline BL10U2 and BL02U1 at the Shanghai Synchrotron Radiation Facility for assistance during the data collection. This work was supported by the National Key Research and Development Program of China (2023YFA0916000), National Natural Science Foundation of China (32371324).

## Author contributions

S.Q.: conceptualization, methodology, datamining, modeling and simulation, computer-assisted engineering, writing-original draft, writing-editing; W.D.: conceptualization, wild-type and mutant library construction, data analysis, activity assays, crystal structure solving and refinement, writing-original draft, writing-editing; JH.M.: protein purification and crystallization; Y.G.: mutants construction, activity assays; JR.M.: discussion of molecular dynamics simulation details; X.H: conceptualization, methodology, writing-editing, supervision, funding.

## Data Availability

Authors declare that all data supporting the findings of this study are available within the paper and its supplementary information files.

