## supporting information for "A Substrate-Responsive Catalytic Gate for Peptide Ligation"

Supplementary Information

**Contents:**

Methods

Supplementary Figures

Supplementary Tables

Supplementary Note 1

Supplementary Note 2

**Methods：**

**Plant RNA extraction, transcriptome sequencing, and identification of legumain homologues**

Fresh *Exallage chrysotricha* plant samples were collected in the field. Total RNA was extracted from leaves using the TRIzol method and reverse-transcribed into complementary DNA (cDNA) for subsequent cloning of legumain genes. Fresh leaf samples were also submitted to BGI Genomics Co., Ltd. for transcriptome sequencing. Sequencing reads were assembled, coding sequences (CDSs) were predicted, and the predicted CDSs were translated into amino acid sequences to construct a protein sequence database. Using the amino acid sequence of the VyPAL2 proenzyme as the query, BLASTP homology searches were conducted. Candidate proteins with sequence identity greater than 60% and query coverage greater than 90% were selected as putative legumain family homologues for subsequent gene cloning and functional characterization.

**Cloning of legumain genes, recombinant expression in 24-well deep-well plates, and preparation of crude enzyme extracts**

Target legumain genes and mutant sequences were amplified by PCR and cloned into the pET28a(+) expression vector using NdeI and XhoI restriction enzymes, yielding recombinant proteins with 6×His tags at both the N and C termini. Sequence-verified recombinant plasmids were transformed into competent *Escherichia coli* SHuffle T7 cells. A single positive colony was inoculated into liquid LB medium and grown overnight to prepare a seed culture. The seed culture was transferred at 1% (v/v) into a 24-well deep-well culture plate and shaken at 37 °C and 220 rpm until the OD₆₀₀ reached approximately 0.4. Isopropyl β-D-1-thiogalactopyranoside (IPTG) was then added to a final concentration of 0.1 mM, and induction was continued at 16 °C and 180 rpm for 24–48 h.

After induction, cells were harvested by centrifugation and resuspended in lysis buffer containing lysozyme (50 mM HEPES, 100 mM NaCl, 1 mM EDTA, 5 mM β-mercaptoethanol, and 0.1% Triton X-100, pH 7.5). The suspension was mixed thoroughly and lysed overnight at 4 °C. Cell debris was removed by centrifugation, and the resulting supernatant was collected as the crude enzyme preparation for subsequent activity assays.

**Classification based on LAD residues**

In the multiple-sequence alignment containing the Butelase1 reference sequence, positions 167, 168, and 237 were mapped to the corresponding alignment columns using the 1-based coordinates of the ungapped Butelase1 sequence. The residues at these columns were then extracted for each sequence. According to the LAD classification rules applied in this study, LAD1 was designated positive when the residue corresponding to position 237 was I, V, C, or P; LAD2 was designated positive when the consecutive residues corresponding to positions 167 and 168 formed GA, AA, or AP. All other cases were designated negative for the respective determinant.

Sequences negative for both LAD1 and LAD2 were assigned the AEP label, those positive for both were assigned the PAL label, and those positive for only one determinant were assigned the HYB label. These labels were inferred from sequence-based rules and did not replace experimentally determined functional classifications.

**Construction of the natural enzyme dataset and definition of labels**

We collected 49 legumain records with experimentally assigned functional labels, comprising 30 PALs, 12 HYBs, and 7 AEPs. This dataset included naturally occurring enzymes and engineered variants. For binary classification, PALs and HYBs were combined into the positive CYC class, while experimentally annotated AEPs formed the negative class. From 1,500 background legumain records, sequences containing disallowed characters were removed; sequences with a single terminal stop symbol were retained after removal of that symbol. Background records with sequences identical to those in the experimentally labeled set were then excluded, with experimental labels taking precedence.

Background sequences were classified using the rule-based procedure described under 'Classification based on LAD residues.' The 1,334 records lacking experimental labels but classified as AEP were designated LAD_AEP and used as candidate negative samples. The complete modeling source set therefore comprised 1,383 records: 42 CYC, 7 experimentally labeled AEP, and 1,334 LAD_AEP. Counts refer to records, and identical sequences associated with different IDs were not comprehensively deduplicated. LAD_AEP labels were inferred from sequence rules and were not considered individually experimentally validated AEP annotations.

**Extraction of protein sequence embeddings and principal component analysis**

Final-layer sequence embeddings were extracted using the ESMC-600M model. Token vectors, including those for the beginning-of-sequence (BOS) and end-of-sequence (EOS) markers, were arithmetically averaged to generate a 1,152-dimensional sequence-level embedding for each protein.

A matrix was constructed with sequences as samples and embedding dimensions as features. Principal component analysis (PCA) was performed for dimensionality reduction, and the first two principal components were retained for two-dimensional visualization of sequence space. PCA used full singular value decomposition with built-in feature centering, without feature standardization or whitening. The reference sets used to fit PCA are specified for each analysis. Sequences included only for supplementary visualization were embedded using the same procedure and projected with the mean and principal components fitted on the reference set; PCA was not refitted.

**Sequence-space analysis of experimentally annotated enzymes and the expanded sequence library**

The 49 experimentally annotated enzyme records described under 'Construction of the natural enzyme dataset and definition of labels,' and the expanded set of 1,383 records obtained by adding 1,334 LAD_AEP sequences, were embedded and visualized in two dimensions as described above. The experimentally annotated set included natural enzymes and engineered variants. PCA was fitted separately to the 49-record experimental set and the 1,383-record expanded set, and the resulting principal components were used to display their respective datasets. Because these PCA coordinate systems are different, changes in the position of the same enzyme between the two plots were not interpreted as movement within a single space. Analyses were conducted at the record level, retaining identical sequences associated with different IDs.

To display EchPAL1 as an additional reference, its 456-aa sequence, obtained by removing the N-terminal 27 residues, was embedded using the same procedure and projected into both pre-fitted PCA spaces. EchPAL1 was not included in PCA fitting, the original sliding-window analysis, or model training; it was displayed as HYB on the basis of its existing experimental classification. Accordingly, the experimental-set and expanded-set plots contain 50 and 1,384 records, respectively, whereas category-composition plots and model development retain the original totals of 49 and 1,383 records.

The centroid of each category was defined as the arithmetic mean of its two-dimensional coordinates. For the expanded-set plot, nominal 95% bivariate normal data-distribution ellipses were drawn using the within-group sample covariance matrix S, with boundaries defined by (x−μ)ᵀS⁻¹(x−μ)=5.9915, where μ denotes the group centroid and covariance is calculated using n−1 in the denominator. EchPAL1 was included in HYB centroid and covariance calculations for this display. The ellipses describe the dispersion of sample observations and do not represent confidence regions for category means.

**Multiple-sequence alignment and sliding-window features**

Sliding windows were extracted from the MUSCLE multiple-sequence alignment using the full-length 482-aa Butelase1 sequence as reference and a stride of one residue. Window endpoints were expressed as 1-based inclusive reference coordinates. After reference endpoints were mapped to MSA columns, the corresponding aligned segments were extracted for each sequence and gap characters were removed. Owing to insertions and deletions, the actual length of a segment extracted from an individual sequence could differ from the reference window length.

Reference window lengths w from 2 to 9 were evaluated. For each w, fixed character n-gram lengths n from 1 to w−1 were tested, yielding 36 configurations. Character-count bag-of-words features were generated using CountVectorizer(analyzer="char", ngram_range=(n,n)), with feature extraction and the classifier included in a single training pipeline. Bag-of-words features did not encode absolute positions of n-grams within a segment: n=1 counted individual amino acids, whereas n=2 counted adjacent amino acid pairs.

**Balanced sampling and repeated cross-validation**

The aforementioned 1,383 records served as the source population for both sliding-window analysis and subsequent modeling. In each sampling round, all 42 CYC and 7 experimentally labeled AEP records were retained, and 35 additional records were randomly sampled without replacement from the 1,334 LAD_AEP records, producing a balanced set of 84 records with 42 positive and 42 negative examples.

Each window was evaluated with logistic regression (LR), support vector machines (SVM), decision trees (DT), and random forests (RF). Every combination of algorithm, window, and n-gram length was evaluated five times, with auxiliary negative samples resampled in each repeat and randomized stratified fivefold cross-validation performed. The default parameters of the respective scikit-learn classifiers were used, without additional class weighting.

Cross-validation performance was recorded using the scikit-learn ROC-AUC and F1 scorers. Within each repeat, scores were averaged over five folds, and the five repeat means were subsequently averaged to yield a window score. The classification score at a reference residue was defined as the arithmetic mean of scores for all windows spanning that position. An overall configuration score was obtained by further averaging the position scores across all 482 reference positions. This aggregate score was used for descriptive comparison of configurations and was not treated as the overall ROC-AUC or F1 of an independent classifier on a test set.

Within each configuration and algorithm, scores for the 482 positions were ranked in descending order. The average rank for ties was divided by 482 to obtain a rank fraction, with smaller values indicating higher rankings. Random states for repeated sample selection, fold assignment, and classifiers were not fixed in the original workflow. ROC-AUC and F1 were evaluated through separate cross-validation scoring calls and therefore were not guaranteed to use identical fold partitions.

Figures 2d and 2e used w2/n1 and w3–w9/n2 configurations. For Figure 2e, the mean score of the windows covering each position was raised to the 15th power and multiplied by 100 for display, with separate color limits assigned to each algorithm panel. This transformation served only to emphasize high-scoring regions and did not change the ranking of positions; subsequent threshold selection used the untransformed scores.

**Selection of informative windows and training of submodels**

Two configurations, w2/n1 and w9/n2, were selected for further modeling, representing single-character counts in two-residue reference windows and two-character counts in nine-residue windows, respectively. The mean cross-validated ROC-AUC of each window, before the visualization transformation, was used for selection, with thresholds of at least 0.90 for w2 and 0.95 for w9. If multiple algorithms qualified for a given window, the algorithm with the highest mean score was retained; ties were resolved using the first algorithm listed in the results table. The window/n-gram configurations and these thresholds were selected exploratorily, rather than established as optimal by an independent test.

Twenty submodels were independently trained for each selected window–algorithm combination. In each round, 35 LAD_AEP records were newly sampled to form a balanced set of 84 observations, which was randomly split in a stratified 4:1 ratio for fitting and internal holdout evaluation. Feature extraction was fitted only to the training subset. Default training parameters were used without a classifier hyperparameter grid search, and probability=True was specified for the SVM during training. The fitted models were saved directly without retraining on all 84 records or all 1,383 source records after evaluation.

Submodel weights were derived from hard-label scores on the internal holdout set. Specifically, binary predicted labels were passed to roc_auc_score; for binary classification, the resulting internal holdout score equals balanced accuracy, (sensitivity + specificity)/2. The mean and minimum scores across 20 submodels were summarized for each window. The mean-score threshold was set to 0, while minimum-score thresholds of at least 0.70 and 0.75 were applied to w2 and w9, respectively. All 20 submodels from a qualifying window were retained. Because the internal holdout score was used for both window retention and voting weights, this holdout set was part of model development.

**Structural mapping of position classification scores**

Using the Butelase1 structure, the average score of the 20 submodels associated with each qualifying window was assigned to all reference positions covered by that window, with zero contribution outside the window. Contributions from overlapping windows were summed for each position to obtain cumulative scores, without dividing by either window length or the number of covering windows. Scores for all 482 positions were min–max normalized as Bᵢ = 100 × (Tᵢ − min T)/(max T − min T) and written to the B-factor field of the PDB file for structural coloring. Positions with zero scores were included in the minimum, and normalization was performed separately for w2 and w9.

These structural scores reflect both window classification performance and window coverage density. The B-factor field was used solely as a visualization carrier; the values do not represent experimental temperature factors or structure-prediction confidence and were not used to compare absolute scores across configurations.

**Ensemble classification and retrospective evaluation of experimentally annotated records**

The Butelase1 reference sequence was added to each sequence set to be predicted, and MUSCLE alignment was performed using the -align, -output, and -perturb 42 options. Segments corresponding to qualifying reference windows were extracted and entered into the associated trained submodels. For submodel j, let yⱼ(x)=1 denote a CYC prediction, yⱼ(x)=0 an AEP prediction, and aⱼ its weight. The ensemble score was then defined as:

S(x)=M⁻¹ Σⱼ aⱼ[2yⱼ(x)−1]

Here, M is the total number of submodels contributing to prediction. The score is a signed weighted voting value, with the number of models rather than the sum of weights in the denominator. Sequences with S>0 were classified as CYC, whereas those with S≤0 were classified as AEP. S was not interpreted as a classification probability.

The original 49 experimentally annotated records were retrospectively evaluated after excluding the additionally introduced reference sequence. With CYC as the positive class, confusion matrices, accuracy, balanced accuracy, and the Matthews correlation coefficient were calculated. Because these records overlapped with the model-development source data and predictions were not restricted to submodels that had held each record out, this analysis constituted a retrospective evaluation rather than an independent test or out-of-fold assessment.

**Stability prediction using Rosetta Cartesian ΔΔG**

Effects of amino acid substitutions on protein stability were predicted using the Rosetta 2020.08.61146 Cartesian ΔΔG approach, with RosettaDDGPrediction 0.0.1 used for batch scheduling and result aggregation. All target proteins were analyzed with the same workflow and parameters. Starting from the reference structure of each target protein, a Cartesian FastRelax pre-optimization was first performed with the ref2015_cart scoring function and four internal repeats. For each target and batch, one pre-optimized structure was used as the common starting structure for the reference sequence and its mutants.

Subsequently, five rounds of local structural optimization were performed for both reference and mutant using the legacy implementation of cartesian_ddg, with bbnbrs=1 for neighboring backbone residues and fa_max_dis=9.0 Å in the ΔΔG step. Combined substitutions were introduced simultaneously into the same structure. Following the cartddg2020 aggregation procedure of RosettaDDGPrediction, the round with the lowest total mutant energy was selected. The difference between the total energy of that mutant and that of the reference structure from the correspondingly numbered round was computed (mutant minus reference) and divided by 2.94 to convert to kcal/mol. Negative ΔΔG indicated predicted stabilization, whereas positive ΔΔG indicated predicted destabilization.

**In silico single-site saturation mutagenesis of EchAEP1 and initial candidate selection**

EchAEP1 and Butelase1 were pairwise aligned with MUSCLE, and the endpoints of qualifying reference windows were mapped to MSA columns. All nongap EchAEP1 residues within the covered intervals were collected and deduplicated, including EchAEP1 insertions relative to the reference within these windows. At each selected position, the other 19 standard amino acid substitutions were generated in silico while retaining the alignment-column correspondence; individual mutant sequences were not realigned.

The w2 and w9 ensemble models were used to score EchAEP1 wild type and each mutant, and ΔS = S(mutant) − S(wild type) was calculated (output field: ave_change). The union of substitutions with ΔS>0 under either configuration was taken, and mutations at positions 26–301 of the original EchAEP1 sequence were retained as the candidate set for the core region. ΔS was used only to prioritize mutations and was not interpreted directly as a change in ligation activity, ligation-to-hydrolysis ratio, or stability.

To reduce the potential influence of conformational uncertainty in the N-terminal propeptide region of EchAEP1 on subsequent Rosetta ΔΔG estimates, the first 25 residues of the input sequence were removed before structural modeling, retaining the core and cap regions. Candidates were retained if Rosetta ΔΔG < 4.5 and at least one of the following was satisfied: a w2 model-score increase >0.11 or a w9 model-score increase >0.01. Positions enriched in candidate substitutions were then selected according to the site distribution, together with one candidate substitution at position 146, to construct the set for fluorescence screening.

**Recombinant expression, protein purification, and acid-induced autoactivation of EchAEP1 and its mutants**

Recombinant expression plasmids encoding EchAEP1, EchAEP1-G243Q, and EchAEP1-CL5 were individually transformed into competent *Escherichia coli* SHuffle T7 cells, and positive single colonies were selected for 1-L fermentation cultures. After induction, cells were harvested by centrifugation and resuspended in lysis buffer (50 mM HEPES, 100 mM NaCl, 1 mM EDTA, and 5 mM β-mercaptoethanol, pH 7.5) at a ratio of 10 mL per 1 g wet cell pellet. Cells were disrupted by sonication on ice, and cell debris was removed by centrifugation at 12,000 × g for 30 min at 4 °C. The supernatant was filtered and applied to a HisTrap HP nickel-affinity chromatography column. His₆-tagged proteins were eluted with buffer containing 50 mM HEPES, 100 mM NaCl, 1 mM EDTA, 5 mM β-mercaptoethanol, and 500 mM imidazole (pH 7.5). Fractions containing target protein were pooled, concentrated with Amicon® Ultra centrifugal filters (Merck & Co., Inc., USA), and buffer-exchanged into binding buffer (50 mM HEPES, 100 mM NaCl, 1 mM EDTA, and 5 mM β-mercaptoethanol, pH 7.5) to remove residual imidazole.

Purified proenzyme solutions were adjusted to 1 mg/mL, acidified to pH 4.2, and incubated at 37 °C for 30–120 min to induce autocatalytic activation. Activated proteins were further purified by size-exclusion chromatography on a HiLoad 16/600 Superdex 75 column with sodium citrate buffer (20 mM sodium citrate, 100 mM NaCl, 1 mM EDTA, 5 mM β-mercaptoethanol, and 5% (w/v) glycerol, pH 4.2) as the mobile phase. Target fractions were collected, and protein concentrations were calculated from the absorbance at 280 nm (A₂₈₀) measured on a NanoDrop 2000 spectrophotometer. Purified proteins were adjusted to pH 5.0 and stored at 4 °C for short-term use or supplemented with sucrose to a final concentration of 20% and stored at −80 °C for long-term use.

**Post-fermentation processing of single mutants and rapid NanoLuc activity screening**

Crude enzyme supernatants obtained from expressed mutants as described above were screened for peptide ligation activity using a chemiluminescence assay based on NanoLuc luciferase complementation. LigBiT and SmBiT were used as ligation substrates at final concentrations of 25 μM and 100 μM, respectively. Crude enzyme (12 μL) was added to a 25-μL reaction containing 20 mM sodium citrate (pH 5.0), 1 mM EDTA, and 5 mM β-mercaptoethanol and incubated at 37 °C for 30 min. After the reaction, a 4-μL aliquot was added to 100 μL phosphate-buffered saline (PBS) containing 5 μM furimazine. Peptide ligation products were detected by NanoLuc-catalyzed chemiluminescence, with the signal at 460 nm used to evaluate ligation activity. Screening results are provided in Table X.

**HPLC-based screening of cyclization and ligation activities of mutants**

The 53 favorable mutants identified in the initial NanoLuc chemiluminescence screen were further assessed for peptide cyclization and ligation by high-performance liquid chromatography (HPLC). Cyclization assays were performed in 20 mM sodium citrate buffer containing 1 mM EDTA and 5 mM β-mercaptoethanol (pH 5.0), with 40 μM GL12 peptide (GIARGDYKSNSL) as substrate and 0.05–10 μL crude enzyme. Reactions were conducted at 37 °C for 0.5–120 min. The volume of crude enzyme and reaction time were adjusted according to the catalytic activity of each mutant. Products were analyzed by HPLC, and observations with substrate conversion between 20% and 60% were retained for subsequent analysis of cyclization and hydrolysis.

Peptide ligation reactions used the same buffer, with A1 (Ac-RGDYNSL-NH₂) and A2 (GIKRK) as substrates at final concentrations of 20 μM and 100 μM, respectively, and 10 μL crude enzyme. Reactions were performed at 37 °C, with durations adjusted according to the activity of each mutant. Products were analyzed by HPLC, and observations with substrate conversion between 40% and 60% were selected for subsequent analysis of ligation and hydrolysis.

**Construction of the combinatorial candidate library**

Historical numerical selection criteria for positive single mutants were a mean conversion of 0.2–0.6 and C/H≥1.1, with both endpoints included. Combinations were generated by first selecting 2–5 distinct positions and then taking the Cartesian product of the permitted amino acid substitutions at each selected position. This yielded 46,582 combinatorial candidates, comprising 538 double, 3,886 triple, 14,581 quadruple, and 27,577 quintuple mutants.

**Construction of the C/H prediction model**

From cleaned data of prediction project round_1, 50 records with mean conversions in the inclusive interval 0.2–0.6 were selected. These comprised one EchAEP1 reference record and 49 single mutants; no additional C/H ≥ 1.1 criterion was applied. The 1,152-dimensional ESMC representations were used as input features and preprocessed C/H values as regression targets.

Using random seed 88, the data were randomly split 60:40 into a training set of 30 records and a holdout set of 20 records for model comparison. Pipelines comprising standardization, 15-component PCA, and a regressor were built to compare random forest, gradient boosting, XGBoost, support vector regression, Ridge, and Lasso. During model comparison, standardization, PCA, and regressors were fitted exclusively on the training set. Models were assessed on the holdout set using mean squared error, mean absolute error, and R², and the final model was selected by holdout R².

After random forest was selected, the complete pipeline was refitted using all 50 records and used to predict C/H across the historical combinatorial library. The saved random forest contained 100 trees, employed bootstrap sampling, used a maximum feature fraction of 1.0, imposed no maximum tree depth, and had minimum sample sizes of 2 for internal-node splitting and 1 for leaf nodes, with random seed 88. PCA retained 15 components. The saved model recorded scikit-learn version 1.7.1. The second screening round reused these predictions without refitting the C/H model.

Because the holdout set was also used for model selection, its performance metrics served for model comparison and did not constitute fully independent final generalization validation. Likewise, in-sample metrics after refitting on all records were not treated as independent predictive performance. C/H predictions were used only to prioritize candidates, not interpreted as direct predictions of L/H or as probabilities of candidate success.

**First-round candidate selection**

First, 2,609 candidates with predicted C/H≥50 were selected. The coordinate-wise equally weighted mean of the 1,152-dimensional ESMC vectors of seven reference single mutants—G243P, G243Q, G243Y, G243F, G243N, G243E, and G243R—was used as the reference centroid, and the Euclidean distance from each candidate to the centroid was calculated. Denoting the j-th reference vector as e_j, the centroid was μ=(Σe_j)/7 and the distance for candidate i was d_i=‖e_i−μ‖₂. The centroid and distances were calculated in the original embedding space, without using two-dimensional PCA coordinates or weighting by C/H.

Among the 2,609 candidates meeting the prediction threshold, distances were ranked in ascending order as percentiles, with tied distances assigned the minimum rank. Four right-closed intervals (0–25%, 25–50%, 50–75%, and 75–100%) contained 652, 652, 652, and 653 candidates, respectively. From these groups, 225, 75, 75, and 75 candidates, respectively, were selected by descending predicted C/H, yielding 450 candidates for stability-score filtering.

Using ΔΔG≤10 as the retention criterion, 36 candidates exceeding the threshold were removed. The remaining 414 retained their original distance groups, from which the top 15, 5, 5, and 5 predicted C/H candidates were chosen, respectively, yielding 30 first-round experimental candidates. Distance percentiles were not recalculated after stability filtering. Prediction ties retained the first-listed candidate: at the 450-candidate stage, the order in the original candidate cache was used; at the final 30-candidate stage, the row order after merging with the ΔΔG table as the left table was used.

This distance-stratified strategy increased sampling around the high-C/H reference neighborhood while retaining designs from more distant groups. Distance percentiles were not interpreted as calibrated model confidence.

**Second-round candidate selection and manual supplementation**

In the second round, the predicted C/H inclusion threshold was reduced to ≥30, yielding 35,887 candidates. The centroid was again the mean 1,152-dimensional vector of the seven reference single mutants. Euclidean distances were computed in the original embedding space and ranked as ascending percentiles among the candidates meeting this round's threshold. The top 0.5% were selected, yielding 179 candidates.

Precomputed ΔΔG values were retrieved for these 179 candidates. Based on the total score corresponding to `state=ddg`, 46 designs with ΔΔG≤5 were retained, using the kcal/mol units given in the input table. After removal of G146A_G243R, which duplicated a previously tested candidate, 45 new candidates remained.

In addition, 13 combinations were manually selected from insufficiently sampled regions of the two-dimensional distribution of all 46,582 historical candidates. This manual branch was not restricted to candidates with predicted C/H≥30 and did not apply the aforementioned ΔΔG threshold; selection of the regions relied on human judgment, and the identities are recorded in the saved manually selected list. Combining the 45 automatically selected and 13 manually selected candidates yielded 58 second-round candidates.

**Protein crystallization and crystal structure determination**

Purified EchAEP1 catalytic-domain protein was buffer-exchanged into 20 mM sodium citrate, 100 mM NaCl, 1 mM EDTA, 5% (w/v) glycerol, and 0.5 mM methyl methanethiosulfonate (MMTS), pH 4.2, and concentrated by ultrafiltration. EchAEP1 was mixed with highly purified CtCYS at a 1:1 molar ratio to a total protein-complex concentration of 13 mg/mL for initial crystallization screening. Screening was performed at 16 °C using the JCSG-plus™ HT-96/FX-96 crystallization screening kit, and crystals were observed after approximately 48 h. Crystals grew in 0.04 M potassium dihydrogen phosphate, 16% (w/v) polyethylene glycol 8000 (PEG 8000), and 20% (v/v) glycerol, pH 4.6. X-ray diffraction data for EchAEP1–CtCYS complex crystals were collected at beamline BL10U2 of the Shanghai Synchrotron Radiation Facility (SSRF) at 100 K.

Purified EchAEP1-G243Q proenzyme was concentrated to 6 mg/mL, and crystallization conditions were screened using the JCSG-plus™ HT-96/FX-96 kit. Crystals grew in 0.2 M tripotassium citrate monohydrate and 20% (w/v) PEG 3350. X-ray diffraction data were collected at beamline BL02U1 of the SSRF at 100 K.

Diffraction images were indexed and integrated using iMosflm. The AlphaFold 3-predicted protein structure served as the initial search model for phase determination by molecular replacement (MR) in PHASER. An initial structural model was then built in PHENIX and manually adjusted against electron-density maps using COOT. Iterative refinement was performed in PHENIX, including translation/libration/screw (TLS) refinement, until the final structure model was obtained. The determined crystal structures of the EchAEP1–CtCYS complex and EchAEP1-G243Q proenzyme were deposited in the Protein Data Bank (PDB) under accession codes 9VA4 and 45ZO, respectively.

**Structure prediction of enzyme–substrate complexes**

Initial complexes of wild-type EchAEP1, the G243Q variant, and EchPAL1 with the substrate peptide RGDYNSL were predicted independently using Protenix. The catalytic domains were extracted at the sequence level before structure prediction. The EchAEP1 domain comprised 278 residues corresponding to residues 51–328 of the full-length precursor; G193Q in this domain corresponds to G243Q in the full-length precursor. The EchPAL1 catalytic domain was delineated by sequence alignment with EchAEP1 using MUSCLE 5.3 and comprised 273 residues corresponding to residues 51–323 of the full-length precursor, or residues 24–296 of the input sequence lacking the N-terminal 27 residues.

For each prediction, one enzyme chain and one RGDYNSL peptide chain were supplied as separate protein chains, without an explicitly specified covalent linkage between them. Five candidate structures were obtained for each complex using random seed 101 and 10 recycling iterations. The EchPAL1 predictions used the protenix_base_20250630_v1.0.0 checkpoint with 200 diffusion sampling steps. This checkpoint is described in the Protenix model documentation. For each enzyme, the candidate with the highest Protenix ranking score among the five generated models, designated sample 0 in the saved outputs, was used for subsequent acyl–enzyme model construction. The same selected complex was used to prepare simulations at pH 5.5 and pH 7.0. Unless otherwise stated, enzyme residue identifiers refer to the full-length precursor sequences, whereas substrate residues are numbered according to their positions in RGDYNSL.

**Construction of acyl–enzyme intermediate models**

To represent the state after the first acylation step, the peptide bond between the backbone carbonyl carbon of substrate Asn5 and the backbone nitrogen of Ser6 was removed from the topology, and a thioester bond was established between the Sγ atom of catalytic Cys213 and the carbonyl carbon of Asn5. The departing Ser6–Leu7 dipeptide was retained in the simulation system, with the newly generated Ser6 N terminus modeled as neutral NH₂ to represent a potential incoming peptide nucleophile.

Local thioester atom types, partial charges, and bonded parameters were based on the CYSP-modified residue in the CHARMM36 force field. Boundary parameters connecting the modified region to the protein backbone were approximated by mapping chemically similar environments. The resulting acyl–enzyme models had fixed covalent topologies and were used to investigate conformational sampling and interactions near the reaction center. Bond formation, bond cleavage, and proton transfer were not simulated.

**Protonation states and molecular dynamics simulations**

Molecular dynamics simulations were performed with GROMACS 2023.2 using the CHARMM36-feb2021 force field and TIP3P water model, with Na⁺ ions added to neutralize the net charge. Protonation states corresponding to pH 5.5 and pH 7.0 were assigned before simulation according to PROPKA-predicted pKₐ values and the target pH and then held fixed throughout the simulations. Catalytic His171 was modeled in all systems as neutral HSE protonated at Nε, leaving ND1 available as a potential hydrogen-bond acceptor; other histidines were assigned according to their individual predicted protonation states. The newly formed Ser6 N terminus was kept as neutral NH₂ at both pH values. Thus, the simulations compared conformational sampling under different predefined protonation states.

After L-BFGS energy minimization, each system underwent 100 ps of position-restrained pre-equilibration. A 1-fs integration time step was used for equilibration, with V-rescale temperature coupling at 310.15 K and a coupling time constant of 0.2 ps, and isotropic Berendsen pressure coupling at 1 bar with a coupling time constant of 0.5 ps.

Production simulations used a 2-fs integration time step with bonds involving hydrogen atoms constrained. Separate V-rescale temperature-coupling groups were applied to protein and nonprotein atoms, each at 310.15 K with a coupling time constant of 0.2 ps. Pressure was controlled using isotropic Parrinello–Rahman coupling at 1 bar, a time constant of 2 ps, and compressibility of 4.5×10⁻⁵ bar⁻¹. Three-dimensional periodic boundary conditions and a Verlet neighbor list were used. Long-range electrostatics were calculated using particle-mesh Ewald summation, with real-space electrostatic and van der Waals cutoffs of 1.0 nm, and long-range dispersion corrections were applied to energy and pressure. Trajectories were saved every 50 ps.

Each protein–pH combination comprised three independent production trajectories initialized with different random seeds. For wild-type EchAEP1 and G243Q at both pH values, the first trajectory was 1 μs and the other two were 500 ns; all six EchPAL1 trajectories were 500 ns. For consistent comparisons, only the first 500 ns of each trajectory were used for Figures 4 and 5 and related supplementary analyses.

**Trajectory postprocessing and analysis window**

Trajectory postprocessing consisted sequentially of protein-centered coordinate processing, treatment of periodic boundaries, and overall translational and rotational fitting to the Protein group. Using MDAnalysis and project-specific analysis scripts, frames 0–10,000 of each trajectory were analyzed, including both endpoints (10,001 frames corresponding to 0–500 ns). The geometric analyses described below used the postprocessed trajectory coordinates directly, without additional minimum-image distance correction.

**Hierarchical screening of water molecules**

Water molecules were screened using three progressively stringent criteria, defined as Loose Wat, Medium Wat, and Strict Wat:

Loose Wat: the distance between water oxygen OW and the substrate Asn5 carbonyl carbon C was required to be ≤3.5 Å, and ∠OW–C–O was required to be 80°–100°, with the carbonyl carbon as the vertex. This initial nucleophilic-attack geometry was identical to Strict Nu.

Medium Wat: in addition to Loose Wat, both oxyanion-hole hydrogen bonds had to be present in the same frame.

Strict Wat: in addition to Medium Wat, the same water molecule satisfying the initial attack geometry had to form a hydrogen-bond contact with His171 ND1. The geometric criteria were OW···ND1 distance ≤3.5 Å and ∠ND1–OW–HW ≤30° for at least one water hydrogen HW, with the water oxygen as the angle vertex.

All thresholds were inclusive. The occupancy at each level was defined as the number of frames containing at least one water molecule satisfying that level divided by the total number of analyzed frames. A frame containing multiple qualifying water molecules was counted only once, and no minimum continuous residence time was imposed.

**Passing rates for hierarchical water screening**

Per-replicate frame occupancies under Loose Wat, Medium Wat, and Strict Wat were averaged across replicates to obtain group means. Two progressive screening passing rates were then defined:

Oxyanion-hole passing rate: the ratio of the group-mean Medium Wat occupancy to the group-mean Loose Wat occupancy, representing the fraction of water events satisfying initial nucleophilic-attack geometry that also passed the oxyanion-hole criterion.

His-geometry passing rate: the ratio of the group-mean Strict Wat occupancy to the group-mean Medium Wat occupancy, representing the fraction of water events passing the oxyanion-hole criterion that additionally satisfied the hydrogen-bond geometry between the same water molecule and His171 ND1.

All metrics were based on frame occupancies, with multiple qualifying water molecules in one frame counted once. When the number of analyzed frames was identical across replicates, the ratio of group-mean occupancies was equivalent to the ratio of pooled frame counts. Passing rates were calculated from unrounded values and reported separately from equally weighted averages of per-replicate conditional probabilities.

**Nucleophilic-attack geometry**

The backbone carbonyl carbon C and carbonyl oxygen O of substrate Asn5 defined the reaction center. The nucleophilic atom was the oxygen of water or the N-terminal nitrogen of incoming peptide Ser6. Attack geometry was defined by the distance d(Nu,C) between nucleophilic atom Nu and carbonyl carbon C and by ∠Nu–C–O with the carbonyl carbon as the vertex.

Two nucleophilic-attack geometry criteria were established:

Loose Nu: d(Nu,C)≤4.5 Å and ∠Nu–C–O between 70° and 110°.

Strict Nu: d(Nu,C)≤3.5 Å and ∠Nu–C–O between 80° and 100°.

All thresholds were inclusive. Neither criterion required prior formation of the oxyanion hole or a hydrogen-bond contact with His.

**Oxyanion-hole and hydrogen-bond states**

The oxyanion-hole geometric criterion was defined as the simultaneous acceptance of two hydrogen bonds by the backbone carbonyl oxygen of Asn5 from the backbone N–H groups of Cys213 and Gly172. Each hydrogen bond required an acceptor-O to donor-N distance ≤3.5 Å and ∠O–N–H ≤30° with the donor N as the vertex.

The formation probabilities of the Cys213 and Gly172 hydrogen bonds were calculated separately. Each frame was assigned to one of four mutually exclusive states: neither hydrogen bond present, only the Cys213 hydrogen bond present, only the Gly172 hydrogen bond present, or both hydrogen bonds present. The probability of each state was the corresponding frame count divided by the total number of analyzed frames. The probability of simultaneous formation of both hydrogen bonds was defined as the background oxyanion-hole probability.

**Conditional probability of oxyanion-hole formation**

Let H denote satisfaction of the oxyanion-hole geometric criterion, and C denote satisfaction of the corresponding nucleophile criterion. For each replicate trajectory, the background probability was defined as P(H)=N(H)/N, and the conditional probability as P(H|C)=N(H∩C)/N(C), where N is the total number of analyzed frames, N(H) is the number of frames satisfying the oxyanion-hole criterion, N(C) is the number of frames satisfying the nucleophile criterion, and N(H∩C) is the number satisfying both criteria.

The primary analysis used the Loose Nu criterion to calculate oxyanion-hole formation probabilities conditioned on peptide NAC or water NAC. A water-NAC frame was defined as one containing at least one water molecule satisfying the relevant distance and angle criteria. Frames containing multiple qualifying water molecules were counted once, and the conditional probability was calculated at the frame level.

A consistent rule was used for zero-condition frames in all conditional analyses. If no frame satisfied a condition, the conditional probability was recorded as undefined rather than zero; if condition frames existed but oxyanion-hole formation was never observed, the conditional probability was zero. Condition-frame counts and the number of valid replicates were reported for each group.

**Calculation of substrate-binding descriptors and oxyanion-hole states**

A common 0–500 ns window was analyzed for each of the three independent simulations of every system. Frames were sampled at 50 ps intervals, yielding 10,001 frames per trajectory. Enzyme residues were numbered according to the full-length sequences, and chains A and B denote the enzyme and substrate, respectively. The A:E211 OE1–B:N5 ND2 distance and the A:G173 Cα–B:S6 Cα distance were calculated for each frame; A:A173 was used instead of A:G173 for EchPAL1. The former distance described the position of the N5 side chain relative to the pocket base, whereas the latter described the relative position of the P1′-side peptide segment using B:S6 as a reference. These interatomic distances were not used alone to classify complete withdrawal of the side chain from the active-site pocket. Distances were calculated using the minimum-image convention for the periodic simulation box.

The two monitored oxyanion-hole hydrogen bonds were donated by the backbone N–H groups of A:C213 and A:G172 to the backbone carbonyl O of B:N5. An individual hydrogen bond was considered present when the O···N distance was ≤3.5 Å and the O–N–H angle was ≤30°, with the angle vertex at the donor N atom. No additional periodic-boundary correction was applied when evaluating hydrogen-bond geometry. Frames in which both hydrogen bonds were present were classified as the oxyanion-hole-formed state (“Both”), whereas frames in which neither bond was present were classified as the fully suppressed oxyanion-hole state (“Neither”) for this analysis. Here, “fully suppressed” refers specifically to the absence of both monitored hydrogen bonds and does not directly imply complete loss of catalytic activity. Frames containing only one of the two hydrogen bonds belonged to neither category; consequently, Both and Neither were mutually exclusive but not complementary.

Distance traces were displayed as means over non-overlapping 2 ns bins. Hydrogen-bond state strips were calculated using non-overlapping 1 ns bins, with occupancy defined as the number of frames assigned to a state divided by the total number of sampled frames in that bin. No preselection based on side-chain distance or other catalytic-geometry descriptors was applied. Bins were left-inclusive and right-exclusive, except that the final bin included the frame at 500 ns. Thus, regular 1 ns bins contained 20 frames and the final bin contained 21 frames. All occupancy strips used a common 0–100% color scale. Replicas were processed separately without averaging across simulations. Comparisons between the first and second halves of a trajectory used [0, 250) ns and [250, 500] ns, respectively. Mean distances and hydrogen-bond occupancies were calculated directly from the frames within each interval. These comparisons were descriptive and were not treated as tests of statistical significance.

**Sensitivity analysis of oxyanion-hole conditional probabilities**

To assess sensitivity of oxyanion-hole conditional probabilities to the definition of the nucleophile condition, Strict Nu was additionally applied to both water and incoming peptide, and a Proximity criterion was added for the incoming peptide, alongside the main Loose Nu analysis. Background P(H) and conditional P(H|C) probabilities were calculated identically for all criteria.

Proximity: a minimum heavy-atom distance strictly less than 5.0 Å between either Ser6 or Leu7 and the LAD2 residue, without restricting the attack angle. The LAD2 residues were Gly173 for EchAEP1 and Ala173 for EchPAL1.

In the supplementary sensitivity plots for EchAEP1, background and conditional probabilities were summarized using their respective valid replicates. In the paired EchPAL1 oxyanion-hole sensitivity plots, background and conditional probabilities were summarized using only complete paired replicates in which both probabilities were defined. Consequently, when some replicates had zero condition frames, the mean background probability in an EchPAL1 paired plot could differ from the background mean based on all three replicates.

**Geometric analysis of catalytic histidine hydrogen bonding**

Geometric conditions for donation of a hydrogen bond from the nucleophile to His171 ND1 were used to characterize local conformations potentially favorable for proton transfer. The donor D was the Ser6 N-terminal nitrogen or water oxygen. The required geometry was D···ND1 distance ≤3.5 Å and ∠ND1–D–H ≤30° for at least one hydrogen directly bonded to D, with donor D as the angle vertex. Peptide analysis included only hydrogens directly bonded to the N-terminal nitrogen and excluded the Ser side-chain hydrogen.

This geometric criterion was used as a proxy for His involvement in proton transfer and was not interpreted as an actual deprotonation event or a proton-transfer rate.

**Background and conditional probabilities for catalytic histidine**

Using the aforementioned His hydrogen-bond geometry, background probabilities were calculated separately for peptide and water:

Peptide background probability: the fraction of all analyzed frames in which the Ser6 N terminus satisfies the hydrogen-bond geometry with His171 ND1.

Water background probability: the fraction of all analyzed frames containing at least one solvent water molecule that satisfies the hydrogen-bond geometry with His171 ND1.

Neither background probability included a nucleophilic-attack geometry requirement.

Conditional probabilities were evaluated by screening with the corresponding Nu geometric criterion. The denominator was the number of frames satisfying Nu and the numerator was the number simultaneously satisfying Nu and the His hydrogen-bond criterion. For water, the same molecule had to satisfy both NAC and His hydrogen-bond criteria; contact of a different water molecule with His in the same frame could not substitute. Frames with multiple qualifying water molecules were counted once.

Neither the background nor conditional probability analysis required formation of the oxyanion hole.

**Residue fluctuation analysis**

After overall translational and rotational fitting, the root mean square fluctuation (RMSF) of each residue Cα atom relative to its time-averaged position was calculated independently for each replicate trajectory to quantify positional fluctuations. Between-group comparisons used chain A residues, comprising 278 sites for EchAEP1 and 273 for EchPAL1. No additional fit to the mean structure was performed before RMSF calculation.

RMSF was calculated independently within each replicate and then summarized using replicates as statistical units. Group means were equally weighted means across valid replicates, and error bars show the standard error of the replicate means (sample standard deviation divided by the square root of the number of valid replicates). Per-residue RMSF values from individual replicates were compared using two-sided Welch t tests, and fold changes (FC) were calculated as the ratio of mean RMSF in the comparison group to that in the reference group. Exploratory candidate sites were defined by unadjusted p<0.05 and |log₂FC|>1; Benjamini–Hochberg-adjusted q values were also reported. Meeting these exploratory criteria did not imply significance after multiple-testing correction.

**Statistical analysis of probabilities and frame occupancies**

Probabilities and frame occupancies were first calculated for each replicate separately and then summarized with replicates as statistical units. Group means were equally weighted averages of valid replicates, and error bars represent the standard error among replicate means (sample standard deviation divided by the square root of the number of valid replicates). Trajectory frames were not treated as independent statistical replicates.

For combined plots of the three oxyanion-hole probabilities, pairwise comparisons among background, peptide-conditioned, and water-conditioned probabilities within a group used two-sided paired t tests; between-group comparisons of the same metric used two-sided Welch t tests. Nine comparisons within each plot were jointly adjusted by the Holm–Bonferroni method. Stars in plots correspond to unadjusted p values, with adjusted p values reported separately. Water-frame occupancy and hydrogen-bond state plots were summarized descriptively.

For His-geometric probability analyses, each group was tested using two-sided paired t tests for water-conditioned versus peptide-conditioned probabilities, water background versus water-conditioned probabilities, and peptide background versus peptide-conditioned probabilities. Between-group water-conditioned and peptide-conditioned probabilities were each assessed using two-sided Welch t tests. For each geometric criterion, the four conditional-probability comparisons and the four within-category background-versus-conditioned comparisons were adjusted separately using the Holm method. Stars in plots indicate unadjusted two-sided p values; adjusted p values are reported separately. Paired tests required at least three complete valid pairs, and Welch tests required at least three valid replicates in each group.

**Supplementary Figures**

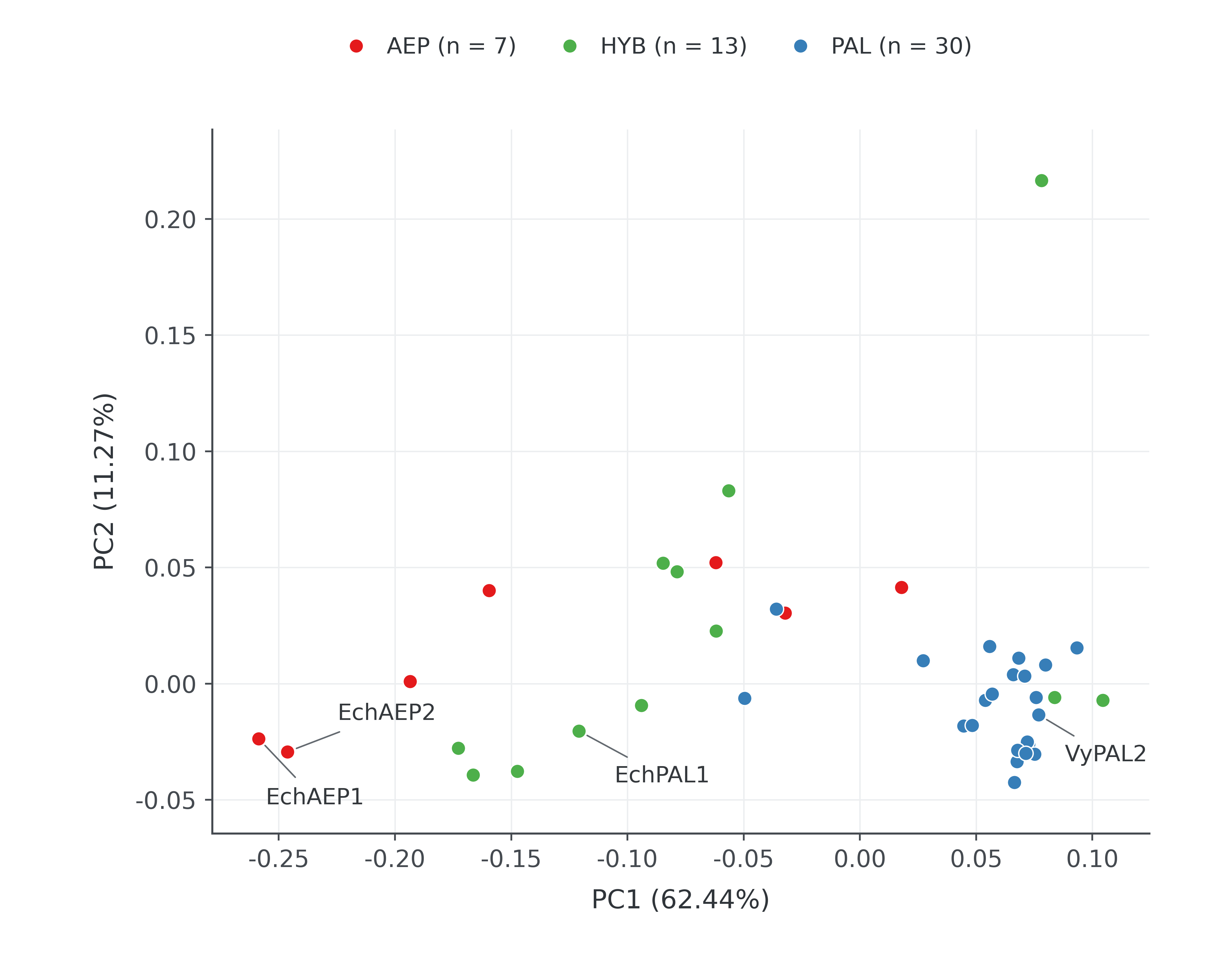

Figure S1. Distribution of the natural enzyme dataset in sequence space. Four naturally occurring enzymes from Exallage were labeled in the plot. Because discovery and characterization of natural enzymes require considerable time, the natural enzyme dataset used for the machine-learning sliding-window analysis comprised 49 enzymes and did not include EchPAL1. To maintain continuity with the main text, EchPAL1 was added to this figure, bringing the displayed total to 50 enzymes.

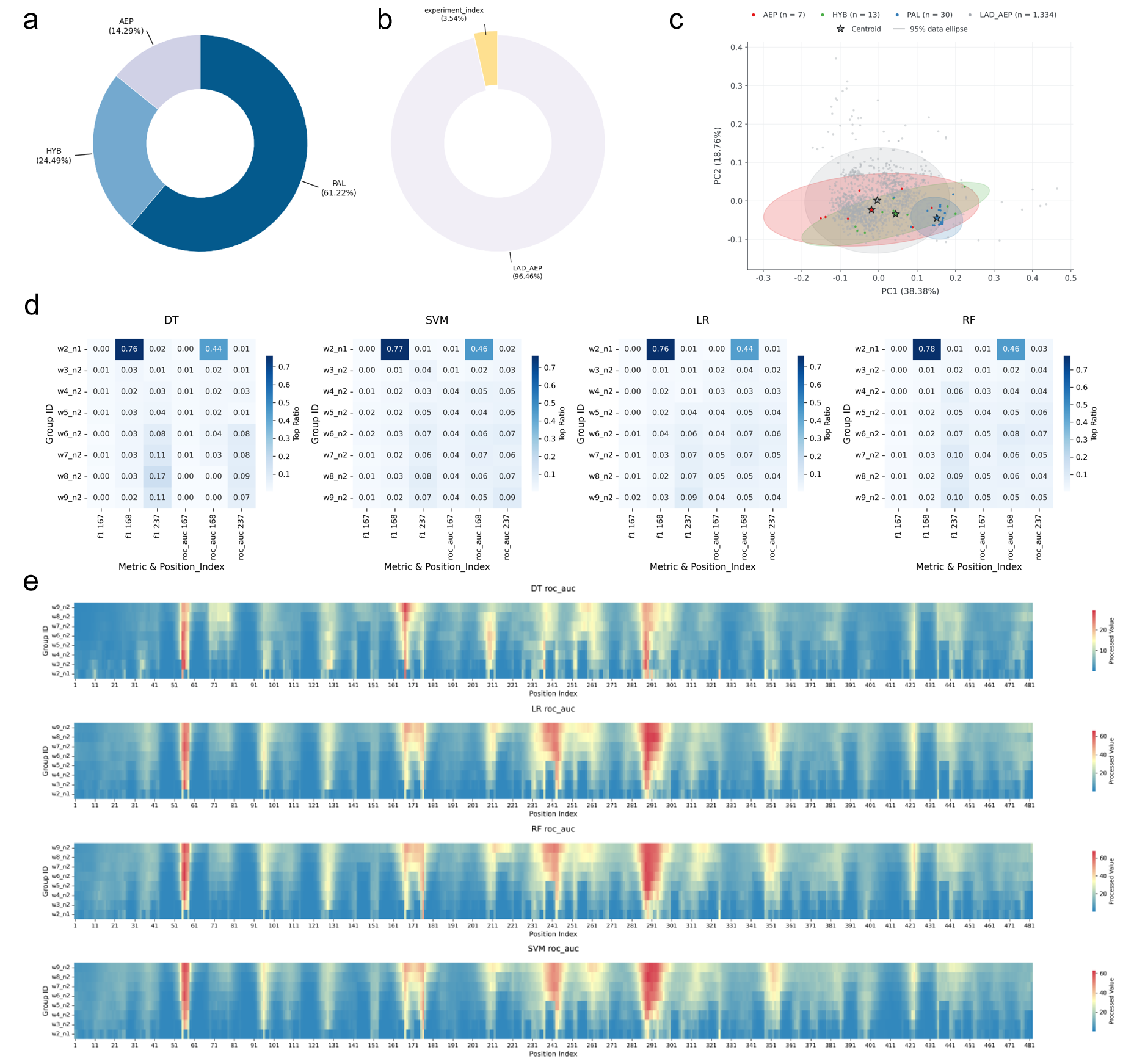

Figure S2. Machine-learning sliding-window screening based on a natural legumain dataset. (a) Proportions of PAL, HYB, and AEP in the experimentally labeled dataset; these labels were assigned according to experimental measurements. (b) Proportions of experimentally labeled and LAD_AEP records; LAD-based labels were inferred from the LAD hypothesis and were not experimentally validated. (c) Distribution of the natural enzyme dataset in protein sequence space. Circles represent individual sequences, stars mark the centroid of each category, and translucent colored regions show nominal 95% ellipses. (d) Rankings of position importance at LAD sites. Each window size was paired with a specified best-performing n-gram length, with F1 or ROC-AUC as the evaluation metric. (e) Cross-validation-based position classification-importance analysis. Each window size was paired with a specified best-performing n-gram length, and position classification importance reflects the association of a site with reaction directionality.

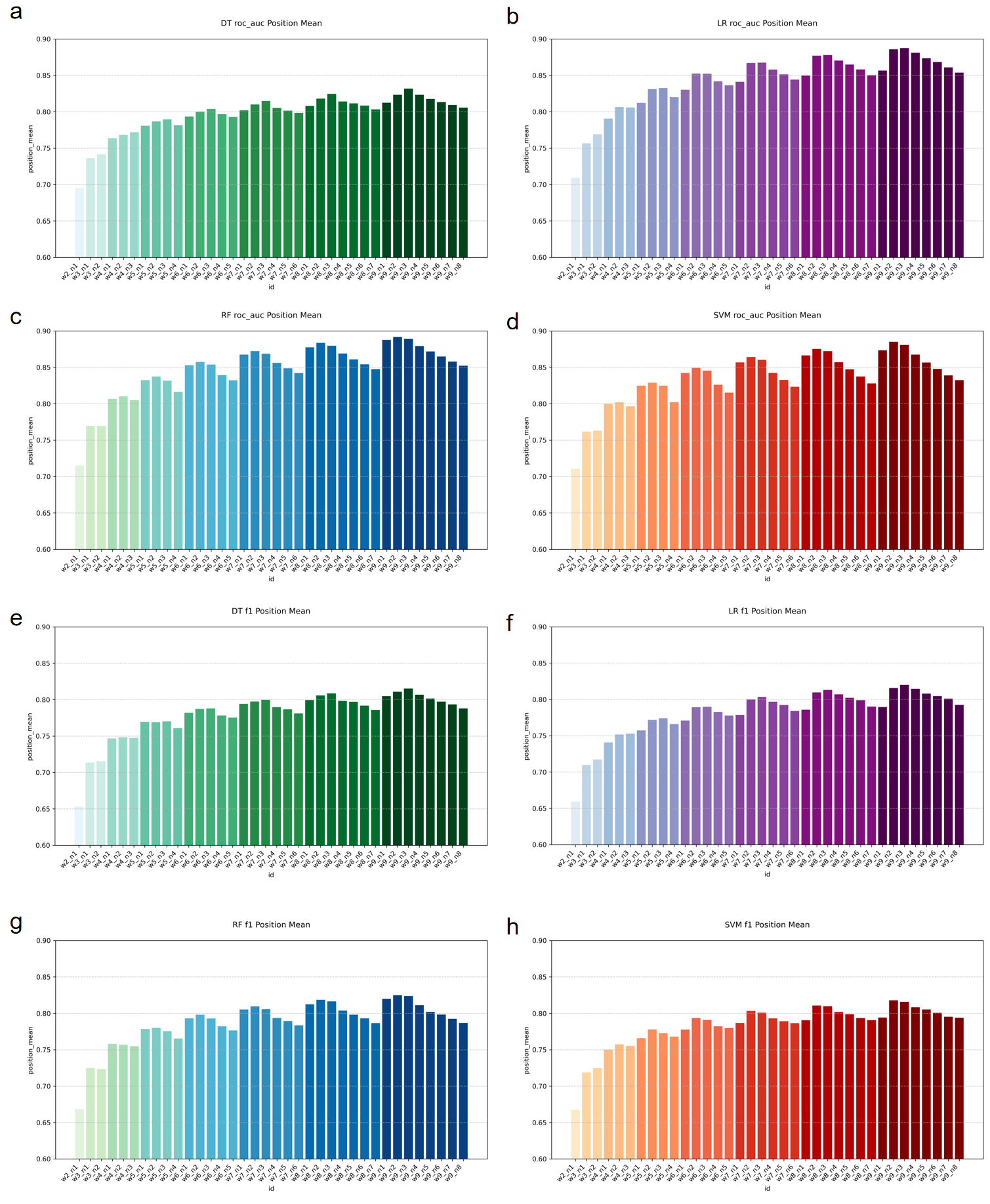

Figure S3. Grid-search results for sliding-window size and bag-of-words n-gram length. (a–d) Grid searches evaluated using ROC-AUC for decision tree (DT), logistic regression (LR), random forest (RF), and support vector machine (SVM), respectively. (e–h) Grid searches evaluated using F1 for DT, LR, RF, and SVM, respectively.

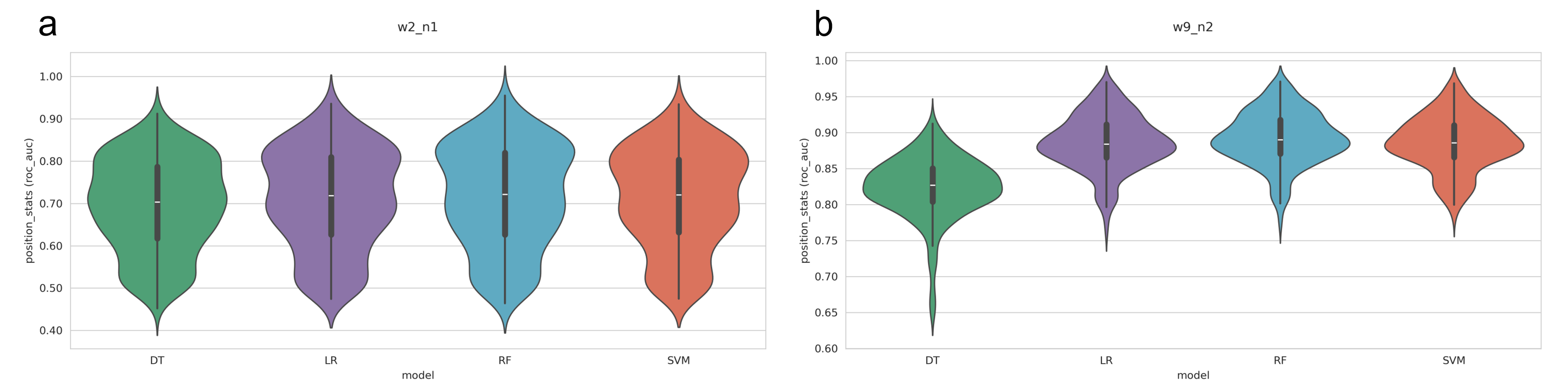

Figure S4. Distribution of position-level classification importance based on cross-validation. (a) Position importance with a window size of 2 and an n-gram length of 1. (b) Position importance with a window size of 9 and an n-gram length of 2.

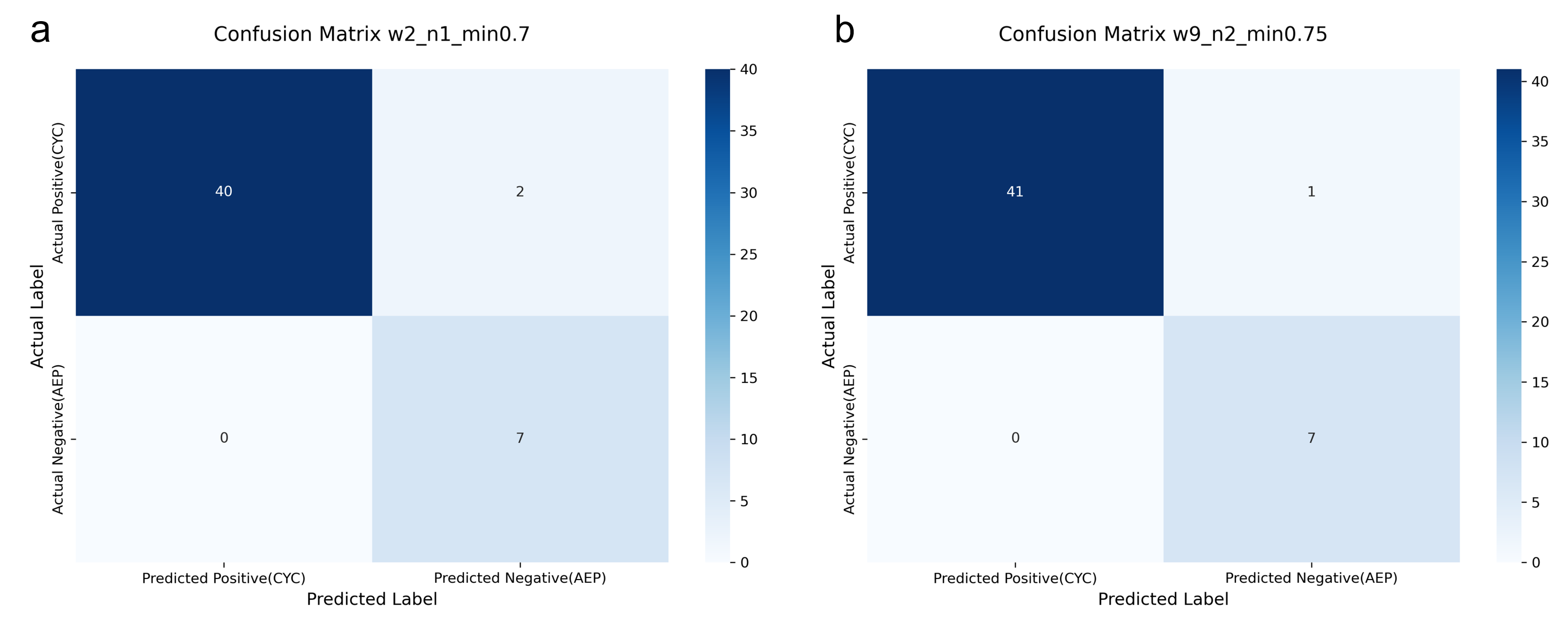

Figure S5. Confusion matrices for binary predictions by the ensemble model. (a) Confusion matrix for the ensemble model constructed with window size w2 and a minimum submodel-score threshold of 0.7. (b) Confusion matrix for the ensemble model constructed with window size w2 and a minimum submodel-score threshold of 0.75.

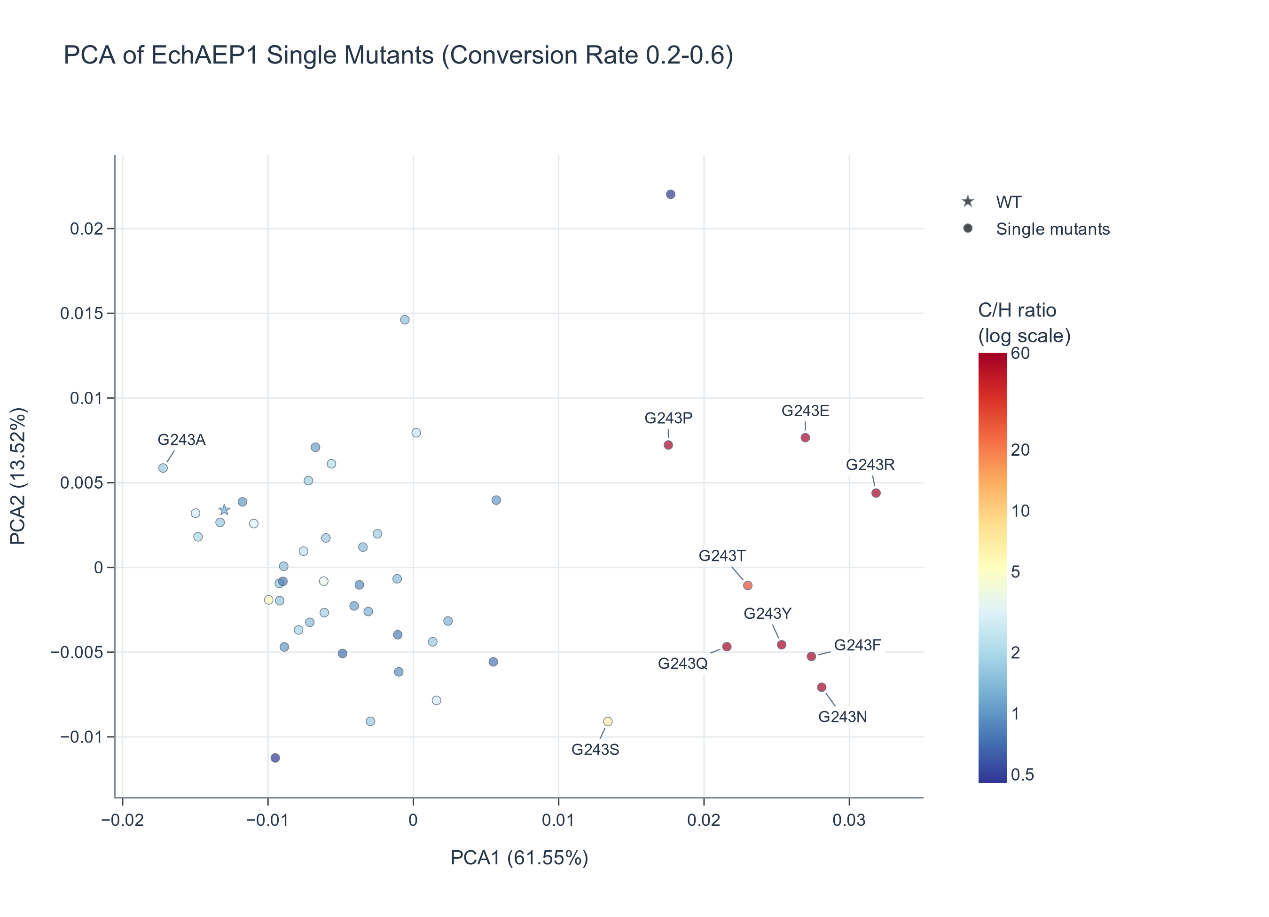

Figure S6. Sequence-space distribution of EchAEP1 and round-0 single mutants. Points were colored according to cyclization assay results. Seven single substitutions at position 243 (243 Q/N/P/Y/F/E/R) produced virtually no hydrolysis product; their ch_ratio values were therefore manually assigned a value of 60.

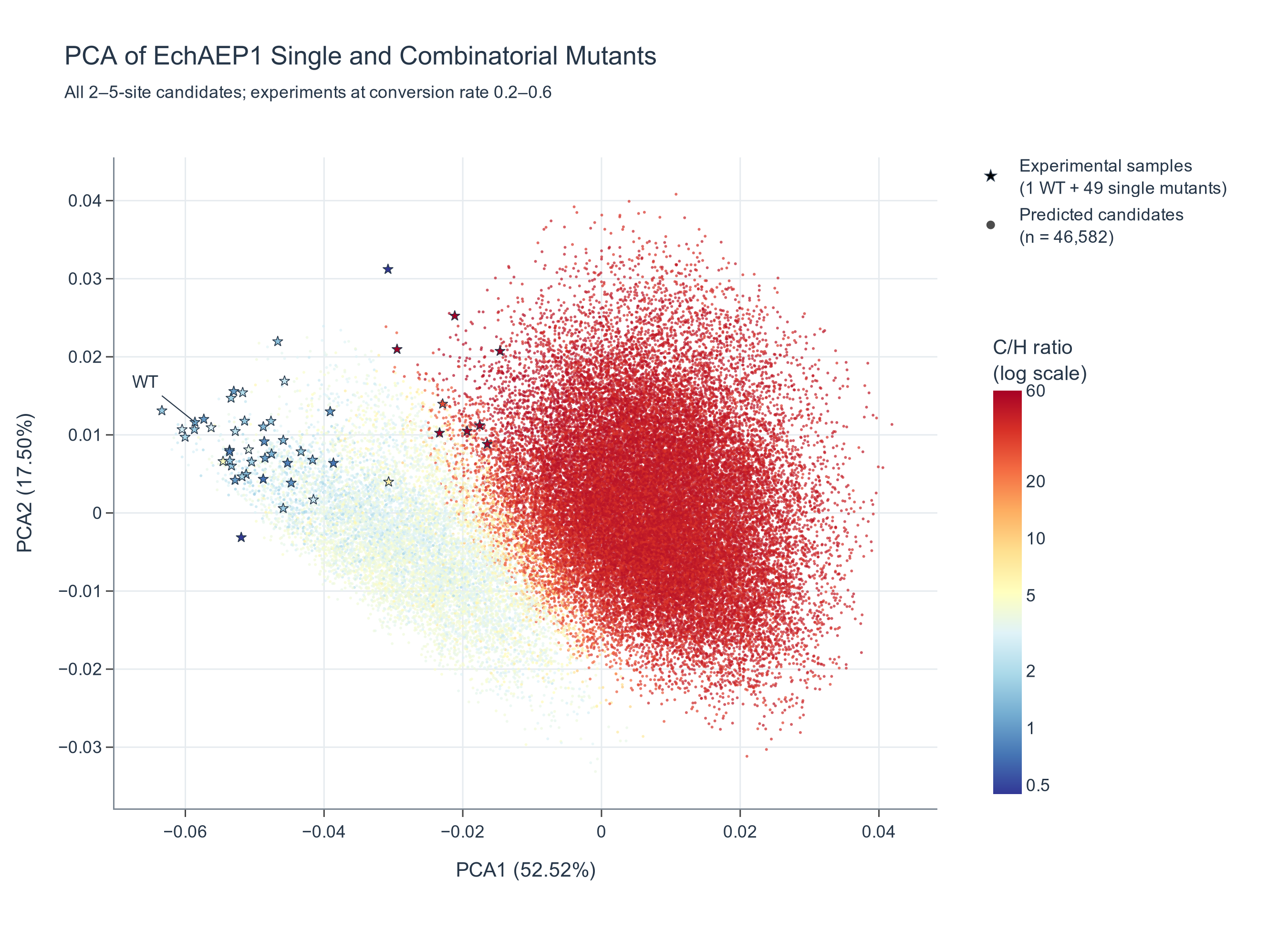

Figure S7. Sequence-space distribution of EchAEP1, round-0 single mutants, and all possible 2–5-site combinations of positive single mutants.

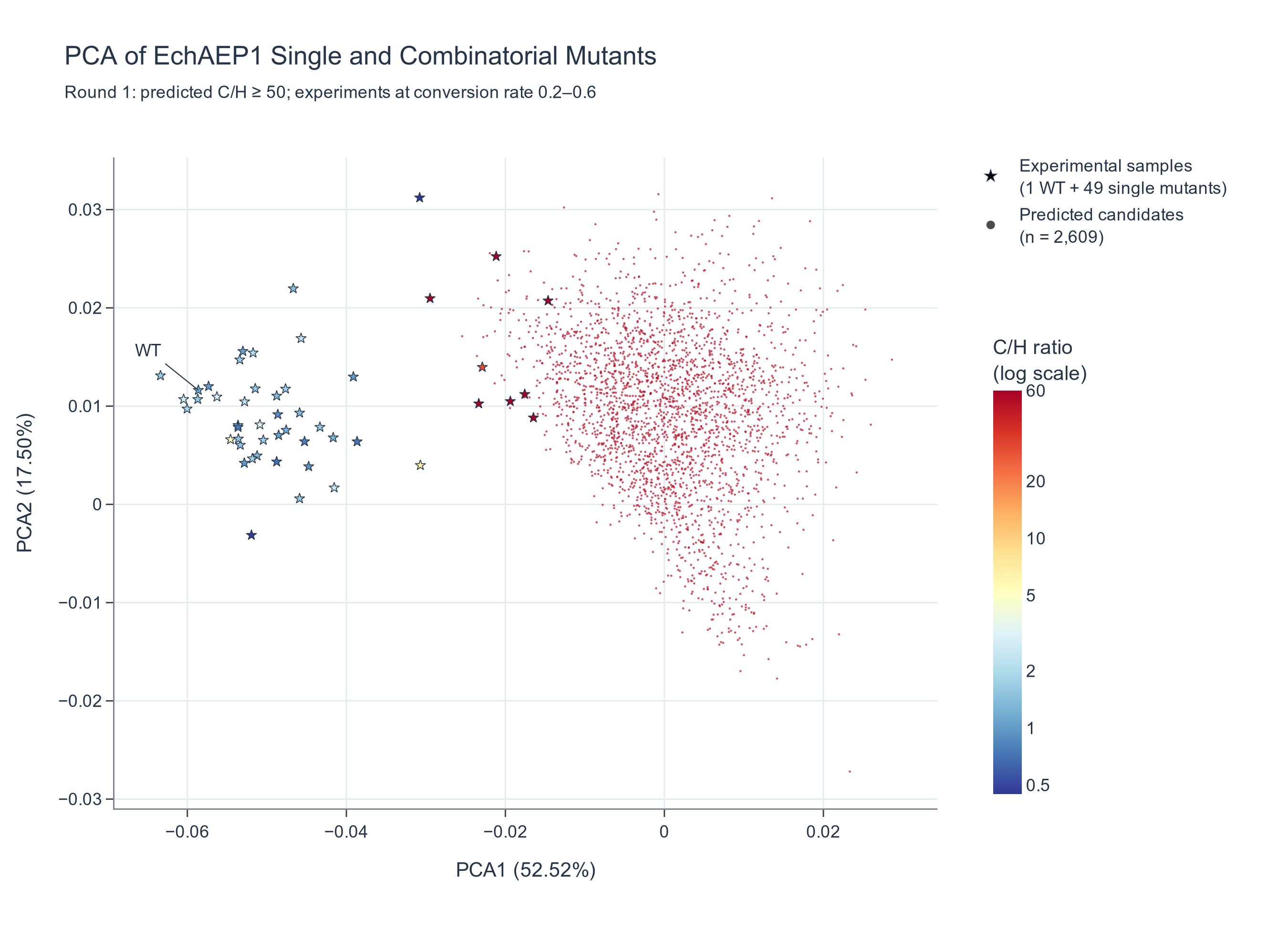

Figure S8. Sequence-space distribution of EchAEP1, round-0 single mutants, and high-prediction combinatorial mutants. Combinations with predicted ch_ratio≥50 were classified as high-prediction candidates.

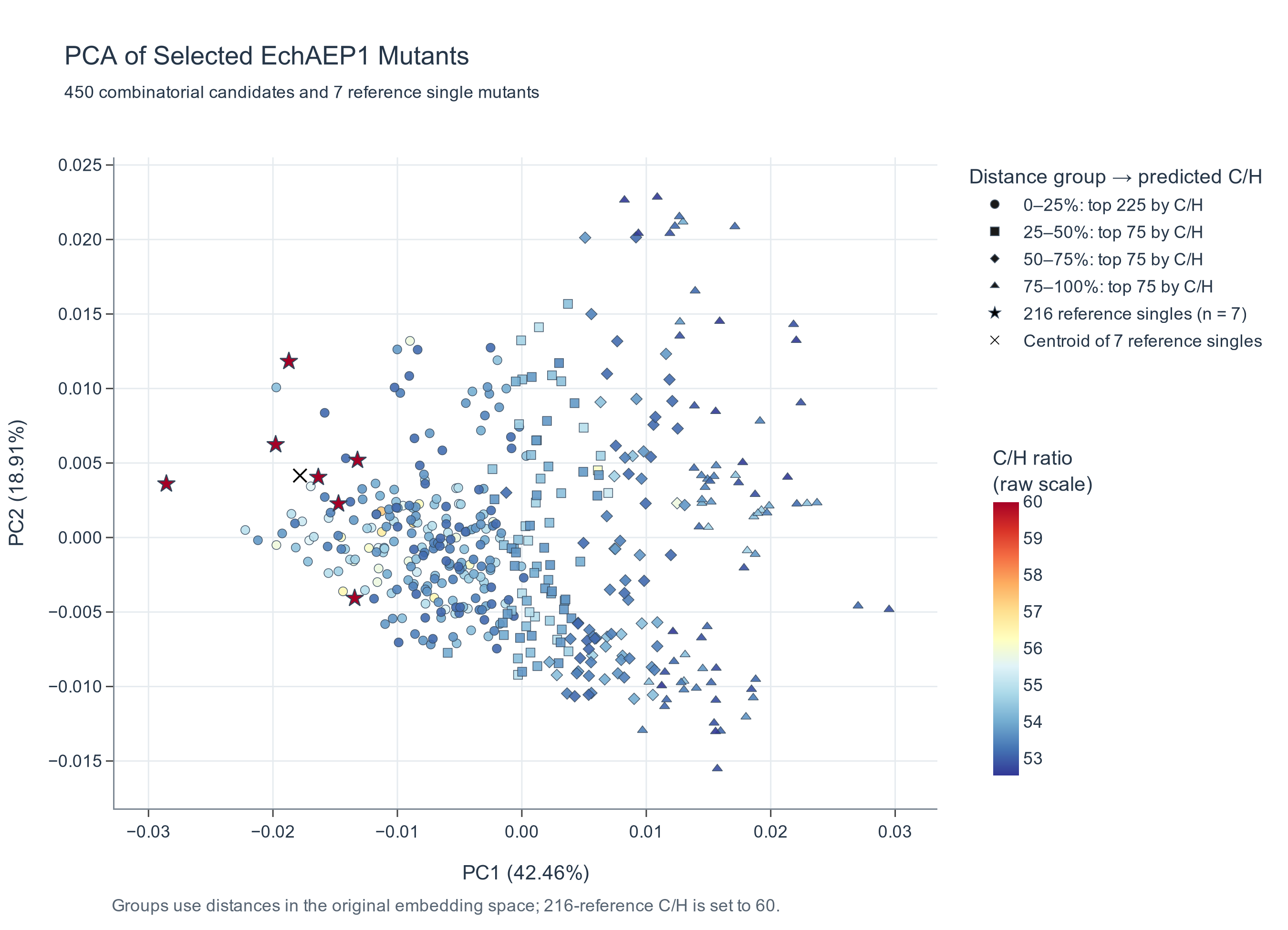

Figure S9. Sequence-space distribution of seven position-243 single mutants showing no detectable hydrolysis during cyclization and round-1 combinatorial candidates requiring ddg calculations. Cyclization data for candidate combinations are predictions.

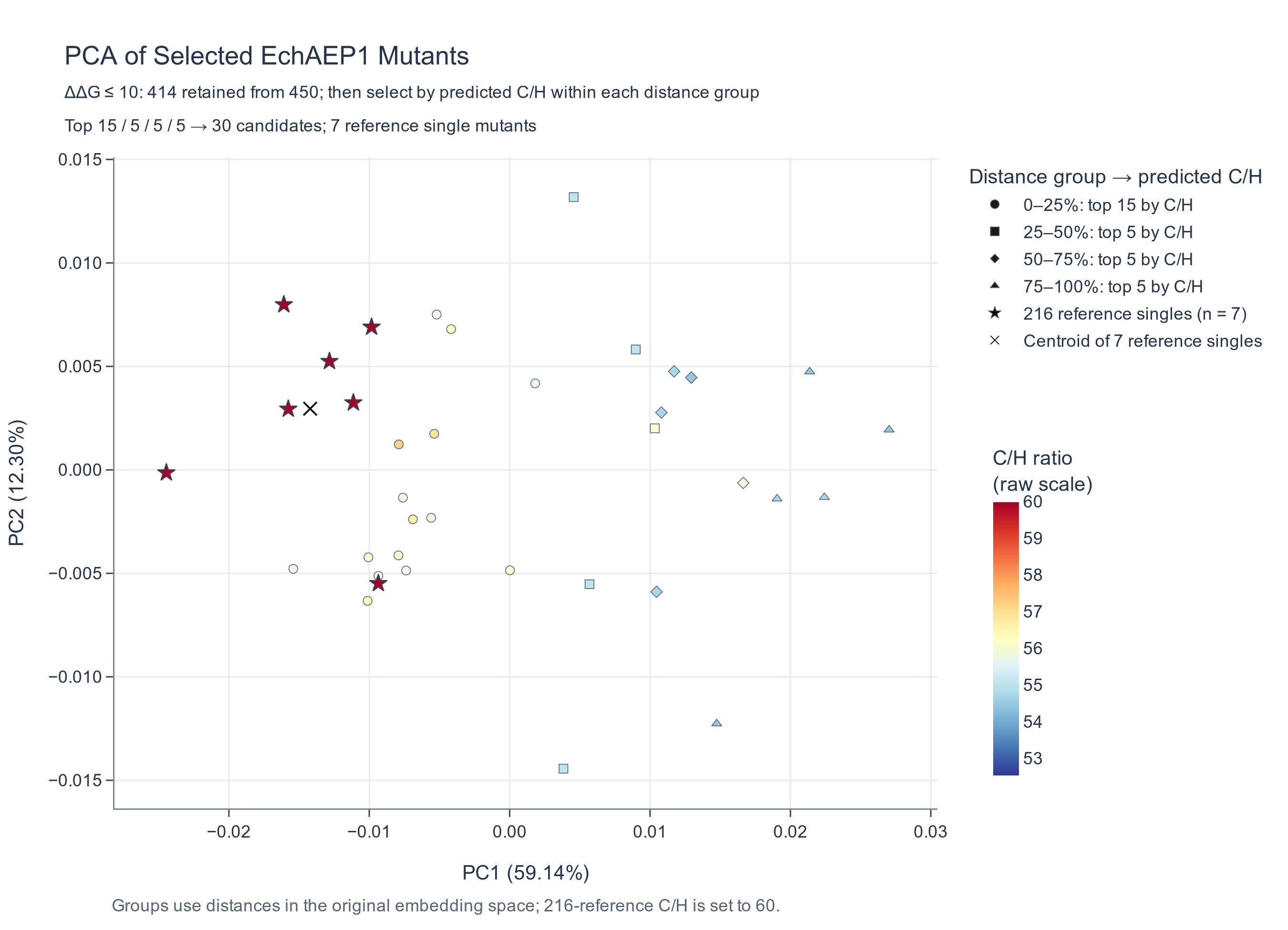

Figure S10. Sequence-space distribution of seven position-243 single mutants showing no detectable hydrolysis during cyclization and round-1 combinatorial candidates selected for experimental validation. Cyclization data for candidate combinations are predictions.

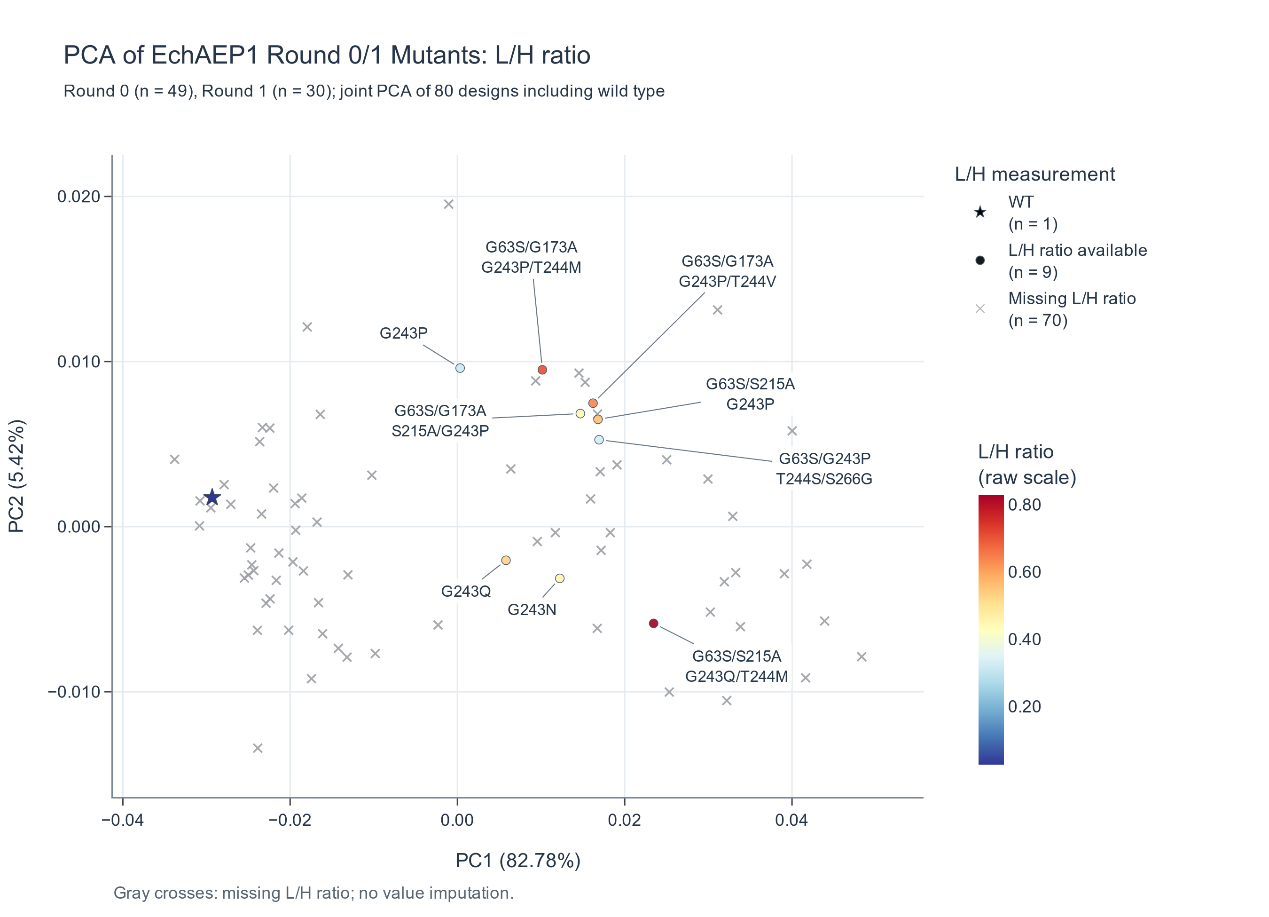

Figure S11. Sequence-space distribution of EchAEP1, round-0 single mutants, and round-1 combinatorial mutants. Points with available ligation assay data were colored accordingly.

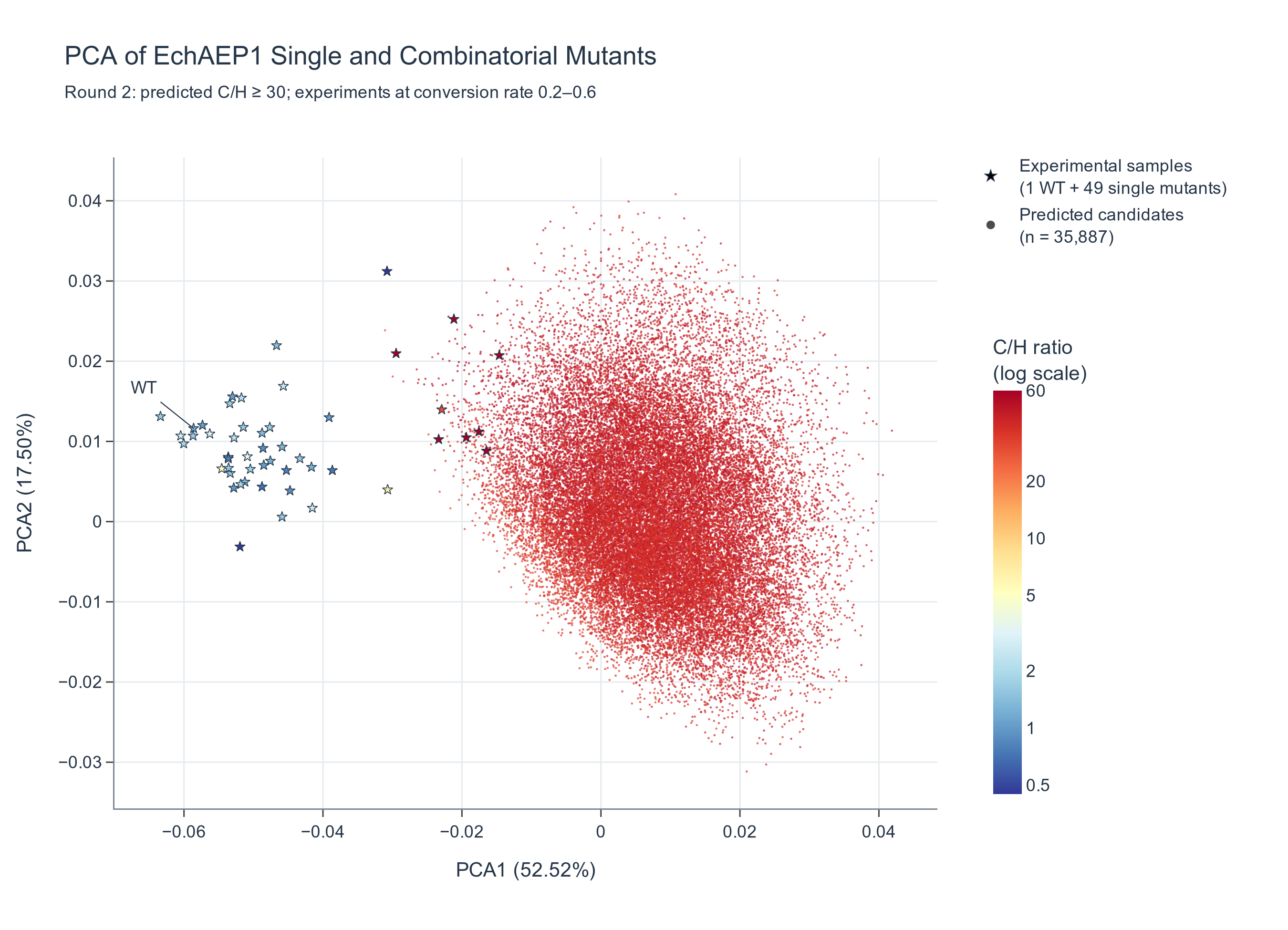

Figure S12. Sequence-space distribution of EchAEP1, round-0 single mutants, and medium-to-high-prediction combinatorial mutants. Combinations with predicted ch_ratio≥30 were classified as medium-to-high-prediction candidates.

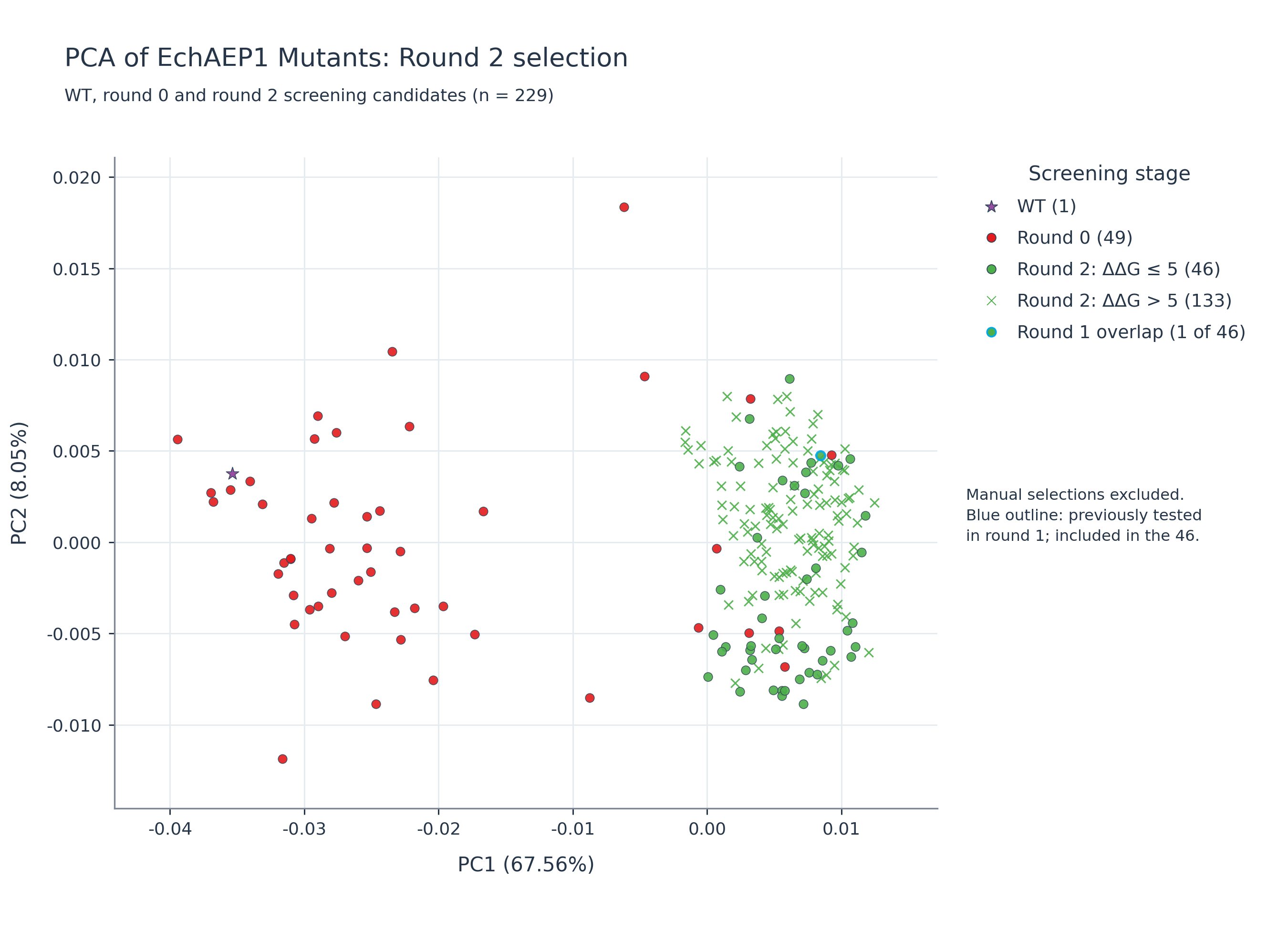

Figure S13. Sequence-space distribution of EchAEP1, round-0 single mutants, and round-2 combinatorial candidates selected by distance.

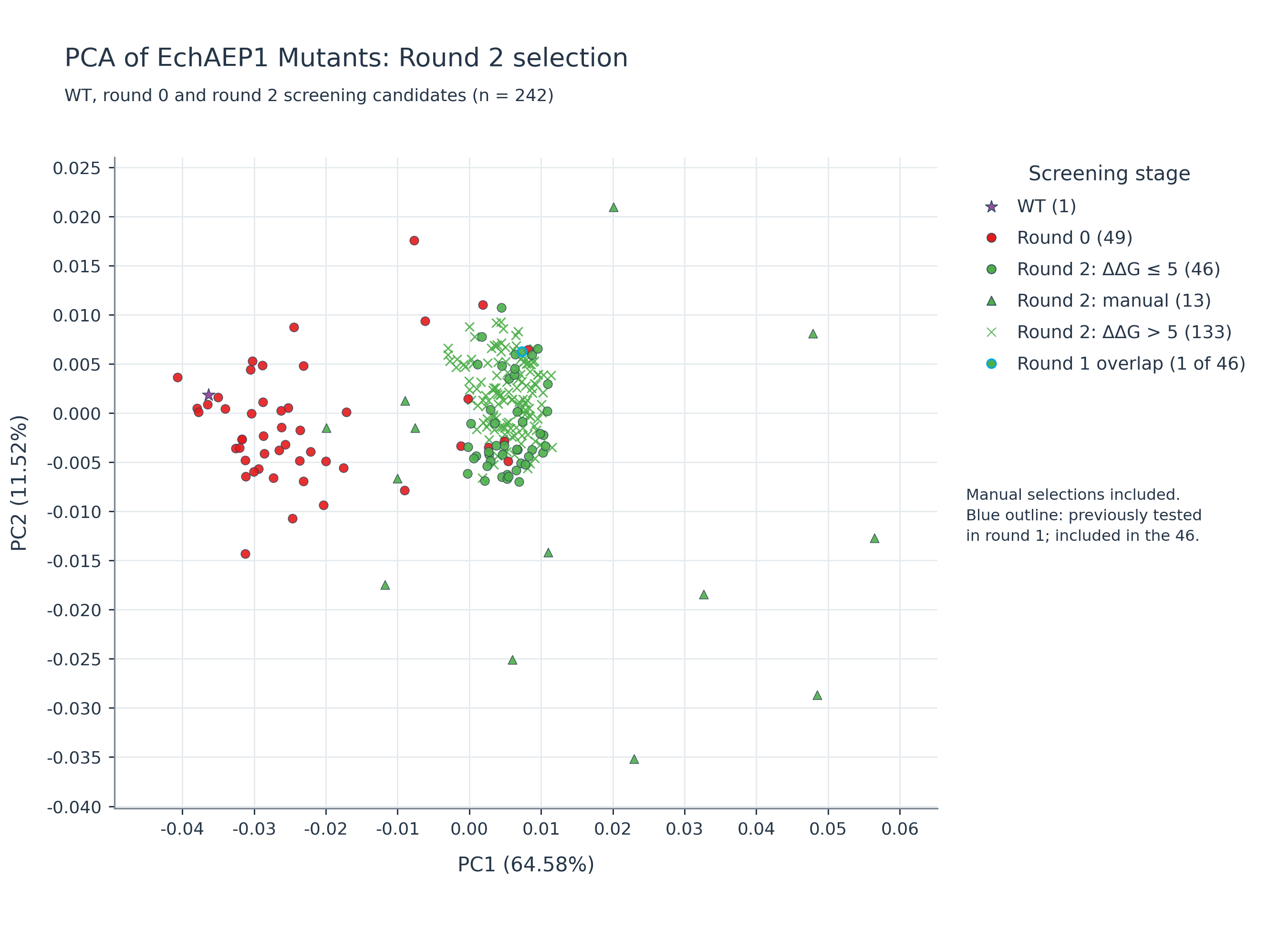

Figure S14. Sequence-space distribution of EchAEP1, round-0 single mutants, and round-2 combinatorial candidates selected by distance or manually.

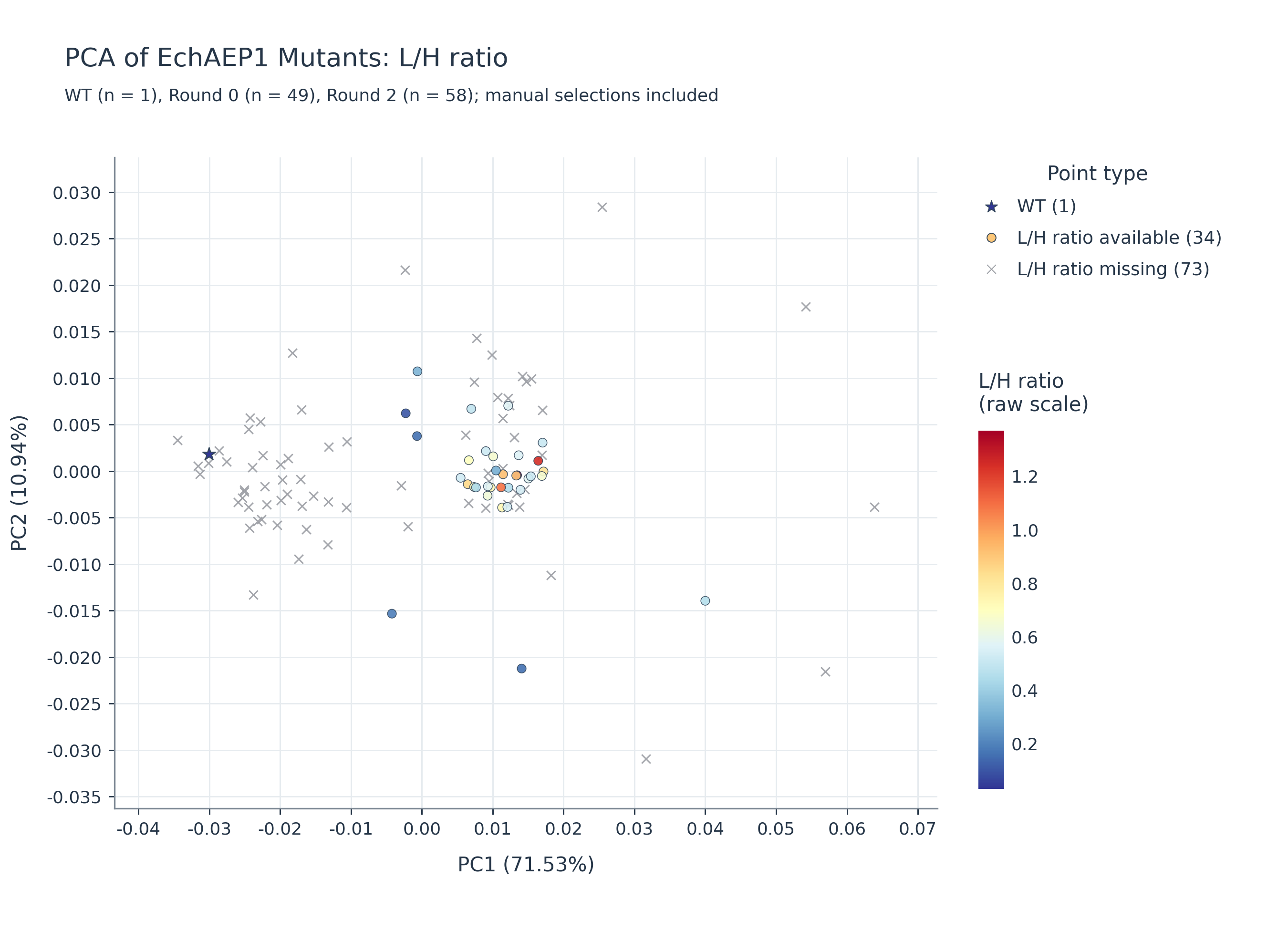

Figure S15. Sequence-space distribution of EchAEP1, round-0 single mutants, and round-2 combinatorial mutants. Points with available ligation assay data were colored accordingly.

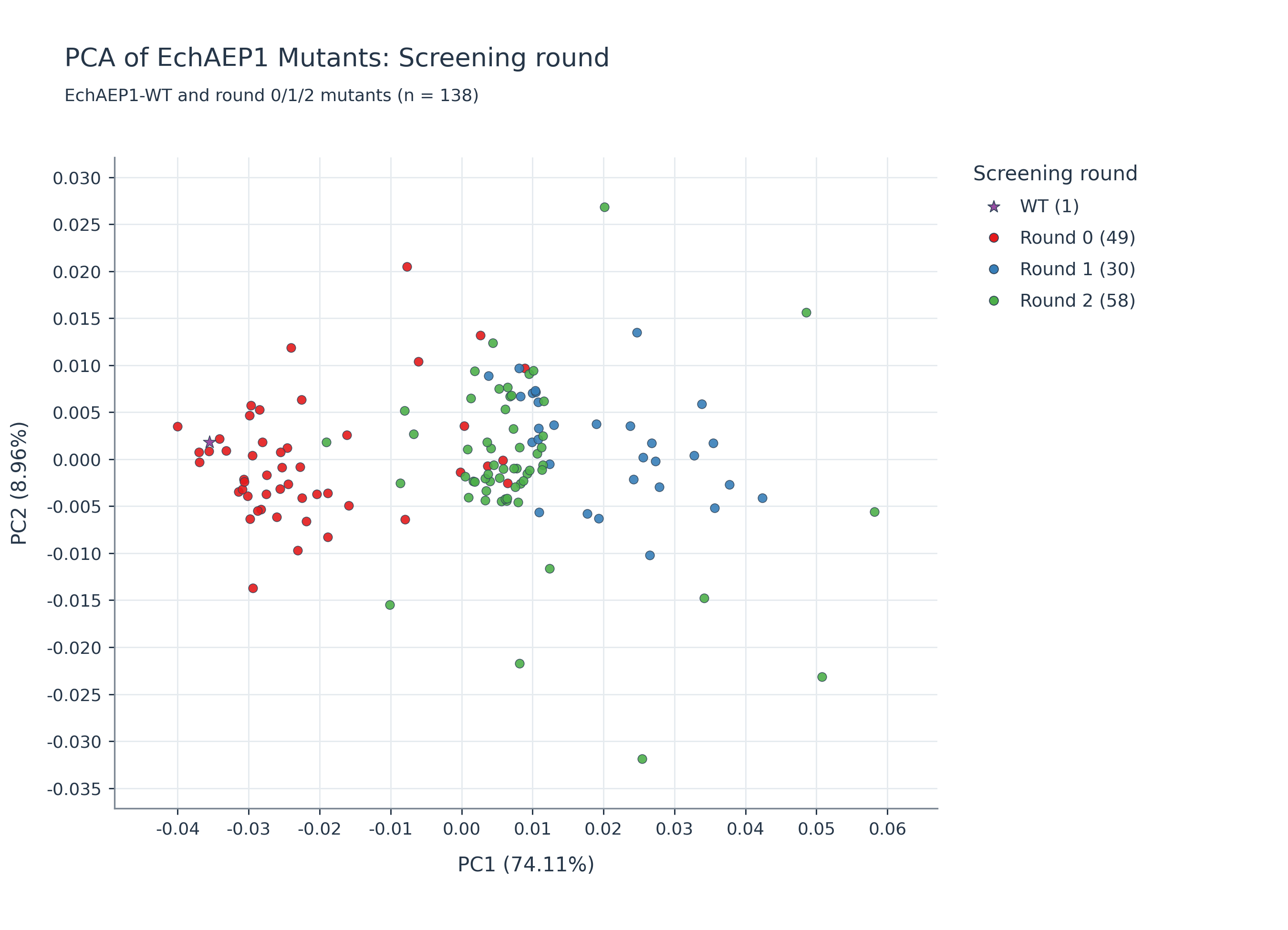

Figure S16. Sequence-space distribution of EchAEP1 and mutants from rounds 0, 1, and 2.

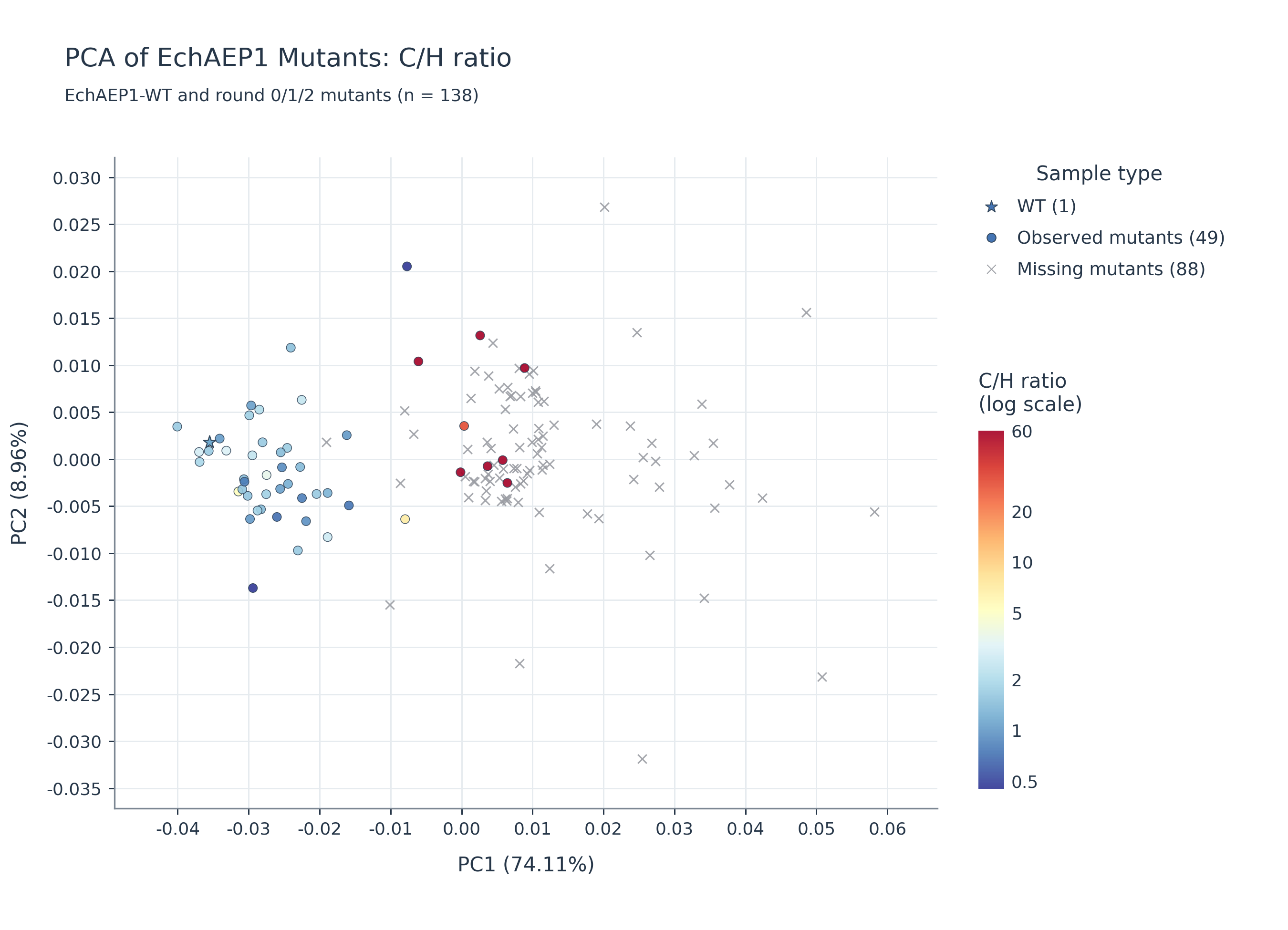

Figure S17. Sequence-space distribution of EchAEP1 and mutants from rounds 0, 1, and 2, with corresponding cyclization activity labels.

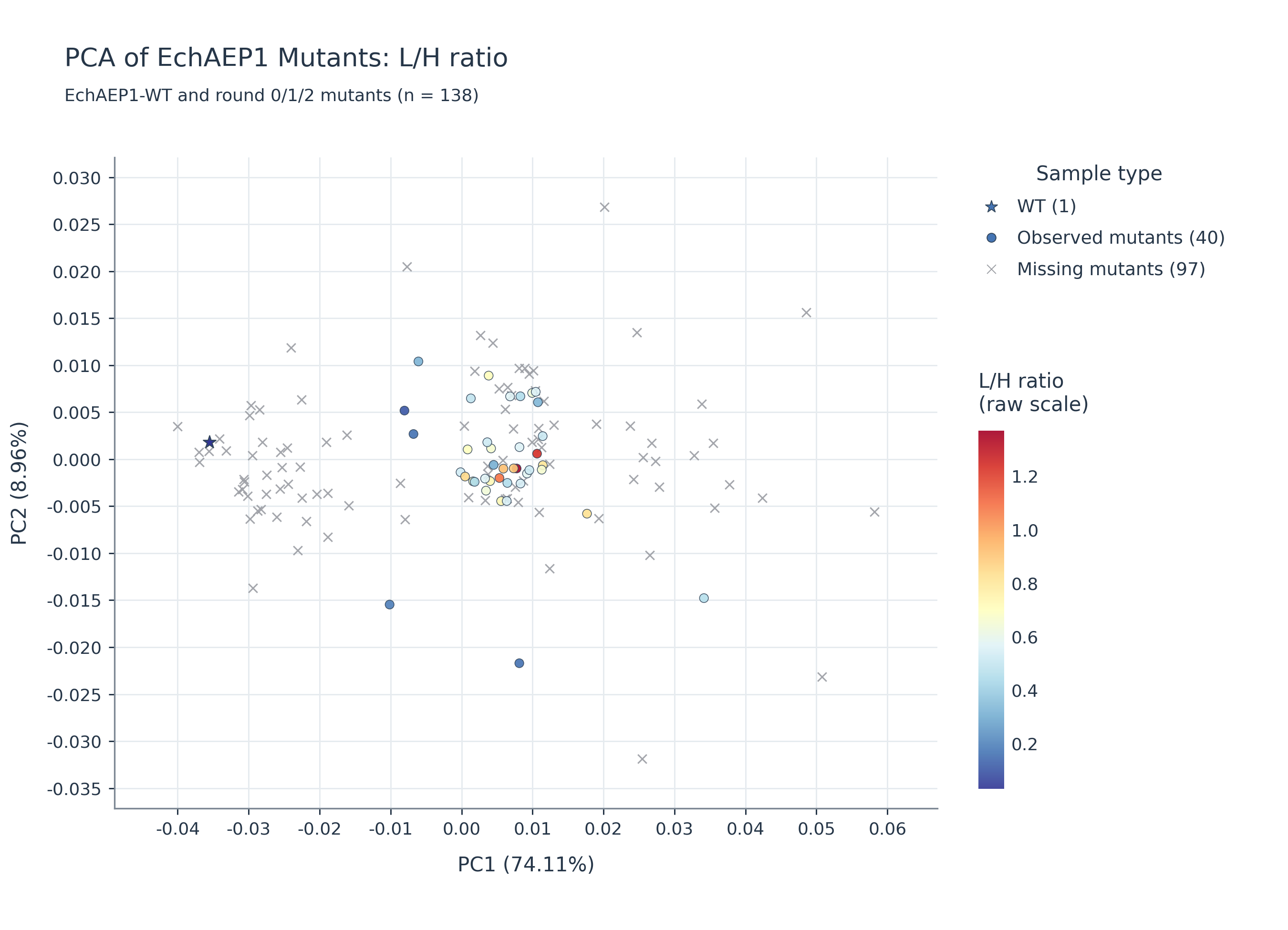

Figure S18. Sequence-space distribution of EchAEP1 and mutants from rounds 0, 1, and 2, with corresponding ligation activity labels.

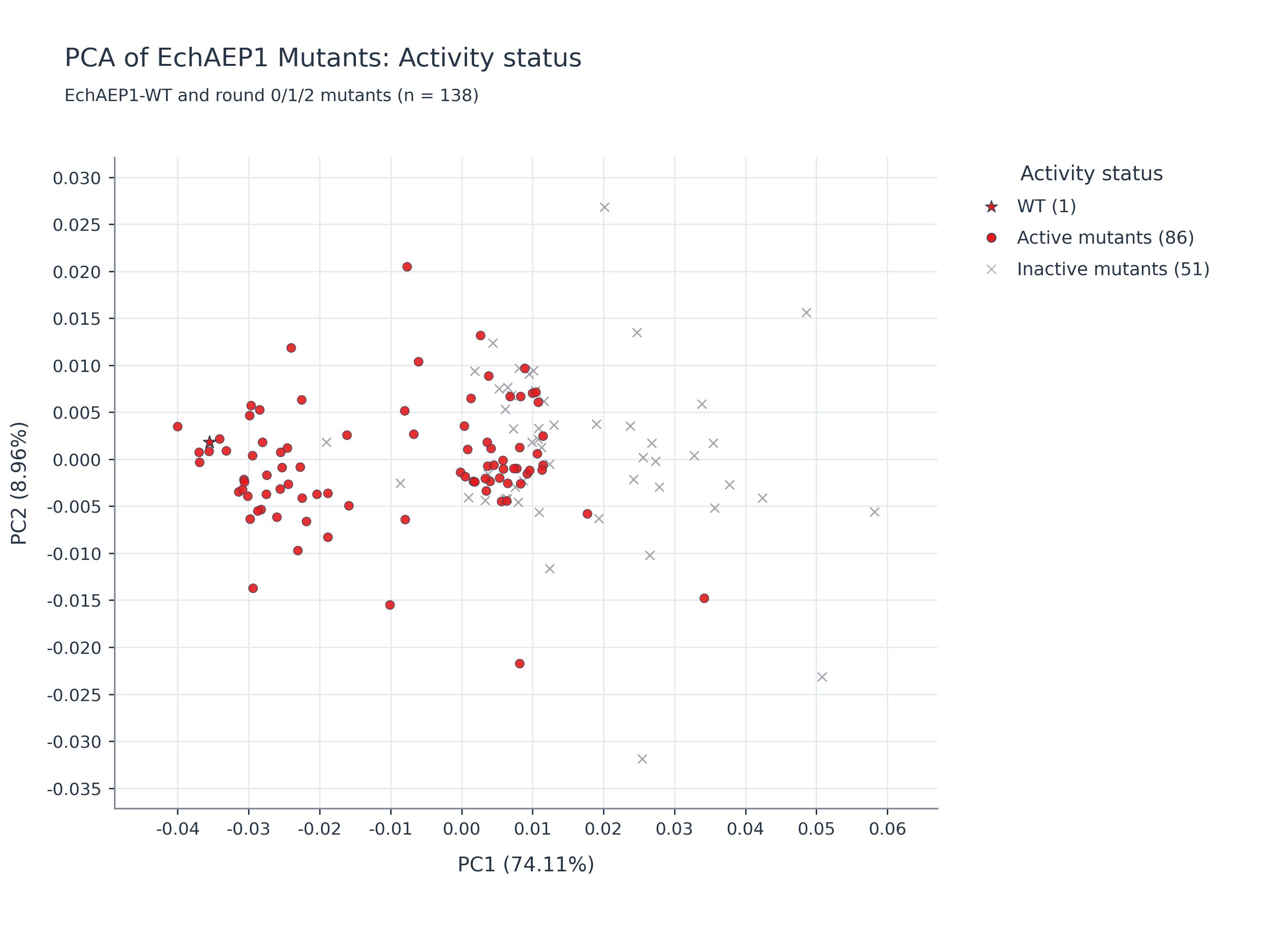

Figure S19. Sequence-space distribution of EchAEP1 and mutants from rounds 0, 1, and 2, with corresponding activity-status labels. After multiple rounds of assays, mutants lacking both cyclization and ligation activity data were designated inactive; all others were designated active.

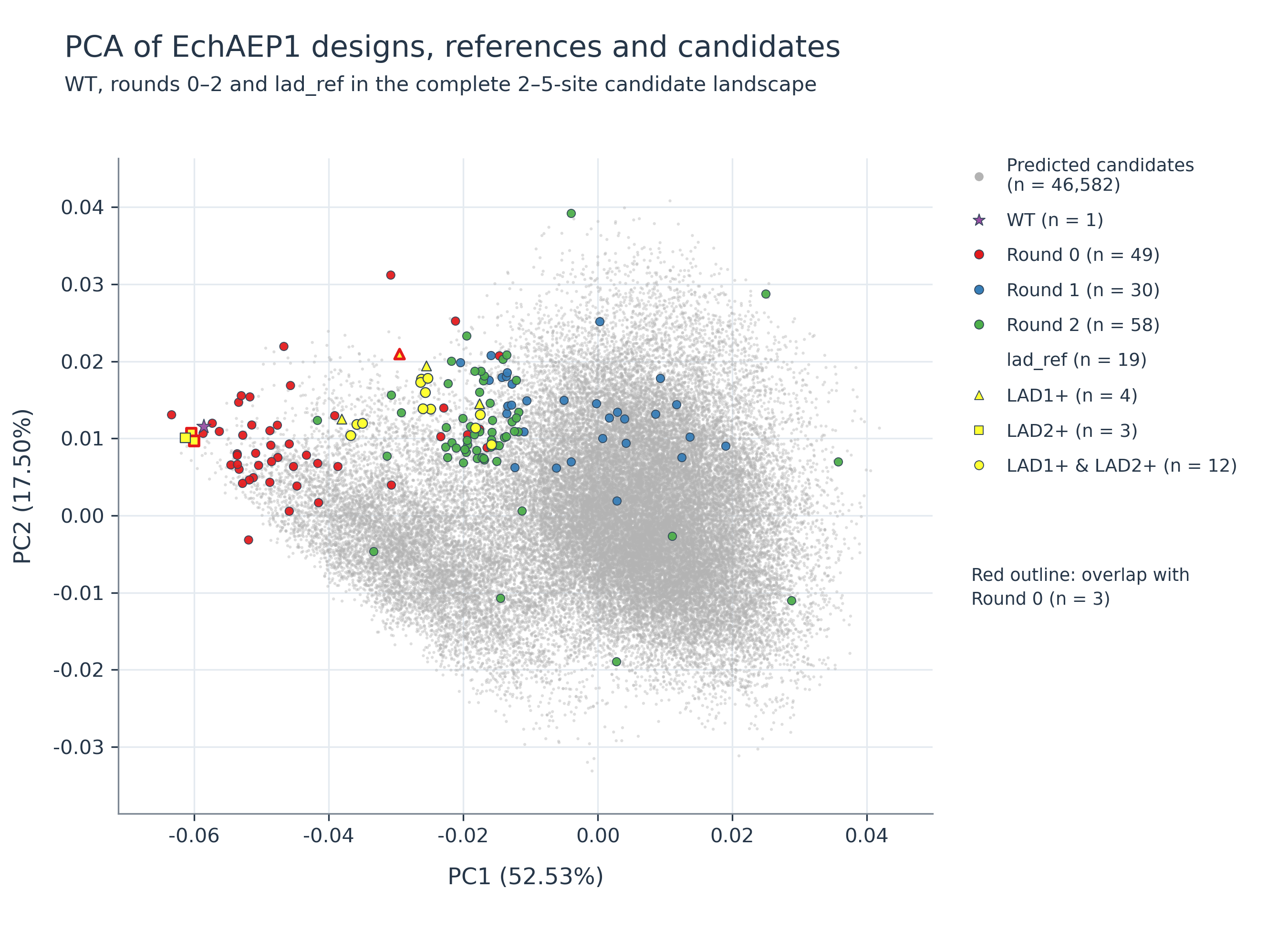

Figure S20. Sequence-space distribution of EchAEP1, mutants from rounds 0, 1, and 2, all possible 2–5-site combinations of positive single mutants, and LAD-based mutants.

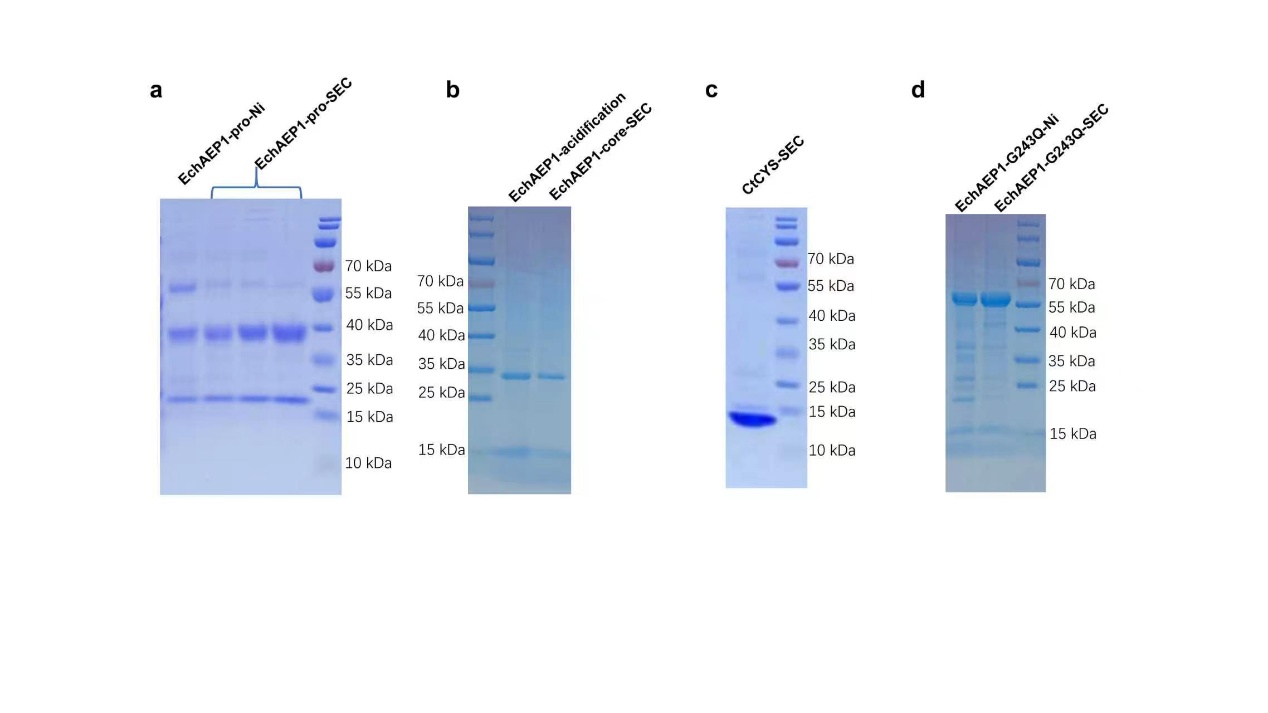

Figure S21. Expression and purification of EchAEP1, EchAEP1-G243Q, and CtCYS. (a) EchAEP1 proenzyme following Ni-NTA affinity chromatography and size-exclusion chromatography (SEC). (b) Acid-treated EchAEP1 and catalytic core domain purified by SEC. (c) CtCYS purified by SEC. (d) EchAEP1-G243Q after Ni-NTA affinity chromatography and SEC. Protein molecular-weight standards (kDa) are indicated at the sides.

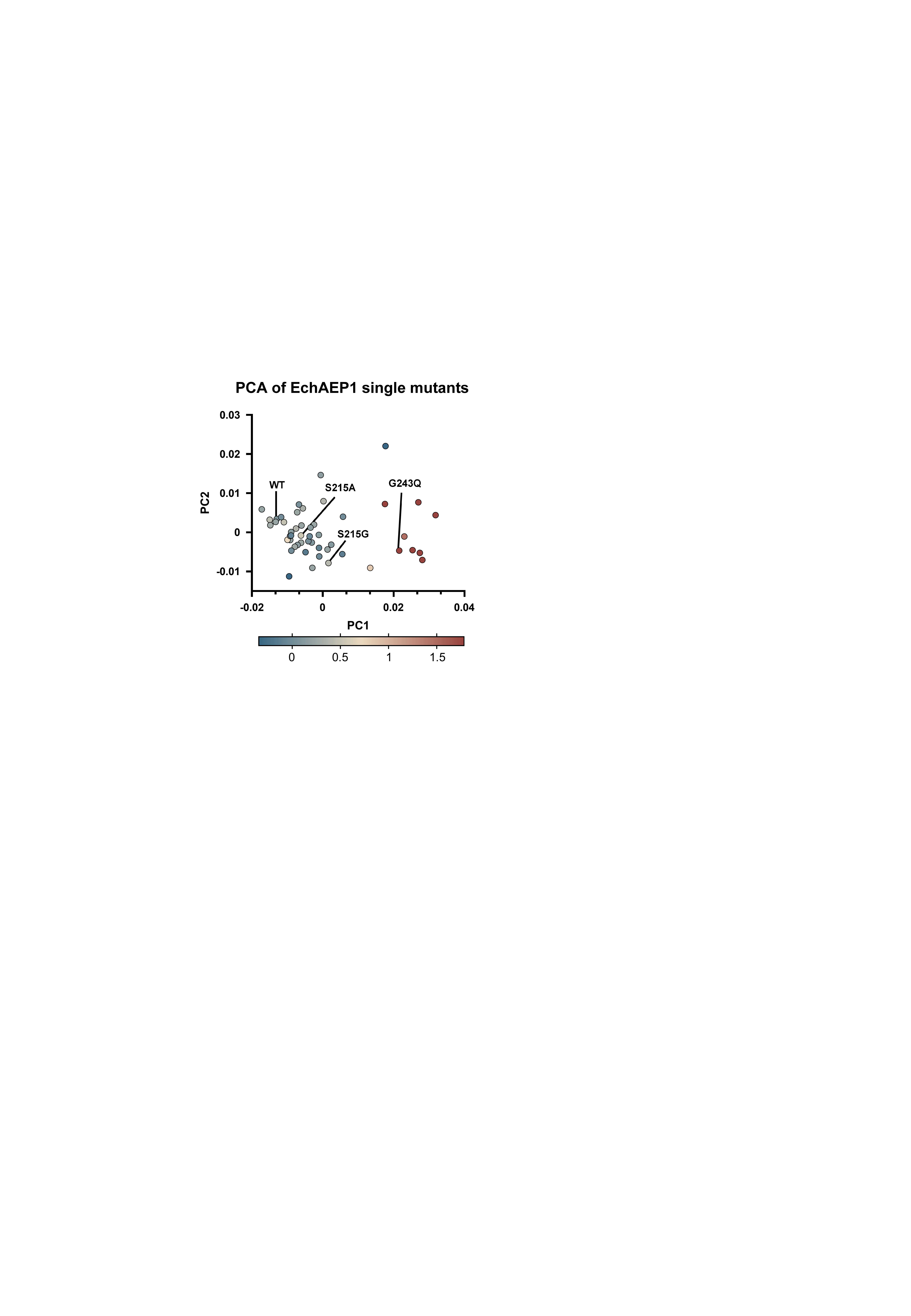

Figure S22. Distribution of round-0 single-substitution variants in sequence space. Each dot represents an individual single-substitution variant, colored according to its corresponding cyclization activity (ch_ratio). The color gradient indicates the activity range from low (blue) to high (red).

**Supplementary Tables**

**Table S1** LAD Characteristics and Cyclization Activities of 36 Positive Single Mutants.

| mut_infor | round_num | lad_ref_type | C_H_ratio | position |
| --- | --- | --- | --- | --- |
| EchAEP1 |  |  | 1.19 |  |
| G243E | 0 |  | >60 | 243 |
| G243F | 0 |  | >60 | 243 |
| G243N | 0 |  | >60 | 243 |
| G243P | 0 | LAD1+ | >60 | 243 |
| G243Q | 0 |  | >60 | 243 |
| G243R | 0 |  | >60 | 243 |
| G243Y | 0 |  | >60 | 243 |
| G243T | 0 |  | 29.49 | 243 |
| G243S | 0 |  | 6.81 | 243 |
| T244A | 0 |  | 5.26 | 244 |
| S215A | 0 |  | 3.57 | 215 |
| T244M | 0 |  | 3.09 | 244 |
| G173A | 0 | LAD2+ | 2.82 | 173 |
| S215G | 0 |  | 2.71 | 215 |
| P174W | 0 |  | 2.5 | 174 |
| T244V | 0 |  | 2.38 | 244 |
| P174F | 0 |  | 2.14 | 174 |
| P174A | 0 | LAD2+ | 1.94 | 174 |
| T244H | 0 |  | 1.85 | 244 |
| P174N | 0 |  | 1.79 | 174 |
| G63S | 0 |  | 1.76 | 63 |
| P174Y | 0 |  | 1.74 | 174 |
| G243A | 0 |  | 1.68 | 243 |
| T244E | 0 |  | 1.67 | 244 |
| T244I | 0 |  | 1.67 | 244 |
| T244S | 0 |  | 1.67 | 244 |
| T244W | 0 |  | 1.66 | 244 |
| P174M | 0 |  | 1.58 | 174 |
| S266G | 0 |  | 1.58 | 266 |
| P174C | 0 |  | 1.48 | 174 |
| P174Q | 0 |  | 1.45 | 174 |
| G63N | 0 |  | 1.42 | 63 |
| G63Q | 0 |  | 1.42 | 63 |
| P174R | 0 |  | 1.39 | 174 |
| G63F | 0 |  | 1.31 | 63 |
| T244R | 0 |  | 1.25 | 244 |

For some single mutants, no hydrolysis products were detected by HPLC at substrate conversions of 20–60%. To facilitate model training, their C_H_ratio values were set to 60, although the actual values were greater than 60.

**Table S2** X-ray data collection and refinement statistics.

| Parameter | EchAEP1–CtCYS | EchAEP1-G243Q |
| --- | --- | --- |
| PDB accession code | 9VA4 | 45ZO |
| Data collection |  |  |
| Beamline | SSRF BL10U2 | SSRF BL02U1 |
| Wavelength (Å) | 0.97918 | 0.979183 |
| Space group | P 3₂ 2 1 | P 2_1_ 2_1_ 2 |
| Unit-cell dimensions |  |  |
| a, b, c (Å) | 84.89, 84.89, 127.43 | 128.01, 78.3, 91.05 |
| α, β, γ (°) | 90.00, 90.00, 120.00 | 90.0, 90.0, 90.0 |
| Resolution range (Å)^a^ | 25.48–2.28 (2.34–2.28) | 53.86–2.02 (2.07–2.02) |
| Unique reflections | 24,842 (1,816) | 60767 (4454) |
| Multiplicity | 18.9 (16.8) | 13.3 (13.4) |
| Completeness (%) | 99.9 (100.0) | 100.0 (100.0) |
| Mean *I*/σ(*I*) | 11.9 (2.0) | 13.8 (2.2) |
| *R*_merge_ | 0.189 (1.949) | 0.183 (1.641) |
| CC_1/2_ | 0.996 (0.639) | 0.998 (0.660) |
| Refinement |  |  |
| Resolution range (Å) | 24.51–2.30 | 52.36–2.02 |
| *R*_work_/*R*_free_ | 0.1942/0.2373 | 0.1812 / 0.2202 |
| No. non-hydrogen atoms | 2,895 | 6761 |
| Protein | 2,824 | 6,334 |
| Water | 71 | 427 |
| No. protein residues | 361 | 814 |
| Mean B factor (Å²) | 53.22 | 33.55 |
| Protein | 53.21 | 33.38 |
| Water | 53.34 | 36.06 |
| R.m.s. deviations |  |  |
| Bond lengths (Å) | 0.006 | 0.0074 |
| Bond angles (°) | 0.806 | 0.8788 |
| Ramachandran favored (%) | 96.86 | 97.24 |
| Ramachandran outliers (%) | 0.0 | 0.25 |
| Rotamer outliers (%) | 1.0 | 0.75 |
| Clashscore | 3 | 4 |

^a^ Values in parentheses refer to the highest-resolution shell (2.34–2.28 Å). Outer-shell statistics not available from the deposited dataset are not shown.

Rmerge is expressed as a percentage; CC1/2 and Rwork/Rfree are dimensionless. R.m.s., root-mean-square; B factors are reported in Å².

**Supplementary Note 1: Results**

**1.1 Identification of functional sequence features and engineering of legumains using machine-learning sliding windows**

Protein language model embeddings can encode numerous sequence features that have not yet been explicitly recognized by human experts. We assembled 49 previously experimentally classified natural enzyme sequences, extracted ESMC-600M embeddings, and examined their distribution in a low-dimensional sequence space using principal component analysis (PCA). Overall, AEP, HYB, and PAL enzymes displayed a clear spatial progression corresponding to changes in ligation versus hydrolysis activity. EchAEP1 was located at the negative extreme of PC1, toward the most AEP-like end, making it a representative model (Figure S1). These findings suggest that the sequence-space distribution of functional classes within the legumain family may reflect evolutionary changes in ligation and hydrolysis activities.

However, earlier sequence discovery efforts were biased toward positive-class enzymes, leading to substantial class imbalance in the natural enzyme dataset, which contained 30 PALs, 12 HYBs, and only 7 AEPs (Figure S2a). To supplement the negative class, we introduced 1,334 putative AEPs predicted using the LAD hypothesis (LAD_AEP) from a database of 1,500 legumain sequences (Figure S2b). To assess this supplementation strategy, the 49 experimentally characterized enzyme sequences and 1,334 LAD_AEP sequences were merged, embedded, and jointly analyzed by PCA. The LAD_AEP centroid was found to lie close to that of experimentally validated AEPs (Figure S2c), suggesting that the LAD hypothesis offers useful AEP classification capability and supporting the use of LAD_AEP as an auxiliary source of negative-class data.

To further associate functional differences with specific sequence segments, we introduced a sliding-window analysis. First, enzyme sequences in the dataset were subjected to multiple-sequence alignment (MSA), and windows were extracted using Butelase1, the earliest discovered PAL, as reference (Figure S2a). Because window size and bag-of-words n-gram length can both affect feature extraction, a grid search was used to identify suitable combinations. For a sequence segment of window size w, bag-of-words features of lengths 1 through w−1 could be constructed. We then recorded and averaged the cross-validation performance of four machine-learning algorithms at each position for comparison (Figure S3).

Overall, model performance improved with increasing window size, although the incremental gains gradually plateaued. N-gram lengths of 2 or 3 generally performed well. Decision tree (DT) and logistic regression (LR) tended to favor length 3 in most cases, whereas random forest (RF) and support vector machine (SVM) performed best with length 2. DT was generally weaker and RF was the strongest overall. Notably, RF with a window length of 9 and n-gram length of 2 achieved the highest position-averaged ROC-AUC and F1 scores among all tested configurations. We therefore prioritized n-gram length 2 for subsequent analyses and examined how different window scales affected the identification of important sites.

To assess whether the models recognized known functional determinants, we calculated rank fractions for individual positions, focusing on LAD1 (reference position 237) and LAD2 (reference positions 167/168). With the small w2 window, position 168 ranked relatively poorly, whereas positions 167 and 237 ranked highly, suggesting that position 168 may have a weaker independent regulatory effect. Under larger windows, however, position 237 declined markedly in rank, suggesting that its functional signal may arise primarily from the residue itself and that neighboring sequence information may introduce interference (Figure S2d). When known LAD sites were used as references, ROC-AUC produced more balanced rankings overall, particularly for position 168 under small windows and position 237 under large windows. Accordingly, ROC-AUC was selected as the evaluation metric for subsequent modeling.

Inspection of the position-importance heatmaps showed that important regions identified by the four machine-learning algorithms were discontinuous in sequence but highly consistent across algorithms. Small and large windows also showed different feature-extraction tendencies. Small windows more readily identified key positions with strong independent effects but were less capable of recognizing potential cooperative sites. Large windows could capture potential relationships among multiple positions but might dilute the importance signals of individual sites (Figure S2e). We therefore combined the small w2 window (n-gram length 1) and large w9 window (n-gram length 2) in subsequent analyses to capture both independently acting positions and potentially cooperative regions.

Optimal window–algorithm combinations were selected from cross-validation results and used to train the corresponding submodels. With Butelase1 as the reference sequence, position-level model scores were calculated and important regions were mapped onto the protein three-dimensional structure. Although the regions identified by the models were discontinuous in sequence, they clustered spatially around the substrate-binding and catalytic interface. Specifically, w2 showed high sensitivity to LAD1 and LAD2, whereas w9 highlighted the MLA region. In addition, w2 identified an autocatalytic activation cleavage site in the distal linker region, suggesting possible coevolutionary relationships between that region and reaction directionality (Figure 2a,b).

The same minimum-score thresholds were then applied to select submodels and construct ensemble models, which were trained and retrospectively evaluated on all 49 non-LAD-labeled sequences. Both w2 and w9 showed high classification performance, with accuracies of 95.92% and 97.96%, balanced accuracies of 97.62% and 98.81%, and Matthews correlation coefficients (MCCs) of 0.861 and 0.924, respectively. Both models correctly identified all seven AEPs, while w9 misclassified fewer CYC sequences and performed slightly better overall than w2 in this retrospective assessment (Figure S5). These findings indicate that multiscale sliding-window models can capture sequence features associated with legumain functional classifications and provide useful classification performance on the natural enzyme collection.

We next applied the model to the directed engineering of EchAEP1. In silico single-site saturation mutagenesis of the EchAEP1 core domain, followed by initial model-based screening, yielded 511 candidate substitutions. Combined selection based on predicted stability changes and model-score gains retained 133 candidates. Considering the distribution density of candidate substitutions across sites, 92 single mutants at positions 86, 196, 197, 238, 243, 244, 288, and 289 were selected for fluorescence screening. Following this screening, 39 variants were excluded and 53 candidates proceeded to chromatographic evaluation.

Four of these single mutants did not yield valid HPLC measurements because of conversion-related limitations. Together with an additional wild-type EchAEP1 HPLC measurement, 50 valid datasets were obtained, comprising 49 single mutants and one wild-type control. Among the 49 single mutants, 36 exhibited greater ligation activity than wild-type EchAEP1, including 33 positive substitutions outside the original prediction range of the LAD hypothesis (Table S1). Specifically, 17 positive substitutions occurred at LAD sites but were not predicted by the original LAD rules, while another 16 positive substitutions occurred at other sites. These results indicate that the machine-learning models not only identify established LAD regions but also extend the potential mutational space for legumain functional engineering beyond the predictions of the original empirical hypothesis.

**1.2 Activity landscape of the local sequence space**

Within the sequence space represented by protein language model embeddings, substitutions at position 243 clustered in a region corresponding to a prominent peak in cyclization activity. Although other positive single mutants shifted toward this region relative to wild-type EchAEP1, they did not cross the cyclization activity cliff (Figure S6). We therefore generated all possible 2–5-site combinations of positive single mutants to examine the influence of combined mutations on reaction directionality. To make full use of round-0 chromatographic cyclization data, we also constructed a ch_ratio prediction model and predicted the activity of all combinatorial mutants. Compared with single mutants, combinatorial variants showed further displacement and a broader distribution in sequence space (Figure S7).

To explore the transpeptidation activity peak and its boundary, we applied a joint candidate-selection strategy based on predicted ch_ratio and sequence-space distance, selecting 30 combinatorial mutants beyond the opposite side of the peak for experimental validation (Figures S9 and S10). Because cyclization near the peak yielded almost no hydrolysis product and could not readily differentiate activities among variants, subsequent measurements were switched to intermolecular ligation assays, which allowed greater competition from hydrolysis. Experimentally, ligation activity did not continue to increase beyond the peak; instead, many mutants remained expressible but had no detectable activity (Figure S11). Thus, moving farther beyond the transpeptidation activity peak in sequence space did not necessarily improve transpeptidation and could instead result in loss of catalytic function.

To increase sampling density near the transpeptidation activity peak and explore insufficiently sampled regions, a second screening round was performed. Activities also varied substantially within the peak region, with several combinations showing favorable ligation activity clustered around the 243Q and 243N local regions. The 243Q-244I and 243N-244I variants were especially prominent, with crude-enzyme lh_ratio values of 1.37 (46.0% conversion) and 1.24 (46.8% conversion), respectively, approaching the positive-control VyPAL2-I244V value of 1.51 (45.1% conversion) (Figure S15). These results indicate that even within nearby high-activity regions of sequence space, differences in local sequence context can substantially affect catalytic behavior.

To map the local activity landscape of EchAEP1 systematically, experimental ch_ratio and lh_ratio data for EchAEP1 and all mutants were integrated to construct a local sequence-space activity map (Figures S16–S18). The cyclization activity cliff was concentrated around changes at a small number of key positions, whereas a broad inactive region lay beyond the other side of the transpeptidation activity peak (Figure S19). These observations reveal pronounced heterogeneity in the local EchAEP1 transpeptidation landscape: substitutions at a few critical positions can produce large activity shifts, but further departure from the peak does not necessarily increase activity and may abolish catalysis. Thus, the evolution of EchAEP1 activity is jointly constrained by key-site effects and the broader sequence context, and its high-activity region has a distinct boundary in sequence space.

The LAD hypothesis has been widely used for functional classification and directed engineering of the legumain family and has shown practical value in multiple natural enzyme backgrounds. Once sequence-space position had been mapped to EchAEP1 catalytic activity, however, an additional question emerged: along what sequence-space trajectories would the combinations of determinants specified by the LAD hypothesis drive EchAEP1? We therefore generated all candidate EchAEP1 sequences guided by LAD rules and extracted their protein language model embeddings to examine the distribution of different LAD combinations. LAD candidates followed a relatively distinct trajectory, beginning in the hydrolytic region, crossing the cyclization activity cliff, and reaching the transpeptidation activity peak without extending into the inactive region beyond it (Figure S20).

Notably, different LAD-site combinations occupied markedly different regions of sequence space. LAD2+ candidates remained largely clustered around EchAEP1, whereas candidates containing LAD1+ shifted substantially toward the transpeptidation activity peak; both LAD1+ and LAD1+&LAD2+ combinations approached or reached the peak. These observations suggest that LAD1 and LAD2 may contribute differently to EchAEP1 movement through sequence space, and that LAD1-related substitutions may be the main driver enabling EchAEP1 to cross the cyclization activity cliff and acquire greater transpeptidation activity. Thus, protein language model embedding space provides not only a representation of function-associated natural legumain sequence distributions but also a perspective for examining site-specific effects and combinatorial rules under the established LAD hypothesis.

**1.3 Relationship between substrate-binding conformations and oxyanion-hole formation**

To examine the relationship between substrate-binding conformations and oxyanion-hole formation, we jointly analyzed substrate-related distances and oxyanion-hole hydrogen-bond states in three independent simulations of EchAEP1, EchAEP1-G243Q, and EchPAL1 at pH 5.5 and 7.0, using a common 0–500 ns observation window (Figures SN1–SN6). The hydrogen-bonding contact between A:E211 and the side chain of substrate residue B:N5 is located at the base of the active-site pocket. Accordingly, the A:E211 OE1–B:N5 ND2 distance was used to characterize the position of the N5 side chain relative to the pocket base. The A:G173 Cα–B:S6 Cα distance was monitored as an additional descriptor of the position of the P1′-side peptide segment relative to the catalytic center; the corresponding enzyme residue in EchPAL1 is A:A173. Oxyanion-hole states were assigned according to the formation of the two hydrogen bonds donated by the backbone N–H groups of A:C213 and A:G172 to the backbone carbonyl O of B:N5.

The trajectories showed that displacement of the N5 side chain from its binding position at the pocket base did not necessarily coincide with loss of the oxyanion-hole hydrogen-bond network. For example, in rep2 of EchAEP1 at pH 5.5, the A:E211 OE1–B:N5 ND2 distance increased from approximately 3–4 Å to approximately 8–10 Å after about 210 ns, while the two oxyanion-hole hydrogen bonds continued to form simultaneously and frequently (Figure SN1). Comparing the first and second 250 ns of this trajectory, the mean distance increased from 4.46 to 9.00 Å, whereas the fraction of frames with both hydrogen bonds present increased from 39.4% to 66.7%. A similar pattern occurred in rep0 of EchAEP1-G243Q at pH 7.0: the mean distances in the first and second 250 ns were 4.31 and 7.19 Å, respectively, with corresponding dual-hydrogen-bond occupancies of 33.8% and 61.9% (Figure SN4). These within-trajectory comparisons show that intervals dominated by larger side-chain–pocket distances can exhibit higher oxyanion-hole occupancies than intervals with smaller distances.

Additional examples were observed during several early intervals of rep2 of EchAEP1-G243Q at pH 5.5. Despite the relatively large separation between the N5 side chain and E211, both oxyanion-hole hydrogen bonds formed frequently, particularly during approximately 50–125 ns and 150–200 ns (Figure SN2). In rep2 of EchPAL1 at pH 5.5, an increase in this distance after approximately 425 ns was also accompanied by an increase in dual-hydrogen-bond occupancy, particularly during the final 50 ns (Figure SN5). Together, these observations indicate that the substrate backbone carbonyl can retain the complete two-hydrogen-bond coordination of the oxyanion hole even when the N5 side chain no longer maintains its original binding position at the pocket base.

Conversely, renewed proximity of the N5 side chain to E211 did not guarantee restoration of the oxyanion-hole hydrogen-bond network. During the later part of rep0 of EchAEP1-G243Q at pH 5.5, the A:E211 OE1–B:N5 ND2 distance decreased to approximately 4–5 Å, yet neither oxyanion-hole hydrogen bond was present in any sampled frame between 350 and 500 ns (Figure SN2). Thus, a shorter side-chain–pocket distance was not invariably associated with oxyanion-hole formation.

Taken together, the sampled trajectories indicate that positioning of the N5 side chain at the pocket base and coordination of its backbone carbonyl within the oxyanion hole are distinct conformational features that need not change synchronously. Displacement of the side chain does not necessarily result in complete suppression of the monitored oxyanion-hole network, and proximity to the pocket base does not necessarily ensure its formation. Oxyanion-hole states therefore cannot be inferred from N5 side-chain positioning alone and should instead be assessed directly from the relevant backbone hydrogen bonds. The higher dual-hydrogen-bond occupancies observed in some larger-distance intervals do not establish that side-chain displacement generally promotes oxyanion-hole formation; rather, they demonstrate that the relationship cannot be described by a simple one-to-one correspondence.

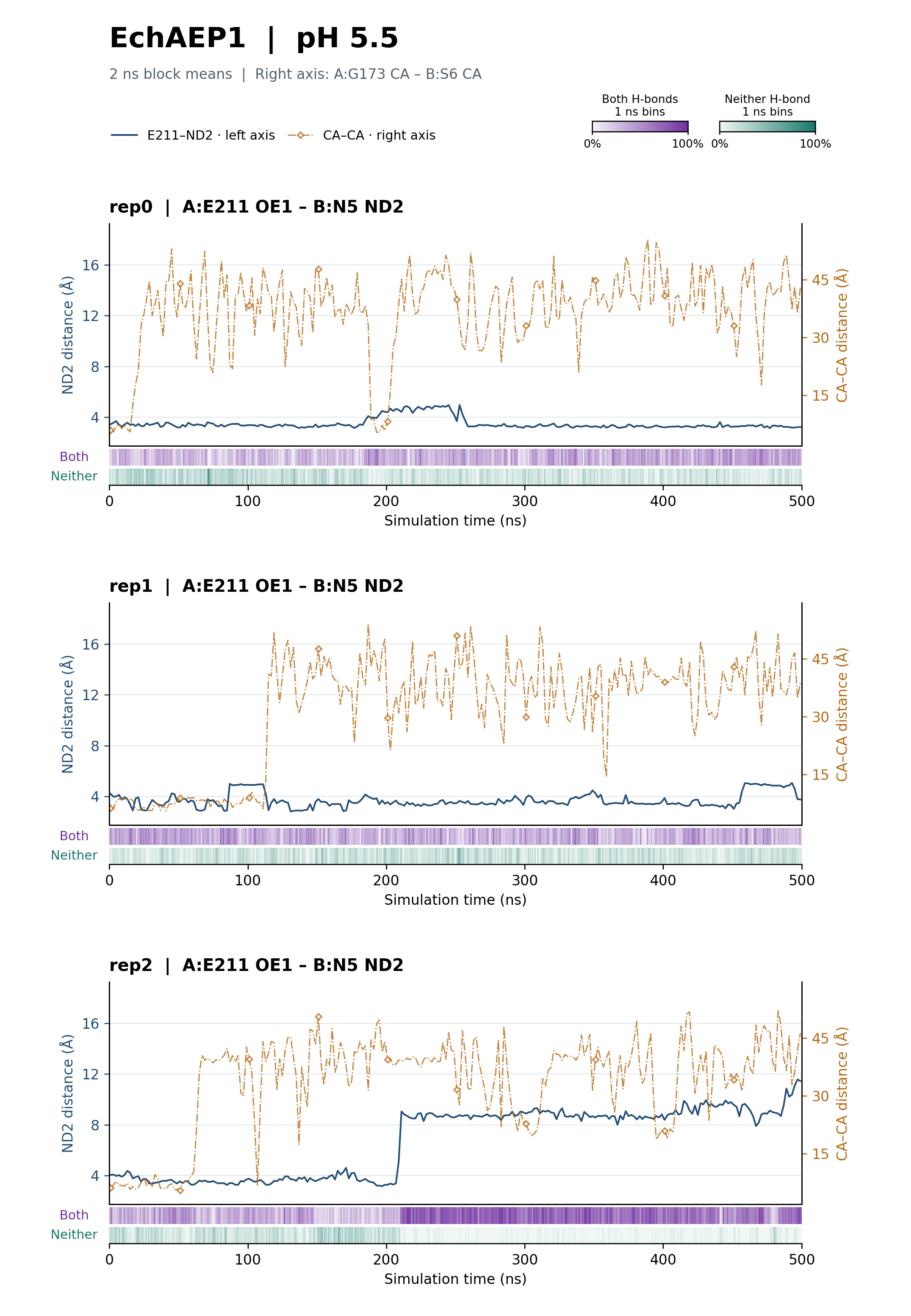

Figure SN1. Substrate-binding descriptors and oxyanion-hole states of EchAEP1 at pH 5.5. The top, middle, and bottom panels show the 0–500 ns trajectories of three independent simulations, rep0, rep1, and rep2, respectively. The blue solid line shows the A:E211 OE1–B:N5 ND2 distance on the left axis, and the orange dash-dotted line shows the A:G173 Cα–B:S6 Cα distance on the right axis. Distances are expressed in Å and displayed as means over non-overlapping 2 ns bins. The purple “Both” and teal “Neither” strips below each panel show the fractions of frames in which both or neither of the hydrogen bonds from the backbone N–H groups of A:C213 and A:G172 to the backbone carbonyl O of B:N5 are present. Fractions are normalized to all sampled frames in each 1 ns bin and mapped to a 0–100% color scale. Frames containing only one hydrogen bond belong to neither state, so the two fractions need not sum to 100%.

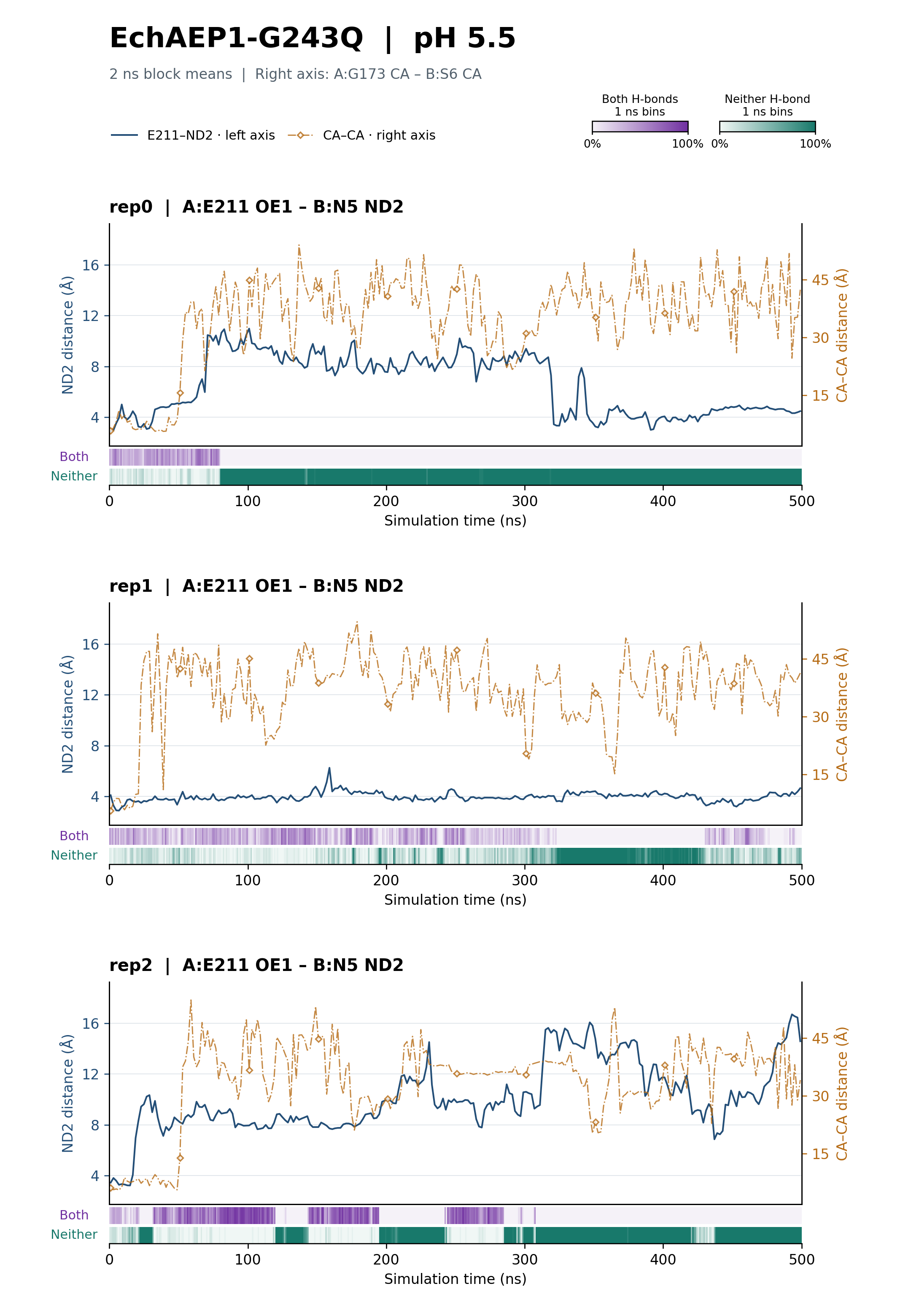

Figure SN2. Substrate-binding descriptors and oxyanion-hole states of EchAEP1-G243Q at pH 5.5. The top, middle, and bottom panels show the 0–500 ns trajectories of three independent simulations, rep0, rep1, and rep2, respectively. The blue solid line shows the A:E211 OE1–B:N5 ND2 distance on the left axis, and the orange dash-dotted line shows the A:G173 Cα–B:S6 Cα distance on the right axis. Distances are expressed in Å and displayed as means over non-overlapping 2 ns bins. The purple “Both” and teal “Neither” strips below each panel show the fractions of frames in which both or neither of the hydrogen bonds from the backbone N–H groups of A:C213 and A:G172 to the backbone carbonyl O of B:N5 are present. Fractions are normalized to all sampled frames in each 1 ns bin and mapped to a 0–100% color scale. Frames containing only one hydrogen bond belong to neither state, so the two fractions need not sum to 100%.

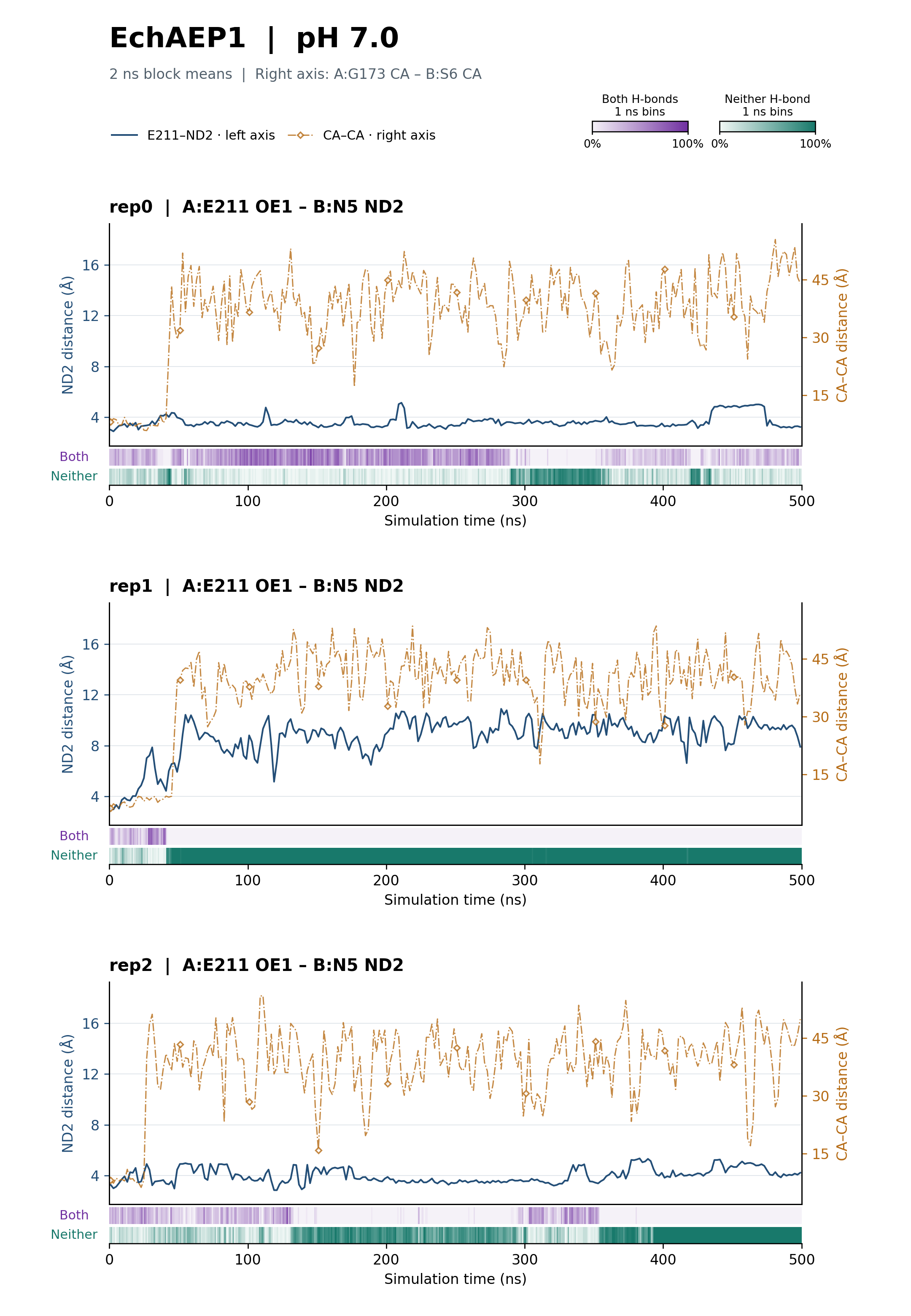

Figure SN3. Substrate-binding descriptors and oxyanion-hole states of EchAEP1 at pH 7.0. The top, middle, and bottom panels show the 0–500 ns trajectories of three independent simulations, rep0, rep1, and rep2, respectively. The blue solid line shows the A:E211 OE1–B:N5 ND2 distance on the left axis, and the orange dash-dotted line shows the A:G173 Cα–B:S6 Cα distance on the right axis. Distances are expressed in Å and displayed as means over non-overlapping 2 ns bins. The purple “Both” and teal “Neither” strips below each panel show the fractions of frames in which both or neither of the hydrogen bonds from the backbone N–H groups of A:C213 and A:G172 to the backbone carbonyl O of B:N5 are present. Fractions are normalized to all sampled frames in each 1 ns bin and mapped to a 0–100% color scale. Frames containing only one hydrogen bond belong to neither state, so the two fractions need not sum to 100%.

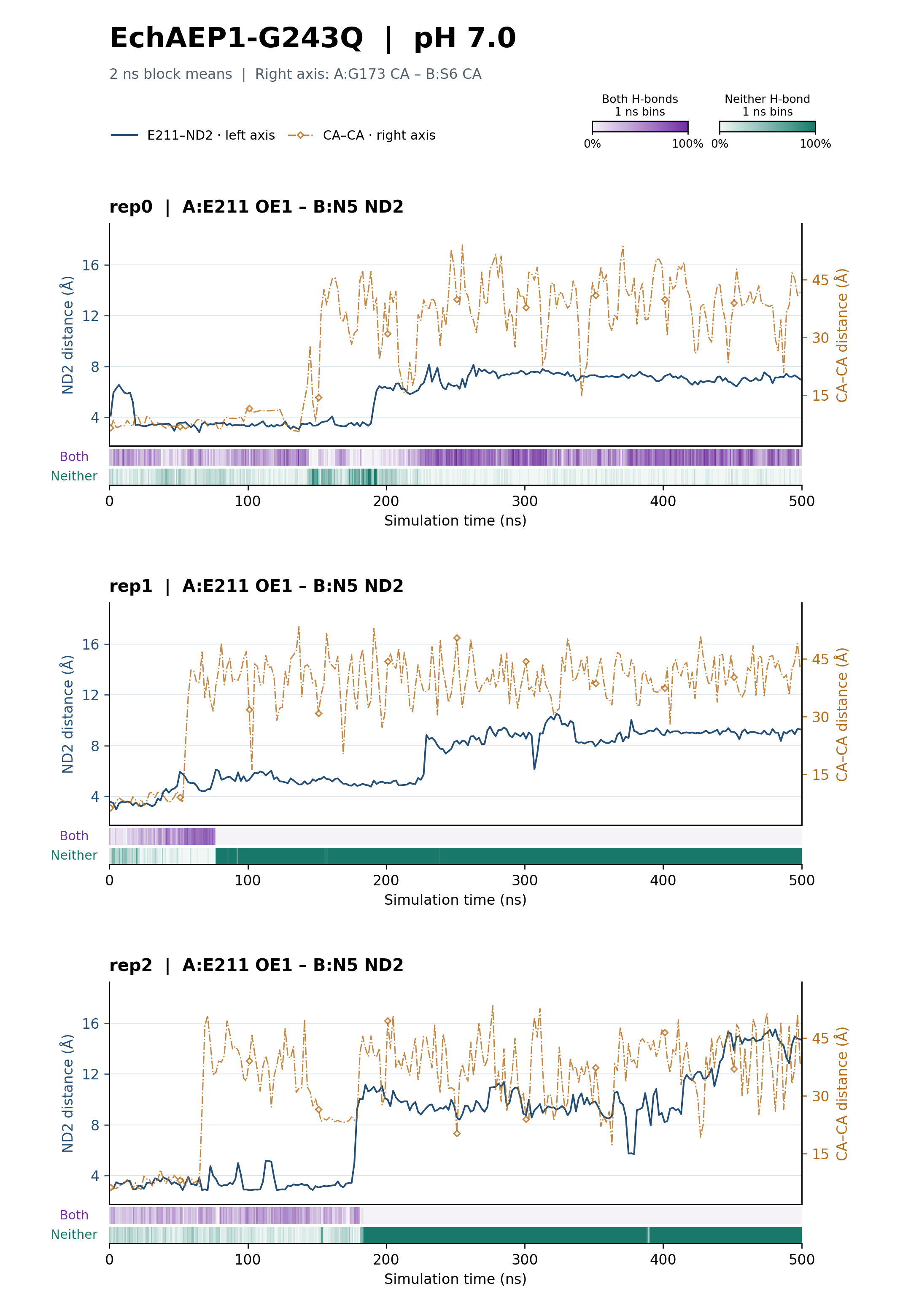

Figure SN4. Substrate-binding descriptors and oxyanion-hole states of EchAEP1-G243Q at pH 7.0. The top, middle, and bottom panels show the 0–500 ns trajectories of three independent simulations, rep0, rep1, and rep2, respectively. The blue solid line shows the A:E211 OE1–B:N5 ND2 distance on the left axis, and the orange dash-dotted line shows the A:G173 Cα–B:S6 Cα distance on the right axis. Distances are expressed in Å and displayed as means over non-overlapping 2 ns bins. The purple “Both” and teal “Neither” strips below each panel show the fractions of frames in which both or neither of the hydrogen bonds from the backbone N–H groups of A:C213 and A:G172 to the backbone carbonyl O of B:N5 are present. Fractions are normalized to all sampled frames in each 1 ns bin and mapped to a 0–100% color scale. Frames containing only one hydrogen bond belong to neither state, so the two fractions need not sum to 100%.

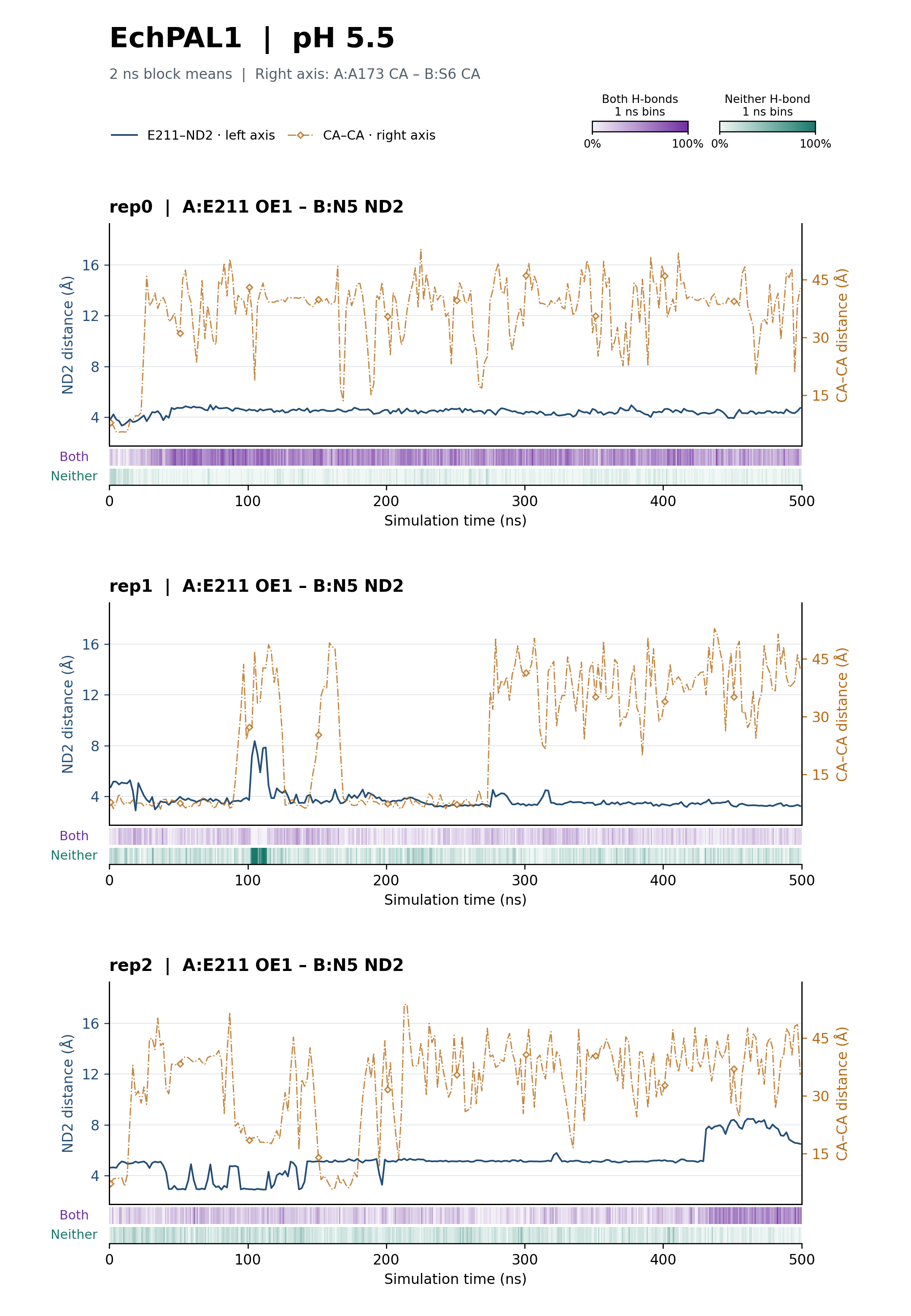

Figure SN5. Substrate-binding descriptors and oxyanion-hole states of EchPAL1 at pH 5.5. The top, middle, and bottom panels show the 0–500 ns trajectories of three independent simulations, rep0, rep1, and rep2, respectively. The blue solid line shows the A:E211 OE1–B:N5 ND2 distance on the left axis, and the orange dash-dotted line shows the A:A173 Cα–B:S6 Cα distance on the right axis. Distances are expressed in Å and displayed as means over non-overlapping 2 ns bins. The purple “Both” and teal “Neither” strips below each panel show the fractions of frames in which both or neither of the hydrogen bonds from the backbone N–H groups of A:C213 and A:G172 to the backbone carbonyl O of B:N5 are present. Fractions are normalized to all sampled frames in each 1 ns bin and mapped to a 0–100% color scale. Frames containing only one hydrogen bond belong to neither state, so the two fractions need not sum to 100%.

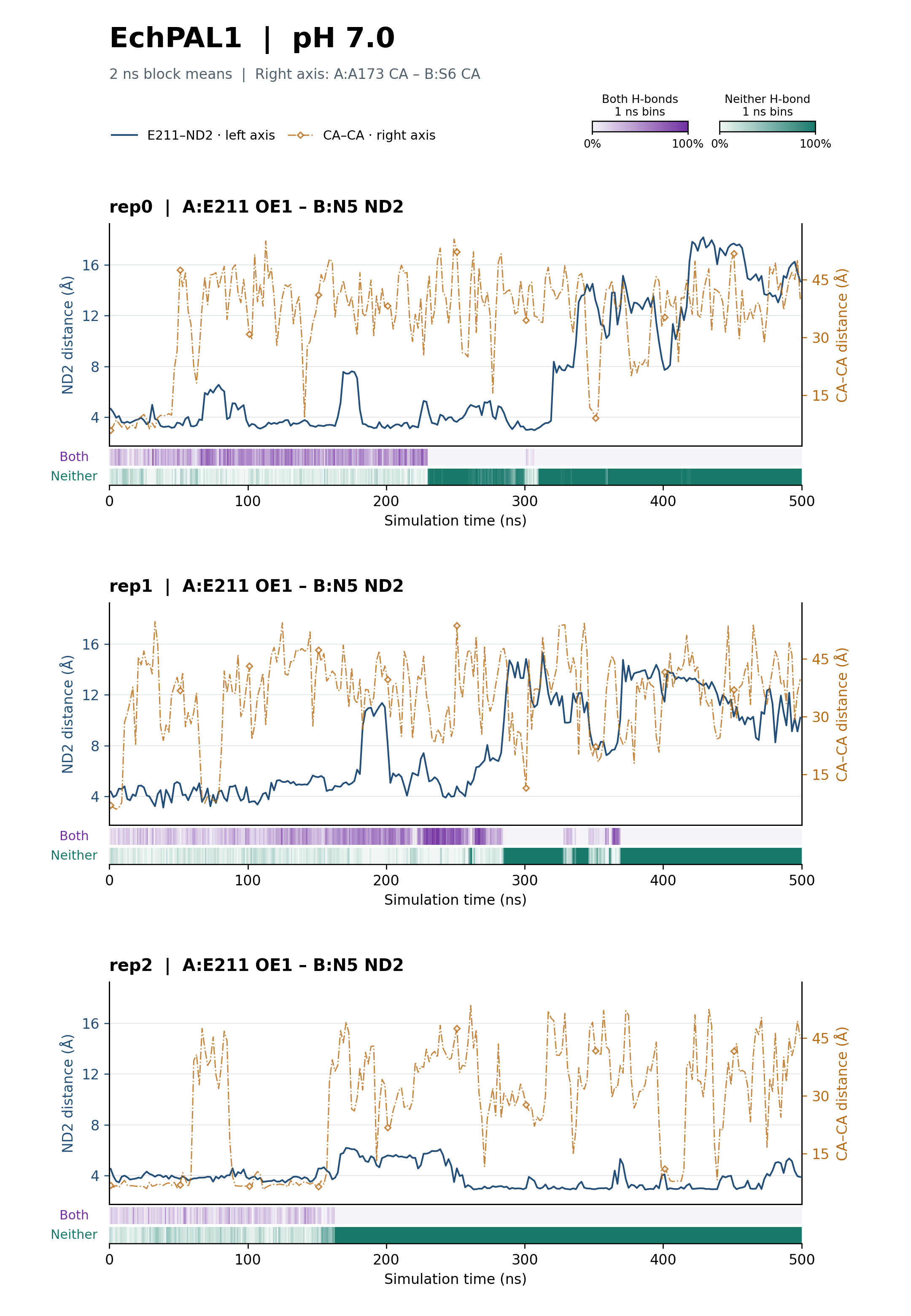

Figure SN6. Substrate-binding descriptors and oxyanion-hole states of EchPAL1 at pH 7.0. The top, middle, and bottom panels show the 0–500 ns trajectories of three independent simulations, rep0, rep1, and rep2, respectively. The blue solid line shows the A:E211 OE1–B:N5 ND2 distance on the left axis, and the orange dash-dotted line shows the A:A173 Cα–B:S6 Cα distance on the right axis. Distances are expressed in Å and displayed as means over non-overlapping 2 ns bins. The purple “Both” and teal “Neither” strips below each panel show the fractions of frames in which both or neither of the hydrogen bonds from the backbone N–H groups of A:C213 and A:G172 to the backbone carbonyl O of B:N5 are present. Fractions are normalized to all sampled frames in each 1 ns bin and mapped to a 0–100% color scale. Frames containing only one hydrogen bond belong to neither state, so the two fractions need not sum to 100%.

**1.4 Sensitivity analysis of conditional probabilities**

**1.4.1 pH5.5**

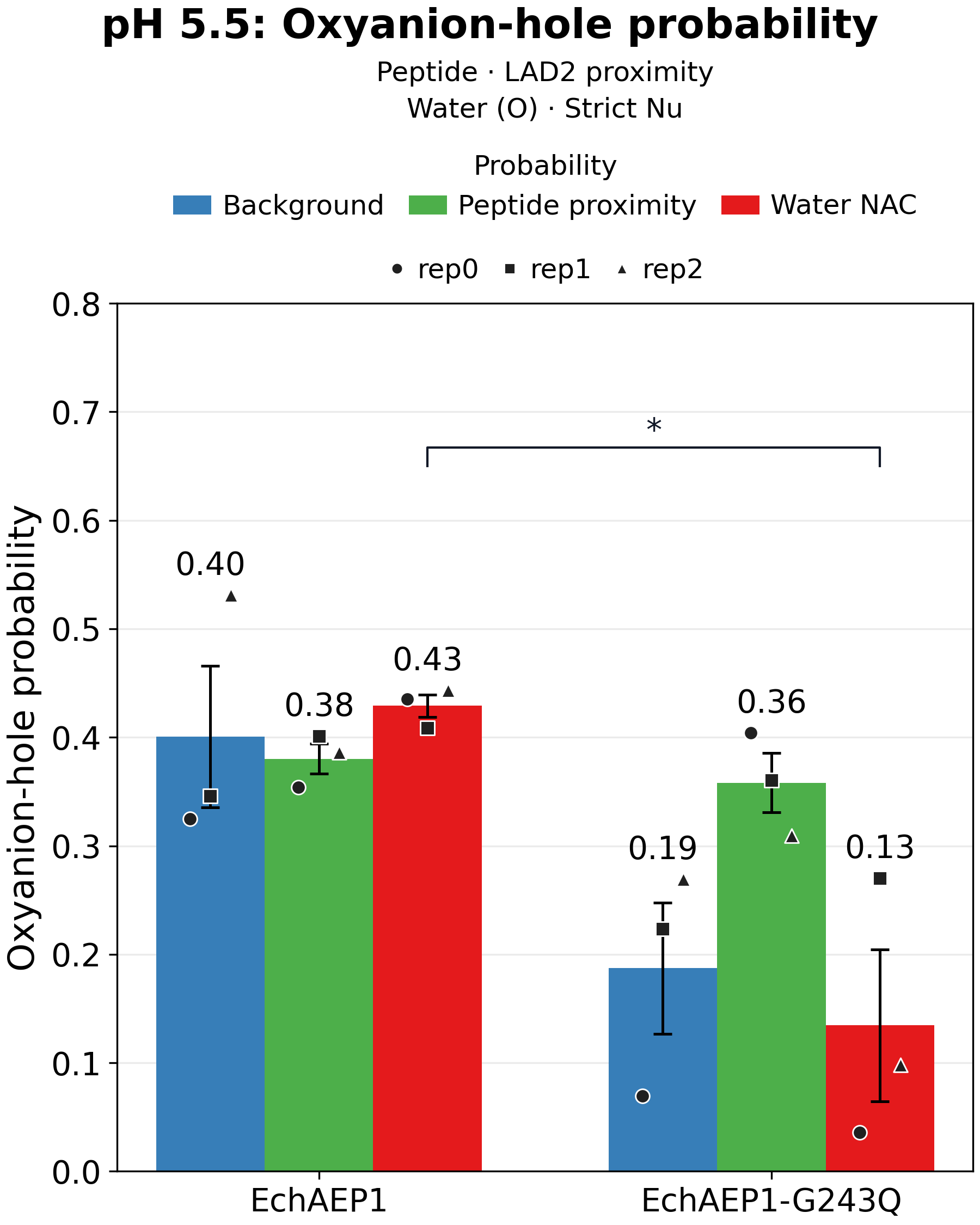

Figure SN7. Background and conditional oxyanion-hole probabilities for EchAEP1 and G243Q at pH 5.5. Blue bars indicate background oxyanion-hole probability, green bars the conditional probability in the presence of an incoming peptide, and red bars the conditional probability in the presence of an attacking water molecule. The Proximity criterion was applied to peptide and Strict Nu to attacking water.

Under the Proximity criterion, the number of peptide-condition frames increased from tens under Loose Nu to hundreds or more. Specifically, the three EchAEP1 replicates increased from 5/33/41 to 571/2220/1238 frames, while G243Q increased from 19/19/21 to 970/430/979 frames (Figure SN8). Under Strict Nu, qualifying attacking-water frames decreased from thousands under Loose Nu to hundreds: for EchAEP1, from 4072/1768/5484 to 825/296/571, and for G243Q, from 6098/5947/6651 to 675/711/959 (Figure SN8). Despite these changes in screening criteria, the observation from the main text—that the oxyanion hole is suppressed in G243Q at pH 5.5 but that this suppression is relieved in the presence of incoming peptide—was retained (Figure SN7).

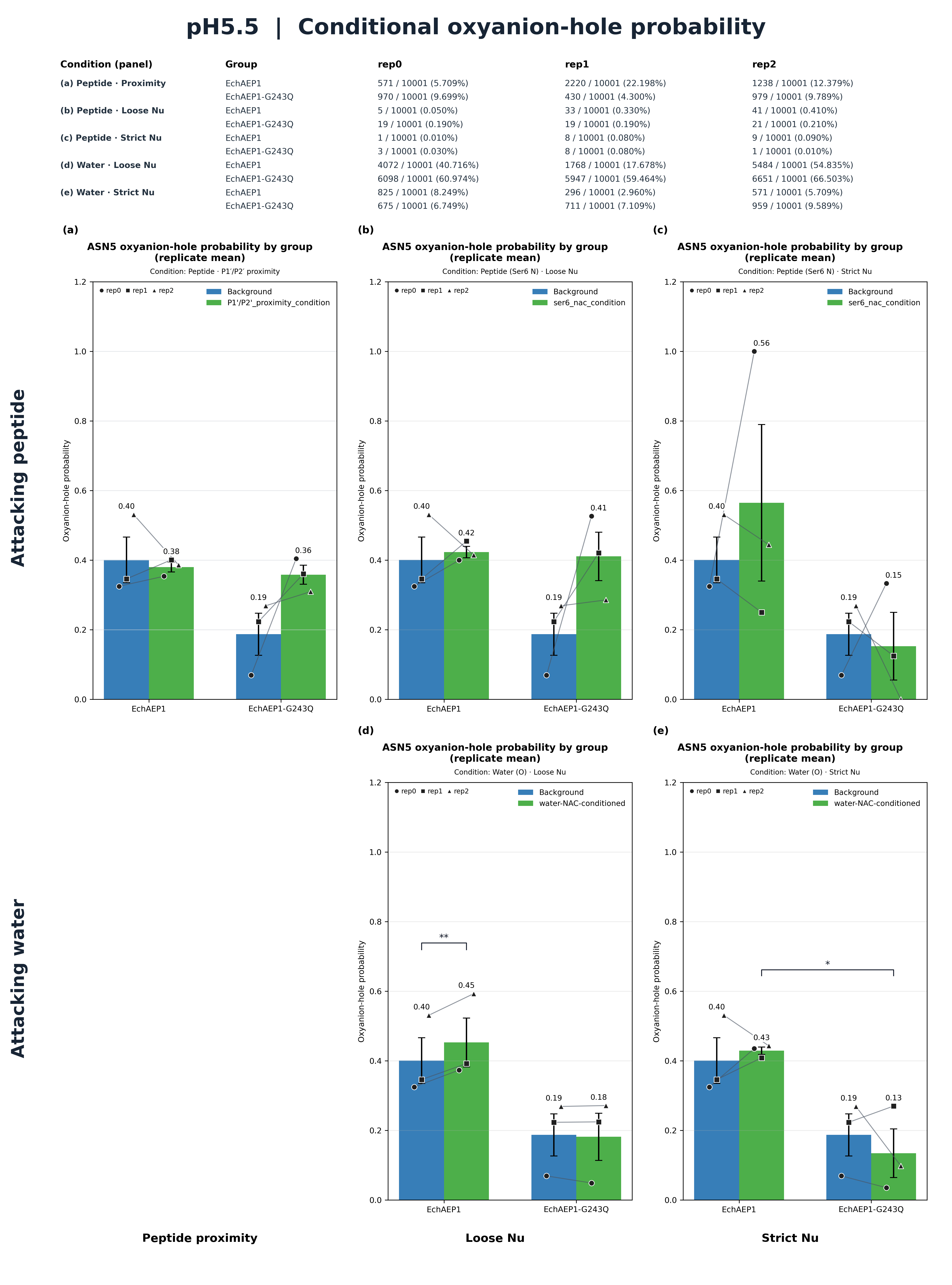

Figure SN8. Comparison of background and conditional oxyanion-hole probabilities in pH 5.5 simulations under different condition definitions. The subtitle gives the number and corresponding fraction of trajectory frames available for conditional-probability calculations in each replicate under each criterion.

Under Strict Nu, peptide-condition frames declined further from tens under Loose Nu to single digits. The three EchAEP1 replicates decreased from 5/33/41 to 1/8/9 frames, and G243Q from 19/19/21 to 3/8/1 frames (Figure SN8). Because some replicates contained only one qualifying frame, conditional probabilities were strongly influenced by individual conformations and showed substantial variability. Under these conditions, the peptide-conditioned probability in EchAEP1 increased 1.4-fold relative to background (0.56/0.40), whereas the G243Q conditional probability was slightly below its background, failing to reproduce the marked increase observed under the other criteria.

**1.4.2 pH7.0**

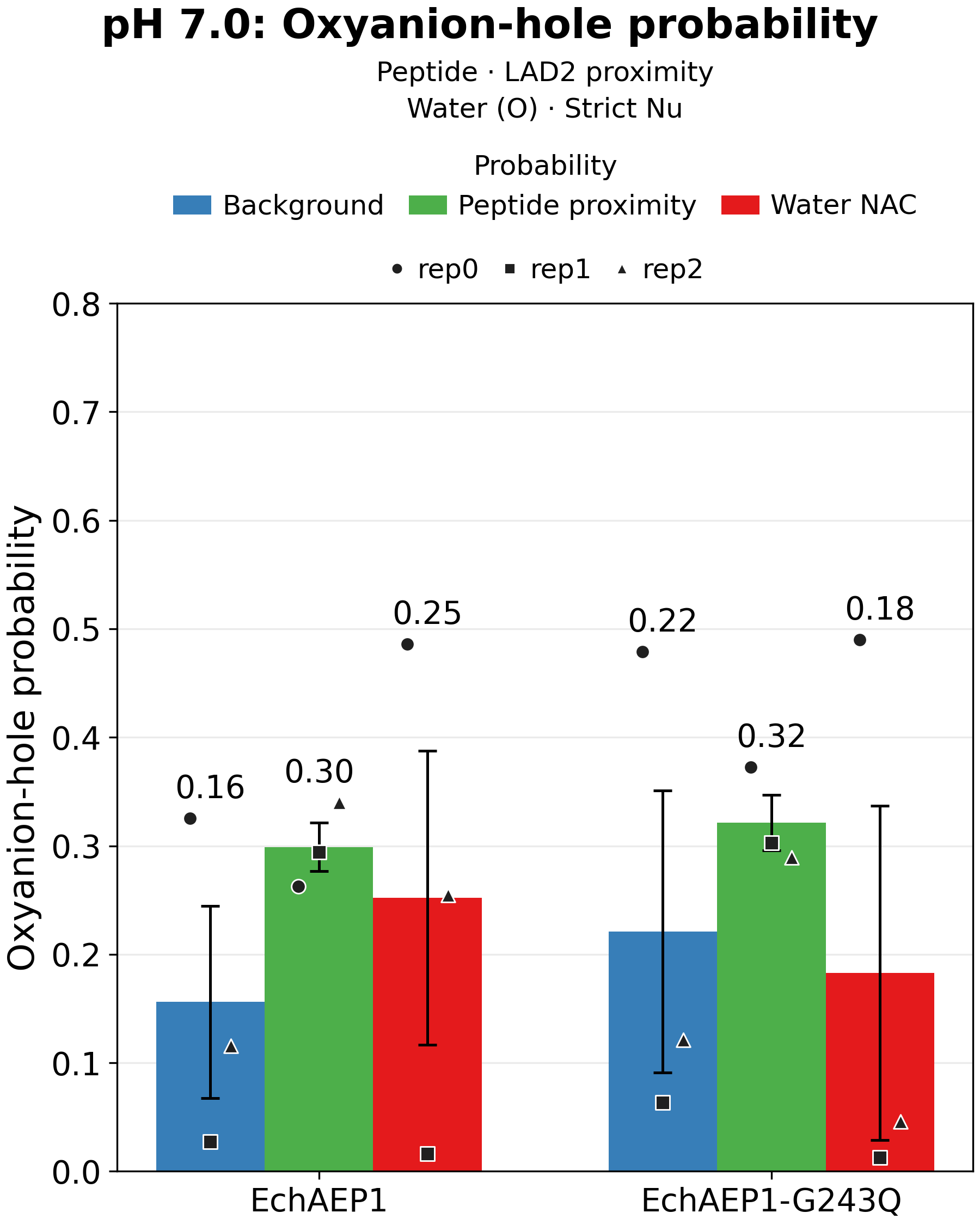

Figure SN9. Background and conditional oxyanion-hole probabilities for EchAEP1 and G243Q at pH 7.0. Blue bars indicate background oxyanion-hole probability, green bars the conditional probability in the presence of an incoming peptide, and red bars the conditional probability in the presence of an attacking water molecule. The Proximity criterion was applied to peptide and Strict Nu to attacking water.

Under Proximity, peptide-condition frames increased from tens under Loose Nu to hundreds or more. For the three EchAEP1 replicates, the counts increased from 52/13/14 to 815/890/497 frames, and for G243Q from 101/13/11 to 2651/1018/1289 (Figure SN10). Under Strict Nu, qualifying water frames decreased from thousands under Loose Nu to hundreds: EchAEP1 decreased from 5452/8319/1809 to 1387/803/114, and G243Q from 6085/5563/6071 to 712/546/810 (Figure SN10). Although the alternative screening criteria reduced peptide-conditioned probabilities in EchAEP1 and G243Q and increased the water-conditioned probability in EchAEP1, the pH 7.0 trend in oxyanion-hole behavior characteristic of ligases, as observed for EchAEP1 in the main text, remained broadly consistent (Figure SN9).

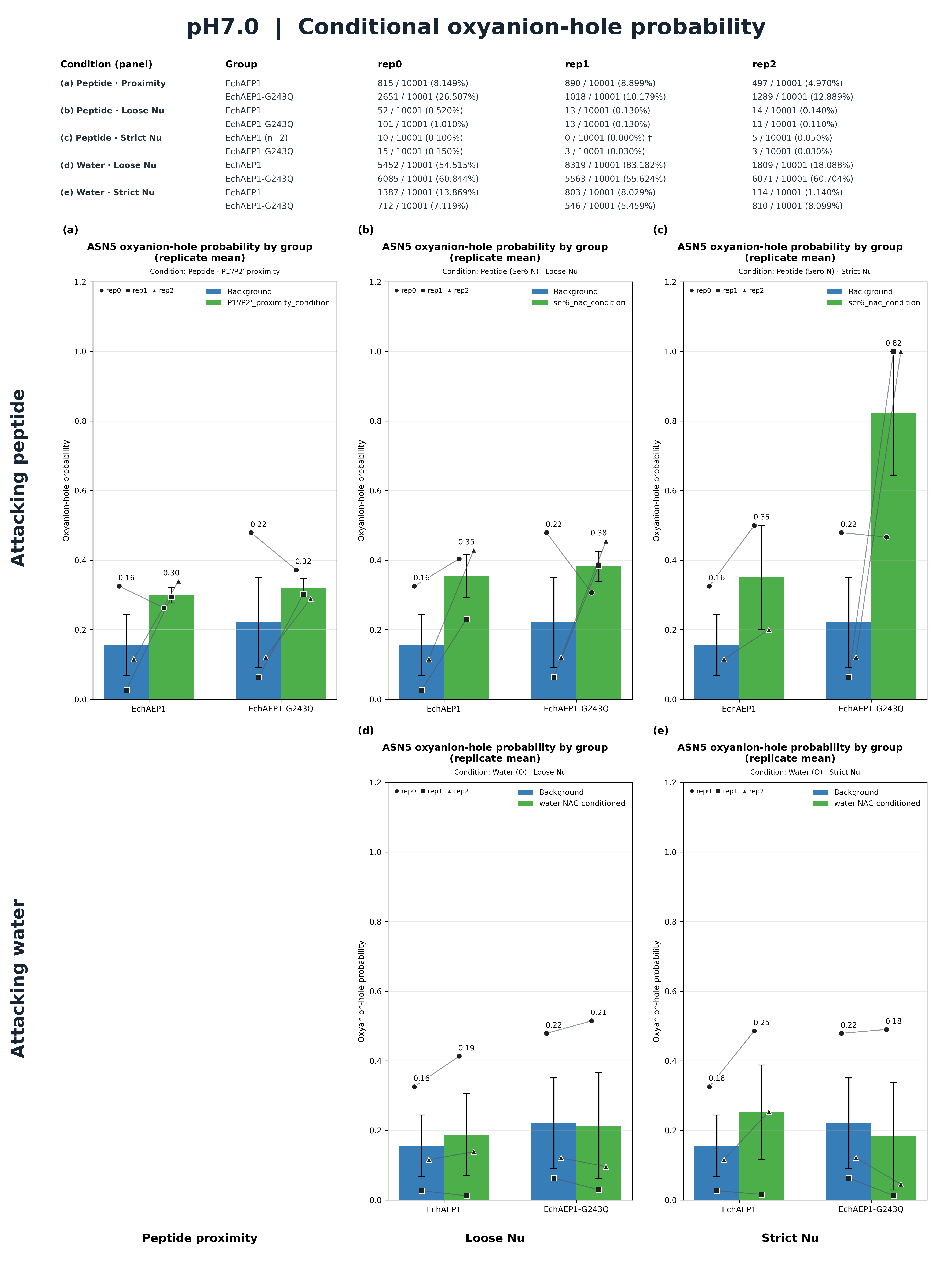

Figure SN10. Comparison of background and conditional oxyanion-hole probabilities in pH 7.0 simulations under different condition definitions. The subtitle gives the number and corresponding fraction of trajectory frames available for conditional-probability calculations in each replicate under each criterion.

Under Strict Nu, peptide-condition frames again decreased from tens under Loose Nu to single digits. EchAEP1 replicates decreased from 52/13/14 to 10/0/5 frames, while G243Q decreased from 101/13/11 to 15/3/3 (Figure SN10). The peptide-conditioned oxyanion-hole probability in G243Q was then 3.72 times its background (0.82/0.22), substantially greater than under Loose Nu or Proximity. However, this result was based on very few qualifying conformational frames and had poor statistical robustness; it was therefore insufficient to establish a stronger G243Q response to incoming peptide under this criterion.

**1.4.3 EchPAL1**

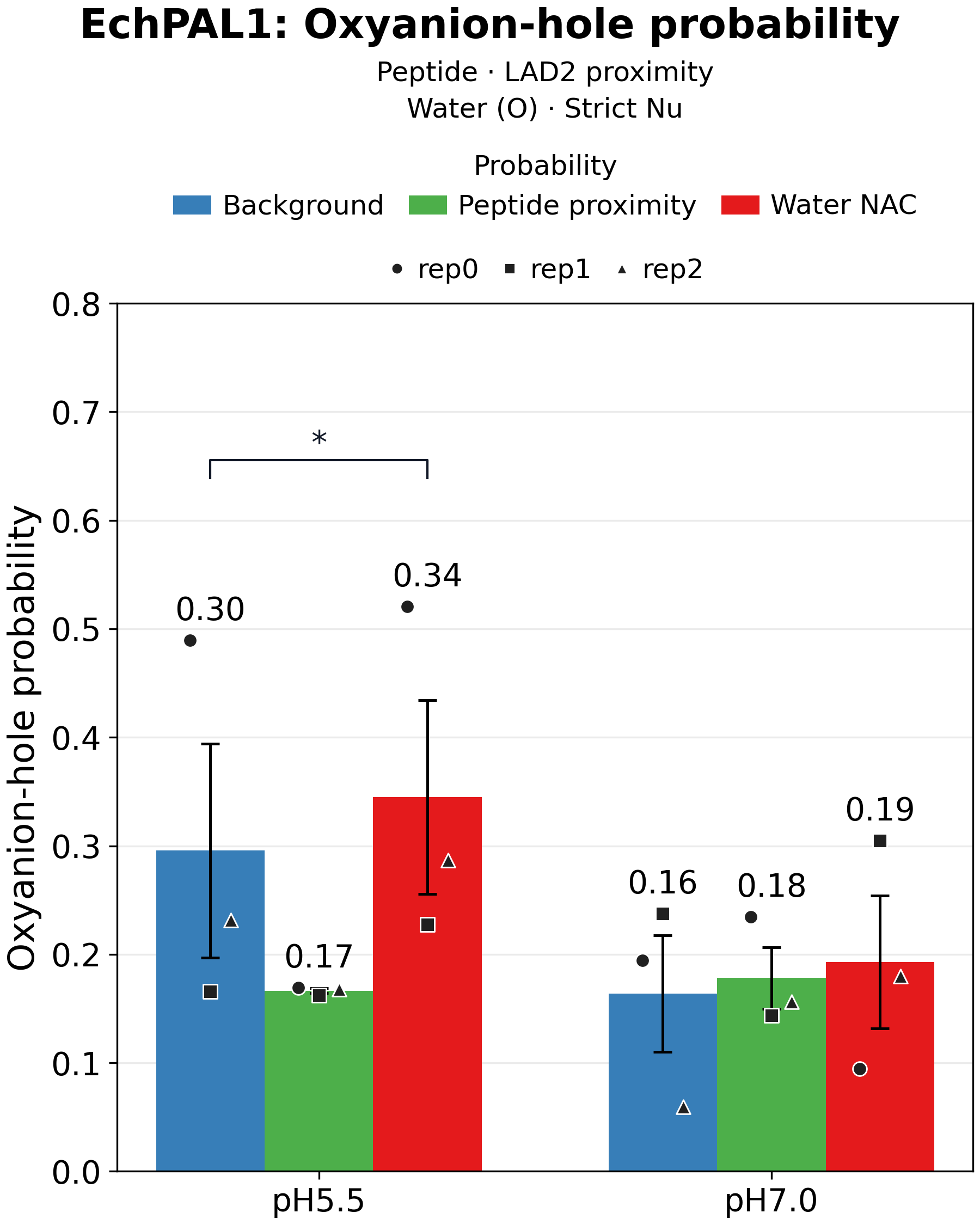

Figure SN11. Background and conditional oxyanion-hole probabilities for EchPAL1 at pH 5.5 and pH 7.0. Blue bars indicate background oxyanion-hole probability, green bars the conditional probability in the presence of incoming peptide, and red bars the conditional probability in the presence of attacking water. Proximity was used for peptide and Strict Nu for water.

Under Proximity, peptide-condition frames increased from tens under Loose Nu to hundreds or more. For the three replicates at pH 5.5, the counts increased from 1/36/38 to 484/4259/835, and for pH 7.0 from 4/35/52 to 1023/528/2972 (Figure SN12). Under Strict Nu, qualifying water frames decreased from thousands under Loose Nu to hundreds: from 8197/5057/4878 to 2142/897/820 at pH 5.5 and from 4048/5933/4677 to 339/758/523 at pH 7.0 (Figure SN12). Despite these changes in screening criteria, neither alternative criterion altered the main conclusion that incoming peptide in EchPAL1 was associated with a reduction, rather than an increase, in oxyanion-hole formation probability (Figure SN11).

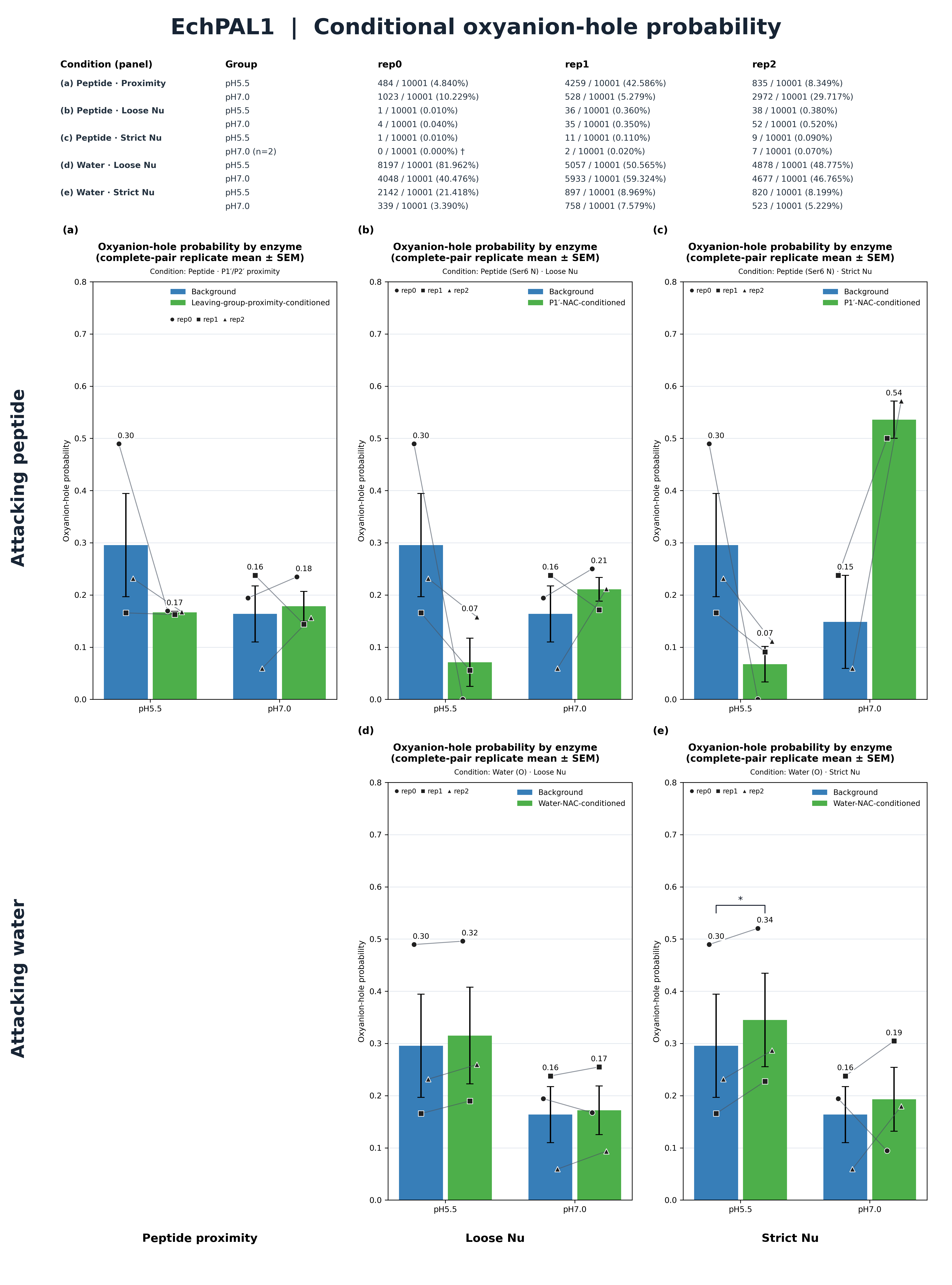

Figure SN12. Comparison of background and conditional oxyanion-hole probabilities in EchPAL1 simulations under different condition definitions. The subtitle gives the number and corresponding fraction of trajectory frames available for conditional-probability calculations in each replicate under each criterion.

Under Strict Nu, peptide-condition frames decreased further to single digits. The three replicates at pH 5.5 decreased from 1/36/38 to 1/11/9 frames, and those at pH 7.0 from 4/35/52 to 0/2/7 (Figure SN12). At pH 7.0, the peptide-conditioned probability under Strict Nu was 3.6 times its background (0.54/0.15), in marked contrast to the trends observed under Loose Nu and Proximity. However, only 0/2/7 qualifying frames were available, severely limiting the reliability of this result.

**Supplementary Note 2**

**2.1 Potential involvement of histidine in reaction directionality**

NAC screening for attacking water used three successively more stringent criteria: Loose Wat, Medium Wat, and Strict Wat. Medium Wat added the oxyanion-hole requirement to Loose Wat, whereas Strict Wat additionally imposed a geometry criterion related to histidine-mediated deprotonation. Because the hierarchical water analysis in the main-text Results suggested a role for the oxyanion hole in reaction directionality, we extended the same approach to the potential involvement of histidine-mediated deprotonation to search for additional mechanistic insights.

For EchAEP1, EchAEP1-G243Q, and EchPAL1, we first quantified the probabilities that attacking water passed the Medium Wat and then Strict Wat criteria and calculated the passing rate for the His-deprotonation-related geometry. At pH 5.5, the rates were 58% (1.43/2.44) for EchAEP1 and 44% (0.46/1.03) for G243Q; at pH 7.0, they were 79% (1.90/2.39) and 9% (0.13/1.31), respectively. For EchPAL1, the rates were 68% (3.53/5.18) at pH 5.5 and 38% (0.46/1.19) at pH 7.0. Thus, the His-deprotonation-related passing rate varied substantially between enzymes and pH conditions (Figures 3e, 4a, and 4e).

In the simulations at both pH 5.5 and pH 7.0, the His-related passing rate was greater for EchAEP1 than for G243Q, qualitatively consistent with the stronger hydrolytic activity of EchAEP1. For EchPAL1, the passing rate was greater at pH 5.5 than at pH 7.0, again consistent with the stronger experimentally observed hydrolysis at pH 5.5. These observations preliminarily suggest a possible relationship between His-mediated proton transfer and hydrolysis efficiency. However, the passing rates only describe occupancy of conformations meeting predefined geometric criteria and cannot be equated with the actual efficiency of His-mediated proton transfer. Additional probability analyses are therefore needed.

Using the Loose Nu criterion, we directly calculated the background and conditional probabilities for catalytic-His deprotonation-related geometry associated with attacking water and incoming peptide in EchAEP1, EchAEP1-G243Q, and EchPAL1. Unlike the background oxyanion-hole probability, the background His-related probability was calculated separately for water and peptide to distinguish the local environment of catalytic His for the two nucleophilic species. The background probability measures how frequently a deprotonation-related His geometry is sampled over the entire trajectory, whereas the conditional probability measures its occurrence among frames satisfying the specified nucleophilic-attack condition. The latter thus evaluates the association of His-related geometry with different pre-attack conformations.

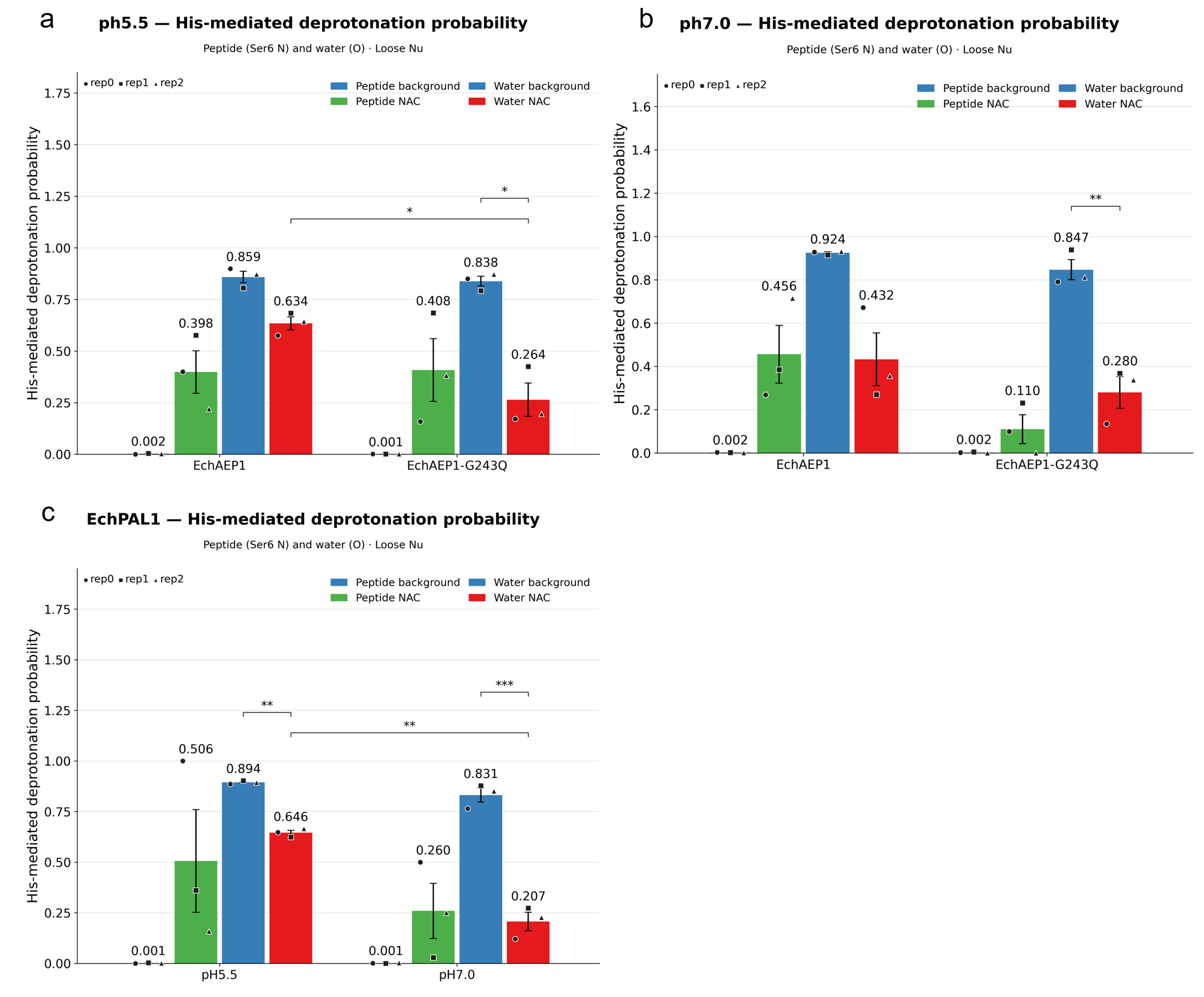

Figure SN13. Background and conditional probabilities of catalytic-His geometry for incoming peptide and attacking water. (a) Analyses at pH 5.5. (b) Analyses at pH 7.0. (c) Analyses of EchPAL1. In each group, from left to right, blue bars represent the peptide-based catalytic-His background probability, green bars the peptide-conditioned catalytic-His probability, blue bars the water-based catalytic-His background probability, and red bars the water-conditioned catalytic-His probability. The nucleophilic-attack criterion was Loose Nu.

The background His-related probability was low when calculated for incoming peptide but high when calculated for water. This difference may reflect the spatial distributions of the two nucleophiles in the simulations: the peptide approached the catalytic center only during some trajectory intervals and remained far from catalytic His for most of the time, whereas water was abundant and could more readily occupy positions near His. The differing background probabilities were therefore sensitive to the spatial distributions of the reacting groups. Conditional probabilities, by contrast, assess His geometry among conformations satisfying a specified nucleophilic-attack criterion and are more appropriate for comparing the relationship between His-deprotonation-related features and competing reaction pathways.

Comparisons of His-related conditional probabilities for incoming peptide and attacking water across the simulated systems showed that both nucleophile-conditioned probabilities were generally higher in the more hydrolytically active systems—EchAEP1 at pH 5.5 and 7.0 and EchPAL1 at pH 5.5—than in the more ligation-favored systems, including G243Q at both pH values and EchPAL1 at pH 7.0. This trend was qualitatively consistent with differences in experimentally observed reaction rates (Figure SN13). Notably, the distinction occurred for both peptide- and water-associated geometries. Thus, the His-deprotonation-related states may not be uniquely associated with attack by one nucleophile but may instead reflect more general features of the local catalytic environment along both pathways, potentially modulating their competition during the second step.

To further compare the relative association of His-deprotonation-related geometries with the two nucleophiles, we calculated the ratio of the peptide-conditioned to water-conditioned His probabilities. At pH 5.5, this ratio was 0.62 (0.398/0.634) for EchAEP1 and 1.54 (0.408/0.264) for G243Q. The higher value for G243Q was consistent with its stronger ligation activity. At pH 7.0, however, the ratios were 1.05 (0.456/0.432) and 0.39 (0.110/0.280), respectively, with G243Q now lower than EchAEP1, contrary to the experimental trend (Figure SN13). This discrepancy may be related to the simplified treatment of His protonation in the simulations: in reality, His can occupy different protonation and tautomeric states that can change with the local environment, whereas all current simulations used fixed HSE, without describing these transitions. For EchPAL1, the ratios at pH 5.5 and pH 7.0 were 0.78 (0.506/0.646) and 1.25 (0.260/0.207), respectively, consistent with stronger ligation at pH 7.0 (Figure SN13).

Taken together, analyses of His-related passing rates and background and conditional probabilities suggest some association between catalytic-His deprotonation-related geometry and reaction directionality across different enzyme backgrounds and pH conditions. Although some results did not agree fully with experimental observations, and the fixed His protonation state in these simulations limits the strength of the conclusions, the partial consistency across multiple analyses suggests that His may contribute to regulation of the second reaction step in legumains. These findings also offer a mechanistic clue to the role of catalytic His in competition between hydrolysis and ligation. Future constant-pH molecular dynamics simulations and more detailed analyses of proton-transfer processes could further examine how dynamic His protonation states affect the competing nucleophilic-attack pathways.
